# Expression of human CEACAMs promotes inflammation and organ damage during systemic *Candida albicans* infection in mice

**DOI:** 10.1101/2025.03.29.646107

**Authors:** Esther Klaile, Mario M. Müller, Johannes Sonnberger, Anne-Katrin Bothe, Saskia Brehme, Juliet Ehrenpfordt, Tilman E. Klassert, Sabina Kuhn, Kristina Dietert, Olivia Kershaw, Jan-Philipp Praetorius, Marc Thilo Figge, Torsten Bauer, Andreas Gebhardt, Gita Mall, Ilse D. Jacobsen, Hortense Slevogt

**Author notes:** **Correspondence:** Esther Klaile.

## Abstract

Invasive candidiasis is a fungal infection characterized by a high mortality rate. CEACAM family receptors play a crucial role in regulating innate responses of both leukocytes and epithelia. Human CEACAM3, CEACAM5 and CEACAM6 receptors recognize *C. albicans* and are expressed in transgenic CEABAC10 mice. In a murine *C. albicans* infection model, CEABAC10 mice exhibited a shortened survival period attributed to an early cytokine storm, an exacerbated acute phase response, and heightened systemic inflammation compared to their wild-type littermates. The livers and kidneys of CEABAC10 mice displayed intensified purulent necrotizing inflammation, accompanied by increased infiltration of neutrophils and macrophages. Our *in vivo* and *in vitro* data indicated that the expression of CEACAM6 on monocytes of CEABAC10 mice caused the elevated cytokine levels and the subsequent exacerbation of the acute phase response upon *C. albicans* infection, resulting in decreased survival.

## 1 Introduction

The fungal pathogen *Candida albicans* is a predominant cause of human *Candida* bloodstream infections, associated with elevated mortality rates^1, 2^. The incidence of systemic candidiasis is on the rise, particularly among patients with high-risk factors such as prior antibiotic exposure, chemotherapy, hematopoietic stem cell transplantation, prolonged intensive care requirements due to other medical conditions, and the use of central venous catheters^3^.

Host defense against systemic candidiasis relies on the crucial role of myeloid phagocytes, including neutrophils, inflammatory monocytes, and tissue-resident macrophages^2^. The initial phase of their interaction with the fungus involves the fundamental step of fungal recognition, which mounts the inflammatory response against the invading fungus, orchestrates pathogen uptake and systemic inflammatory responses during blood stream infections.

In addition to pattern recognition receptors^4, 5^, *C. albicans* is recognized by members of the human CEACAM receptor family, namely CEACAM1, CEACAM3, CEACAM5, and CEACAM6^6^. CEACAM1 and CEACAM3 are modulatory receptors that signal via their intracellular immunoreceptor tyrosine-based inhibitory motif (ITIM) and immunoreceptor tyrosine-based activation motif (ITAM), respectively. CEACAM5 and CEACAM6 are GPI anchored and organized in membrane microdomains^7, 8^. The well-described immunoregulatory function of CEACAM1 is complemented by the impact of CEACAM3, CEACAM5, and CEACAM6 on immune reactions across a diverse array of cell types^9, 10, 11, 12, 13^. *In vitro* data indicate immunoregulatory roles for human CEACAM1 and CEACAM6 in the innate immune response of human intestinal epithelial cells and human neutrophils triggered by *C. albicans* stimulation^6, 14^. However, the expression of human CEACAM1 in transgenic mice had no discernible effect on their susceptibility to systemic candidiasis or on *C. albicans* colonization/dissemination^15^.

Similar to its human homolog, mouse CEACAM1 serves as a receptor for host-specific pathogens such as the murine hepatitis virus (MHV)^16^, and possesses regulatory functions in immune responses across diverse cell types, both *in vivo* and *in vitro*^17, 18^. However, mouse CEACAM1 does not bind to *C. albicans* cell surface structures^6^. Notably, mice lack the homologous genes for CEACAM3, CEACAM5, and CEACAM6 receptors^19^.

Chan and Stanners developed CEABAC10 mice transgenic for human CEACAM3, CEACAM5, CEACAM6, and CEACAM7, including the human promotor region^20^. The extensive epithelial expression of CEACAM5 and CEACAM6 on the surface of various mucosal tissues in CEABAC10 mice, including the intestinal tract, closely mirrors their expression in humans^20^. While CEACAM5 is specifically expressed on apical membranes of epithelial cells, CEACAM6 is also present on immune cells, including neutrophils, alongside the neutrophil-specific CEACAM3^20^. Notably, CEACAM7 does not bind to *C. albicans* and is selectively expressed in a limited number of highly differentiated epithelial cells in the colon^6, 20^. The CEABAC10 mouse model is widely employed for analyzing CEACAM receptor functions in the host’s response to infections with human pathogens. It provided crucial insight into the role of CEACAM6 in conjunction with adhering-invasive *E. coli* (AIEC) in Crohns Disease pathology *in vivo*^21, 22^, and has contributed valuable information regarding the involvement of CEACAM receptors in the neutrophil response to *Neisseria gonorrhea in vitro*^23^.

In the present study, we used the CEABAC10 mouse model to investigate the impact of CEACAM3, CEACAM5, and CEACAM6 on disseminated candidiasis *in vivo* ^15, 24, 25^. The presence of CEACAMs led to severe infection pathology, a heightened IL-6-mediated acute phase response, and early death in candidemic CEABAC10 mice. Through *in vitro* experiments employing primary bone marrow-derived myeloid cells, we found that CEACAM3 expression did not affect inflammatory responses, and we identified CEACAM6^+^ classical monocytes and resident macrophages as the cell populations responsible for the hyperinflammatory response to systemic *C. albicans* infection in CEABAC10 mice. These findings suggest that CEACAM6 may play a previously unknown role in the inflammatory response of mononuclear phagocytes during systemic candidiasis.

## 2 Results

### 2.1 Human CEACAM-transgenic CEABAC10 mice are more susceptible to systemic candidiasis

In an intravenous infection model using two different infectious doses, our results showed a marked increase in susceptibility to systemic candidiasis and earlier death of CEABAC10 mice (CEA) compared to their wild-type (WT) littermates (Figure 1a, Figure S1). Proinflammatory cytokine responses were indistinguishable at 24 h post infection (p.i.) in WT and CEABAC10 mice, but at 72 h p.i., CEABAC10 mice exhibited significantly elevated serum levels of IL-6, TNFα, GM-CSF, IL-2, IL-4, CXCL10, and VEGF, indicative of an exacerbated inflammatory response (Figure 1b-d, Figure S2). Moreover, CEABAC10 mice displayed a diminished count of neutrophils and monocytes in their blood 24 h p.i., suggesting an enhanced recruitment of peripheral leukocytes into infected tissue (Figure 1e, f). Analysis of bone marrow cell populations revealed no disparities in hematopoiesis between CEABAC10 and WT mice during infection (Figure S3). Additionally, CEABAC10 mice exhibited a significant reduction in platelet numbers 24 h p.i., along with enlarged platelet volumes 72 h p.i., suggesting increased consumption and subsequent replenishment (Figure S4), indicative of increased bleeding and thrombosis typical for septic progression^26^.

**Figure 1:**
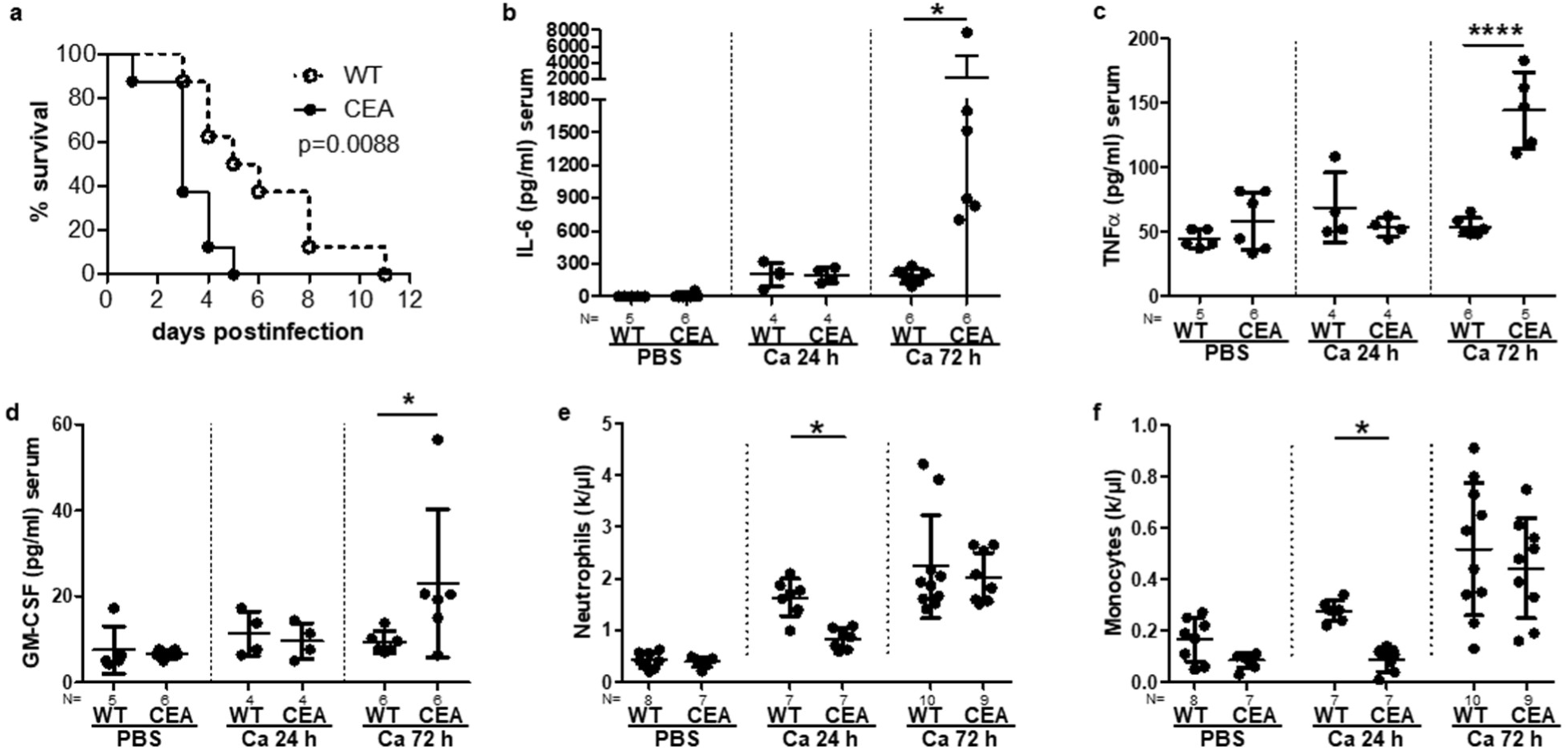
CEABAC10 mice are more susceptible to systemic *C. albicans* infection. (a) CEABAC10 mice (N=8) and wild type littermates (N=8) were injected with 1×10^4^ CFU *C. albicans* /g body weight and were analyzed for survival (one experiment). Note that significant results were also obtained after injection of 2.5×10^4^ CFU/g body weight (Figure S1). (b-f) CEABAC10 mice and wild type littermates were either injected with PBS or infected with 1×10^4^ CFU/g body weight, respectively, and were sacrificed after 24 h or 72 h. Cytokine levels in peripheral blood were determined for IL-6 (b), TNFα (c) and GM-CSF (d) by multiplex assay (Luminex) or ELISA. Additional cytokines are shown in Figure S2. Numbers of neutrophils (e) and monocytes (f) in peripheral blood were analyzed in an automated hemocytometer (additional blood parameters in Figure S4). Statistics: (a) Log rank test (Mantel Cox); (b-f) One-Way-ANOVA and Bonferronis’ Multiple Comparison Test: *p<0.05 ****p<0.001. Data are from one experiment (a) or combined from two independent experiments (b-f). (b-f) Data points with mean and standard deviation.

To explore the inflammatory response and heightened susceptibility of CEABAC10 mice to systemic candidiasis in more detail, we conducted an analysis of serum glycoproteome changes (Figure 2). Principal component analysis revealed that samples from mice injected with PBS separated from those infected with *C. albicans* at 24 h and 72 h, respectively. In addition, CEABAC10 sera notably diverged from WT samples at 72 h p.i. (Figure 2a).

**Figure 2:**
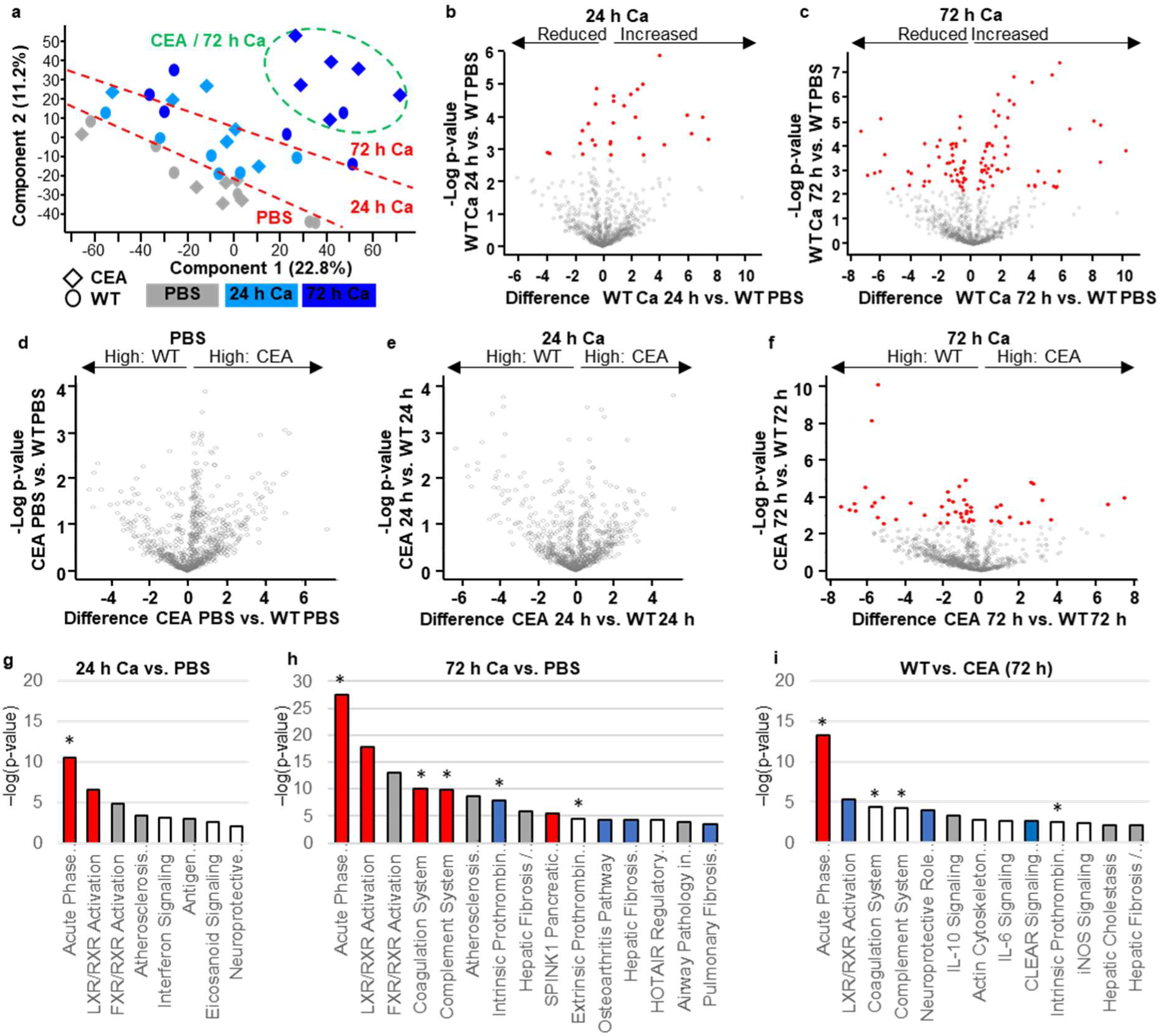
Enhanced acute phase response in CEABAC10 mice after 72 h systemic *C. albicans* infection. CEABAC10 mice and wild type littermates were either injected with PBS or infected with 1×10^4^ CFU/g body weight, respectively, and were sacrificed after 24 h or 72 h (N=6 per group, combined from two independent experiments). Glycoproteins were isolated from serum and analyzed via mass spectrometry. (a) Principal component analysis of identified proteins. Note that PBS injected mice and mice 24 h p.i. and 72 h p.i. separate, respectively, (dashed red lines) but that CEABAC10 samples only separate from WT samples 72 h p.i. (dashed green oval). (b, c) Volcano plots of contrasts of WT samples 24 h p.i. (b) and 72 h p.i. (c) vs. WT PBS, respectively. Plots display –log(p-value) versus the mean difference (log(2) fold-change). Red dots indicate statistically significant changes. All proteins detected and their corresponding values are listed in Tables S1 (b) and S2 (c). (d-f) Volcano plots of contrasts of CEABAC10 vs. WT samples from PBS controls (d), 24 h p.i. (e), or 72 h p.i. (f). Plots display –log(p-value) versus the mean difference (log(2) fold-change). Red dots indicate statistically significant changes. All proteins detected and their corresponding values (f) are given in Table S4. Note that in (d) and (e) no significant alterations were observed. (g-i) Pathway analysis of fold-changes in glycoprotein levels found in (b), (c), and (f), respectively. Asterisks (*) mark extracellular pathways. Bars are color coded for z-scores: red = positive z-score (activated); blue = negative z-score (inhibited); white = z-score equals 0; grey: no activity pattern available. Statistical analysis: MaxQuant, Perseus, and IPA software (settings: materials and methods section).

Consistent with the principal component analysis, the comparison between PBS control-treated and *C. albicans*-infected WT mice revealed 29 and 103 differentially expressed serum glycoproteins at 24 h p.i. and 72 h p.i., respectively (Figures 2b, c and Tables S1, S2). No significant differences in serum glycoprotein abundance were detected between PBS-treated WT and CEABAC10 control mice and between WT and CEABAC10 animals 24 h p.i. (Figure 2d, e). In contrast, at 72 h p.i., the serum glycoprotein composition between WT and CEABAC10 mice revealed significant changes. Overall, 51 glycoproteins were differentially abundant in serum with 14 showing higher abundance in CEABAC10 mice and 37 elevated in WT animals (Figure 2f and Table S5). Pathway analysis with Qiagens IPA software revealed a robust activation of the acute phase response in *C. albicans*-infected WT animals at 24 h p.i. that further progressed at 72 h p.i., accompanied with changes in proteins associated with the activity of the coagulation and complement system and LXR/RXR and FXR/RXR responses (Figures 2g, h and Tables S3, S4). In comparison to WT animals, CEABAC10 mice revealed a stronger acute phase response at 72 h p.i., and alterations of coagulation, complement und prothrombin pathways (Figure 2i, Table S6). Consequently, the serum glycoproteome data suggest an exacerbation of the systemic septic response in CEABAC10 mice initiated by *C. albicans* infection.

### 2.2 Infected kidneys of CEABAC10 mice display pronounced purulent necrotizing nephritis 72 h p.i

To further explore disease progression, we next investigated inflammatory responses in various organs after *C. albicans* infection in more detail. The kidney is the main target organ in the murine intravenous model of systemic candidiasis^27^. Examination of hematoxylin-eosin (HE)-stained kidney sections revealed renal inflammation 24 h p.i. in both WT and CEABAC10 mice (Figure 3a, c, e-h). Subsequently, from 24 h to 72 h p.i., the extent of kidney inflammation escalated in CEABAC10 but remained relatively stable in WT mice (Figure 3b, d, e-h). This was evidenced by the higher inflammation score (Figure 3e), the increased number of inflammatory foci (Figure 3f), and the larger total area of inflamed tissue (Figure 3h). Interestingly, the mean area of inflammatory foci remained similar between WT and CEABAC10 kidneys (Figure 3g). The heightened degree of kidney inflammation in infected CEABAC10 mice was further underscored by significantly higher levels of pro-inflammatory cytokines (IL-6, IL-1β, Figure 3i-j; IL-1α, Figure S5), chemo-attractants (CCL2/MCP-1, Figure 3k; CCL3/MIP1 α, CXCL1/KC, Figure S5), and growth factors (bFGF, GM-CSF, Figure S5) in CEABAC10 kidney homogenates at 72 h p.i.

**Figure 3:**
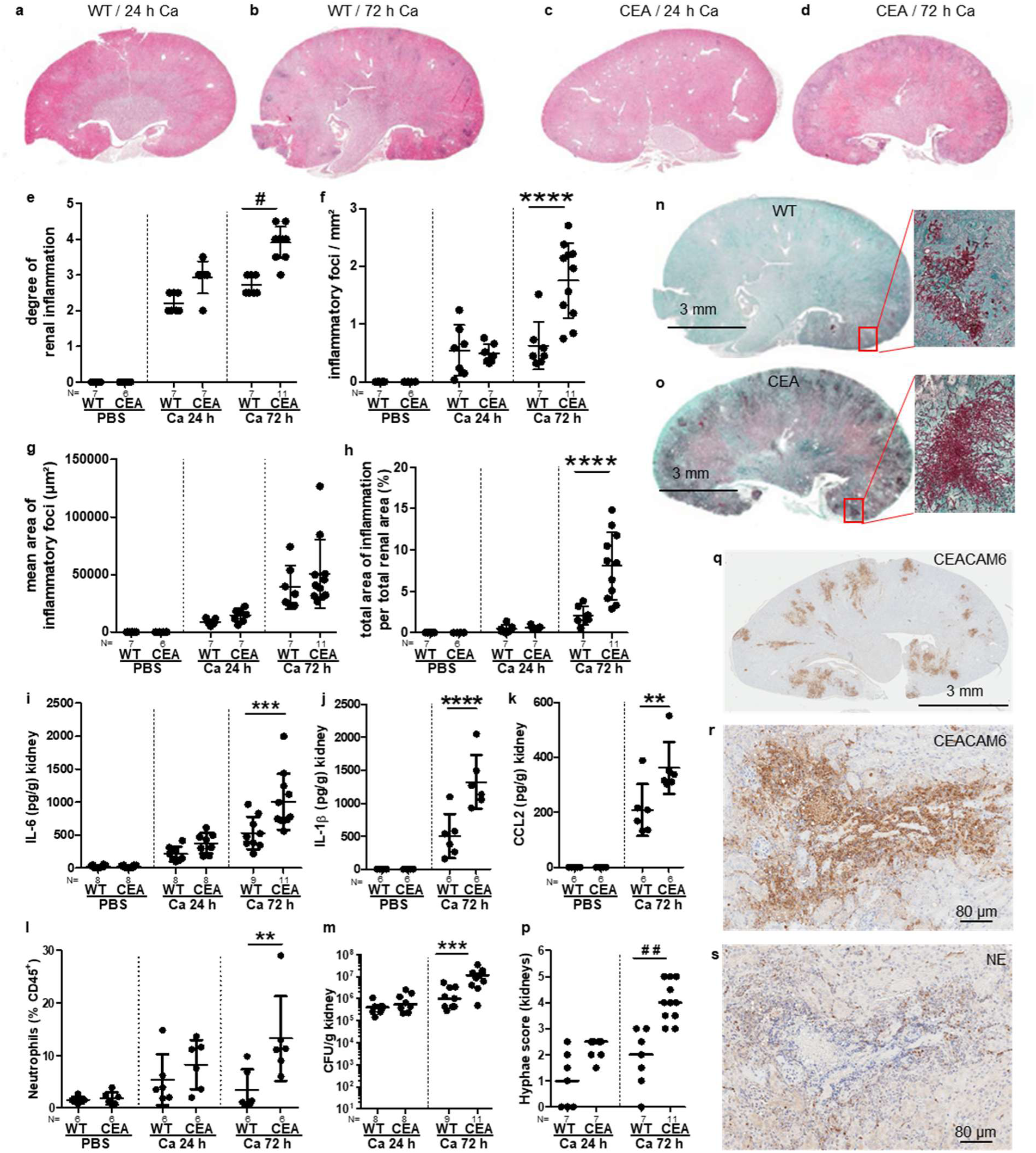
CEABAC10 mice show enhanced kidney inflammation during systemic *C. albicans* infection. CEABAC10 mice and wild type littermates were either injected with PBS or infected with 1×10^4^ CFU/g body weight, respectively, and were sacrificed after 24 h or 72 h. (a-h) Kidney sections were hematoxylin-eosin-stained and analyzed. (a-d) Representative sections of WT and CEABAC10 kidneys. Sections were scored for the degree of renal inflammation (e) and analyzed for inflammatory foci (f, g) and total area of inflammation (h). (i-k, m) Concentrations of IL-6 (i), IL-1β (j) and CCL2/MCP-1 (k), and CFUs (m) were determined in kidney homogenates by multiplex assay (Luminex)/ELISA and serial plating, respectively (additional cytokines in Figure S5). (l) % Ly6G^+^ neutrophils of CD45^+^ leukocytes isolated from kidneys analyzed by flow cytometry (additional immune cell populations in Figure S6, gating in Figure S7). (n, o) Representative Grocott-silver-stained sections 72 h p.i and blow-ups of the indicated regions. (p) Grocott-silver-stained sections were scored for the occurrence of hyphal growth. Note that non-infected kidneys (8 WT and 8 CEABAC10 / PBS) did not show fungal growth (l, o). (q-s) Immunohistochemical staining of consecutive sections of CEABAC10 kidneys 72 h p.i for CEACAM6 (q, r) or neutrophil elastase (s), respectively. Panels display representative images (N=3). Note that only viable neutrophils are NE^+^, but that CEACAM6^+^ cells include viable and dead neutrophils, monocytes, and macrophages, and that CEACAM6^+^ cells (r) outnumber viable neutrophils (s) by ca. one or two orders of magnitude. Statistics: (e, o) Kruskal-Wallis and Dunn’s Multiple Comparison Test, #p<0.05, ##p<0.01; (f-l, p) One-Way-ANOVA and Bonferronis’ Multiple Comparison Test: *p<0.05 **p<0.01 ***p<0.005 ****p<0.001. (e-k) Data points with mean and standard deviation. (m, p) Data points with median and mean, respectively. Data are combined from two independent experiments.

In the kidneys of infected CEABAC10 mice, immune cell populations exhibit a notable relative decrease in NK cells (Figure S6c), but a significant relative increase in neutrophils (Figure 3l) at 72 h p.i.. There was an observed increase in fungal burden, measured as CFU (Figure 3m), visualized by histology (Figure 3n, o), and quantified by automated image analysis of the area covered by fungi (Figure S6f). Also, hyphal growth was increased in CEABAC10 kidneys (Figure 3n, o, p, Figure S6g). Immunohistochemistry (IHC) showed that CEACAM6 was exclusively present in neutrophils, monocytes, and macrophages in CEABAC10 kidney sections (Figure 3q, r). Due to its high glycosylation^28^, CEACAM6 is proteolytically stable, and its presence was detectable in areas with high counts of damaged/deceased immune cells at 72 h p.i., particularly on neutrophils (Figure 3q, r). IHC analysis of consecutive sections for neutrophil elastase (Figure 3s), detectable only in viable neutrophils, indicated that dead cells outnumbered viable neutrophils. Consequently, the quantification of viable neutrophils by flow cytometry likely underestimated the extent of increased neutrophil recruitment into inflamed CEABAC10 kidney tissue.

### 2.3 The expression of human CEACAM3 and CEACAM6 does not alter the response of bone marrow-derived neutrophils to *C. albicans in vitro*

Neutrophils play a pivotal role in the host response against *C. albicans*^3^, yet they also contribute to pathogenesis by inflammation-driven tissue damage^29, 30^. The enhanced recruitment of neutrophils, coupled with a higher fungal burden and increased numbers of inflammatory foci in the kidneys of CEABAC10 mice, implied a potential reduction in the antifungal activity of these cells. Thus, we sought to investigate the response of bone marrow-derived neutrophils (BMN) to *C. albicans* infection *in vitro*. CEABAC10 BMN express CEACAM3 and CEACAM6 on their cell surface (Figure 4a, b), both of which regulate various human neutrophil functions such as pathogen recognition and uptake, modulation of apoptosis, and adhesion to endothelial cells^14, 31, 32, 33^. Given that CEACAM6 is also present in primary/azurophilic granules of human neutrophils and can be de-granulated to increase the cell surface expression level upon neutrophil activation^31^, we assessed the expression of human CEACAM receptors on transgenic BMN by flow cytometry. Upon stimulation of CEABAC10-derived BMNs with *C. albicans*, a significant increase of both, human CEACAM3 and CEACAM6, on BMN cell surfaces was observed (Figure 4c, d), indicating similar behavior of the receptors in transgenic murine neutrophils.

**Figure 4:**
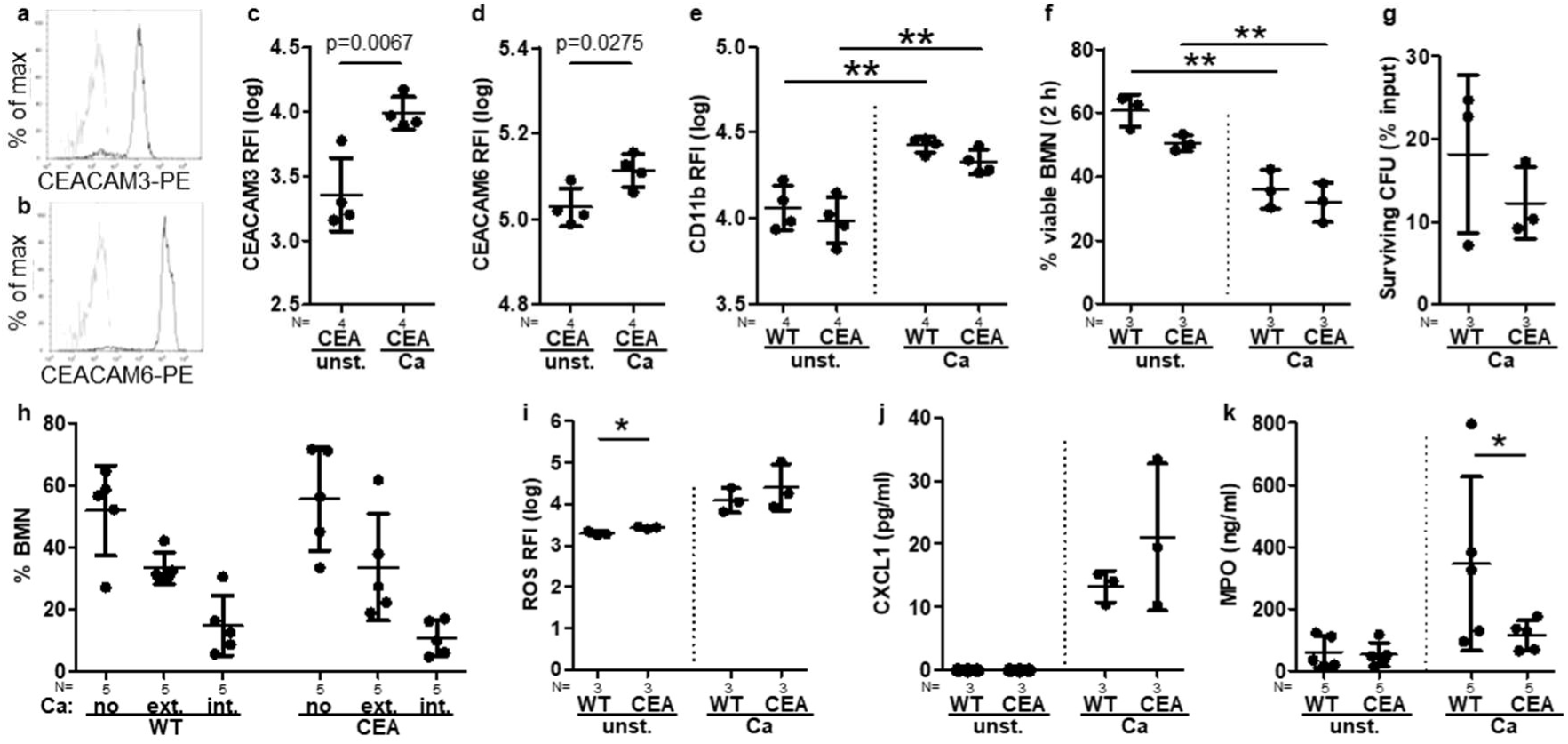
WT and CEABAC10 bone marrow-derived neutrophils display similar reactions to *C. albicans*. (a-e) Degranulation and CEACAM surface expression. WT and CEABAC10 BMN were either left untreated or stimulated with *C. albicans* for 1 h, stained for CD11b and either CEACAM3 (a, c), or CEACAM6 (b, d), respectively, and analyzed by flow cytometry (gating in Figure S8). (a, b) Representative results, grey line: isotype control, black lines: CEACAM3 and CEACAM6, respectively. (c-e) Log data of the relative fluorescence intensity (RFI) with mean and SD. (f) Spontaneous and *C. albicans*-induced cell death after 2 h. Viability was assessed by exclusion of PI and annexin V staining via flow cytometry (% viable BMN with mean and SD). (g) *C. albicans* killing efficiency after 30 min analyzed by XTT assay (% surviving CFUs from the input with mean and SD). (h) *C. albicans* binding/phagocytosis (20 min, FITC-labeled yeast cells at MOI 10, stained for extracellular *C. albicans* via specific antibody) was analyzed by fluorescence microscopy for cells with no contact to *C. albicans* (no), with *C. albicans* bound exclusively extracellular (ext.), and phagocytosed, intracellular *C. albicans* (int.) (% BMN with mean and SD). For each experiment, at least 100 BMN were counted per group. (i) Spontaneous and *C. albicans* - induced production of reactive oxygen species (ROS) after 20 min measured via DHR assay by flow cytometry (log data of RFI with mean and SD. (k, l) Spontaneous and *C. albicans*-induced CXCL1/KC (k) and myeloperoxidase (MPO) (l) release in cell culture supernatants after 24 h with mean and SD. Statistics: (c, d, g) unpaired, two-sided Student’s T test; (e, f, h, i, j, k) One-Way-ANOVA and Bonferronis’ Multiple Comparison Test: *p<0.05 **p<0.01; log data were used for statistical analysis of (c-e, i). (c-k) Data points with mean and standard deviation.

The integrin CD11b/CD18 (CR3, αMβ2, MO-1, Mac-1) plays a crucial role in mediating neutrophil extravasation. CR3 can be activated and upregulated upon human neutrophil activation and ligation of CEACAM3 and CEACAM6^33^. Stimulation of BMNs with *C. albicans* led to a similar increase of cell surface-associated CD11b in both WT and CEABAC10 cells (Figure 4e), indicating similar activation states of BMNs after *C. albicans* recognition. Furthermore, no discernible differences related to spontaneous and *C. albicans*-induced neutrophil death, binding and phagocytosis of *C. albicans*, or fungal killing were observed in isolated BMNs from WT and CEABAC10 mice *in vitro* (Figure 4f-h). While a slightly higher basal level of reactive oxygen species (ROS) was detected in CEABAC10 BMNs, the quantities of ROS induced by *C. albicans* were comparable between WT and CEABAC10 BMNs (Figure 4i). Also, there was no significant difference in the release of proinflammatory CXCL1 (KC), but CEABAC10 BMNs exhibited a less variable and lower release of the degranulation marker myeloperoxidase (Figure 4j, k).

In summary, the expression of human CEACAM3 and CEACAM6 did not fundamentally alter the interaction of BMNs with *C. albicans in vitro*. However, the microenvironment of inflamed tissues may well influence neutrophil behavior *in vivo*, and the elevated numbers of neutrophils likely contribute to heightened inflammation and tissue damage.

### 2.4 CEABAC10 mice develop acute hepatic coagulation necroses, purulent necrotizing hepatitis, purulent splenitis, and brain hemorrhage during systemic candidiasis

Following intravenous infection of WT mice, *C. albicans* is typically cleared from the liver over time without causing visible tissue alterations^27^, a trend also observed in our experiments (Figure 5a, c). In contrast, at necropsy 72 h p.i., prominent white areas were evident on the livers of 15 out of 18 CEABAC10 (Figure 5b, c). Histologically, these alterations correlated with multifocal acute coagulation necroses with immune cell infiltrations (Figure 5d-f). Notably, these lesions were strictly vascular associated, predominantly periportally localized, and nearly exclusive to CEABAC10 livers (Figure 5g). The onset of coagulation necroses and immune cell infiltrations was detected microscopically as early as 24 h p.i., intensifying in numbers and size by 72 h p.i. (Figure 5g), when they became macroscopically visible. Additionally, acute purulent necrotizing hepatitis was observed in 6 out of 11 CEABAC10 mice 72 h p.i., contrasting with the absence of such findings in WT animals (Figure 5h).

**Figure 5:**
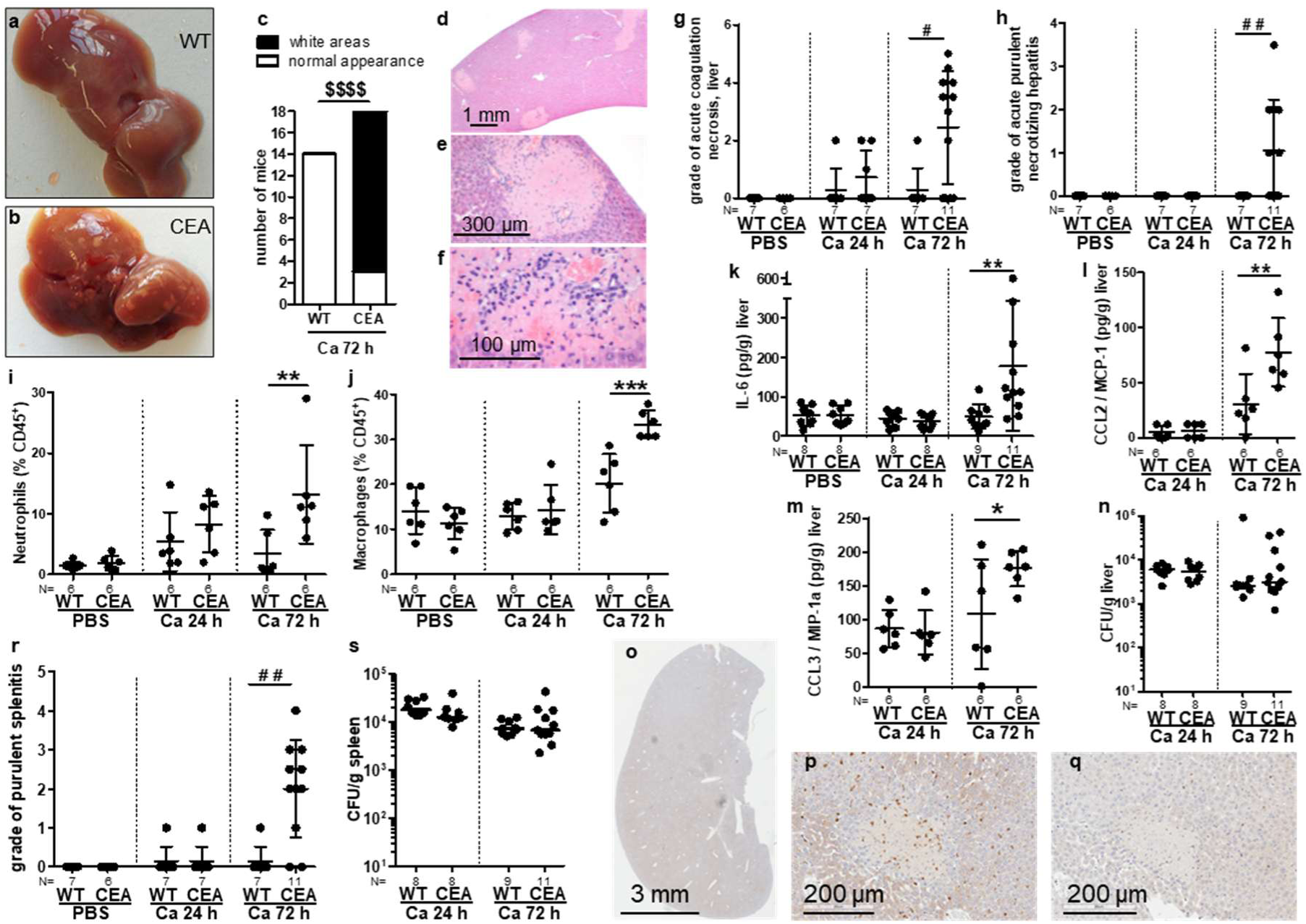
CEABAC10 livers display enhanced inflammation, acute coagulation necroses with immune cell infiltration, and multifocal hemorrhage during systemic *C. albicans* infection. CEABAC10 mice and wild type littermates were either injected with PBS or infected with 1×10^4^ CFU/g body weight, respectively, and were sacrificed after 24 h or 72 h. (a-c) Occurrence of macroscopic pathologic liver abnormalities (white areas) observed during necropsy 72 h p.i. (18 CEA, 14 WT). (a, b) Representative images. (c) Livers without macroscopic abnormalities (open bars), and livers displaying white areas 72 h p.i. (filled bars). (d-h) Liver sections were hematoxylin-eosin-stained and analyzed. (d-f) Representative CEABAC10 liver section 72 h p.i., showing acute coagulation necroses with immune cell infiltration and multifocal hemorrhage. (g, h) Sections were scored for the degree of acute coagulation necroses (g) and for the grade of acute purulent necrotizing hepatitis (h). (i, j) % Ly6G^+^ neutrophils (i), and F4/80^+^ macrophages (j) of CD45^+^ leukocytes isolated from livers (additional immune cell populations in Figure S10, gating in Figure S11). (k-m) Concentrations of IL-6 (k), CCL2/MCP-1 (l) and CCL3/MIP-1alpha (m) were determined in liver homogenates (additional cytokines in Figure S9). (n) CFUs in liver homogenates. (o-q) Immunohistochemical staining of consecutive CEABAC10 liver sections 72 h p.i. for CEACAM6 (o, p) or neutrophil elastase (NE) (q), respectively (representative sections, N=3). Note that only viable neutrophils are NE^+^, but that CEACAM6^+^ cells include viable and dead neutrophils and macrophages/monocytes, and that CEACAM6^+^ cells (p) outnumber viable neutrophils (q) by one or two orders of magnitude. (r) Spleen sections were hematoxylin-eosin-stained and scored for splenitis. (s) CFUs were determined in spleen homogenates. Note that livers and spleens from uninfected animals (8 WT and 8 CEABAC10 for PBS, respectively) did not display any fungal growth and that none of the infected liver and spleen samples displayed any hyphal growth. Data are combined from two (g-n, r, s) or four (c) independent experiments. Statistics: (c) Fisher’s Exact Test, two-sided $$$$ p<0.001; (g, h, r) Kruskal-Wallis and Dunn’s Multiple Comparison Test, #p<0.05 ##p<0.01; (i-n, s) One-Way-ANOVA and Bonferronis’ Multiple Comparison Test: *p<0.05 **p<0.01. For (l, s), log data were used for statistical analysis. (g-n, r, s) Data points with mean and standard deviation.

The inflammatory nature of these lesions was evident through the higher relative numbers of neutrophils and macrophages detected by flow cytometry in livers of CEABAC10 mice (Figure 5i, j), accompanied by elevated levels of pro-inflammatory cytokines and chemokines in liver homogenates (IL-6, CCL2/MCP-1, CCL3/MIP-1α, and CXCL10/IP-10; Figure 5k-m, Figure S9). While the proportion of monocytes among hepatic immune cells was comparable between WT and CEABAC10 mice, there was a tendency towards a relative reduction of NK cell, T cell, and B cell numbers in CEABAC10 livers 72 h p.i. (Figure S10). Despite the heightened inflammation in the livers of CEABAC10 mice, the total fungal burden in the liver tissue was comparable to WT animals (Figure 5n). Similar to the kidneys, only neutrophils, along with macrophages and monocytes stained positive for CEACAM6 (Figure S10), and live, neutrophil elastase (NE)-positive granulocytes were outnumbered by dead CEACAM6-posive neutrophils and macrophages 72 h p.i. (Figure S10).

In addition to the liver pathology, severe purulent splenitis was also observed in 8 out of 11 CEABAC10 mice at 72 h p.i., a condition only detected in 1 out of 7 WT mice (Figure 5r). Despite this discrepancy in pathology, the composition of immune cell populations and fungal load were not significantly different in spleens of infected WT and CEABAC10 mice (Figure S12, Figure 5s).

Interestingly, four out of 18 CEABAC10 mice exhibited brain hemorrhage at 72 h p.i. at necropsy, which was not seen in any of the 14 WT mice at 72 h (Figure S14). No difference in the severity of encephalitis was observed between the two genotypes, although CEABAC10 brains displayed higher fungal burdens and higher hyphae scores (Figure S14). Similar to other organs, CEACAM6 expression in the brain was restricted to neutrophils and some macrophages (Figure S14), and dead CEACAM6-positive neutrophils and macrophages outnumbered viable, neutrophil elastase (NE)-positive granulocytes (Figure S14).

### 2.5 CEACAM6 expression on monocytes, but not macrophages, leads to increased cytokine production in response to *C. albicans*

The activation of leukocytes induces an increased expression of CEACAMs on their surface^31^. In evaluating the activation of myeloid cells following systemic candidiasis, we focused on assessing the expression of CEACAM6 that is expressed by neutrophils, monocytes, and macrophages in CEABAC10 mice. The proportion of CEACAM6^+^ neutrophils increased from approximately 80% to nearly 100% at 72 h p.i. in kidney, liver, and spleen (Figure 6a, d, g). While CEACAM6^+^ monocytes were scarce in PBS control mice, their numbers increased to approximately 50% of the total monocyte population in the kidney, liver, and spleen at 72 h p.i. (Figure 6b, e, h).

**Figure 6:**
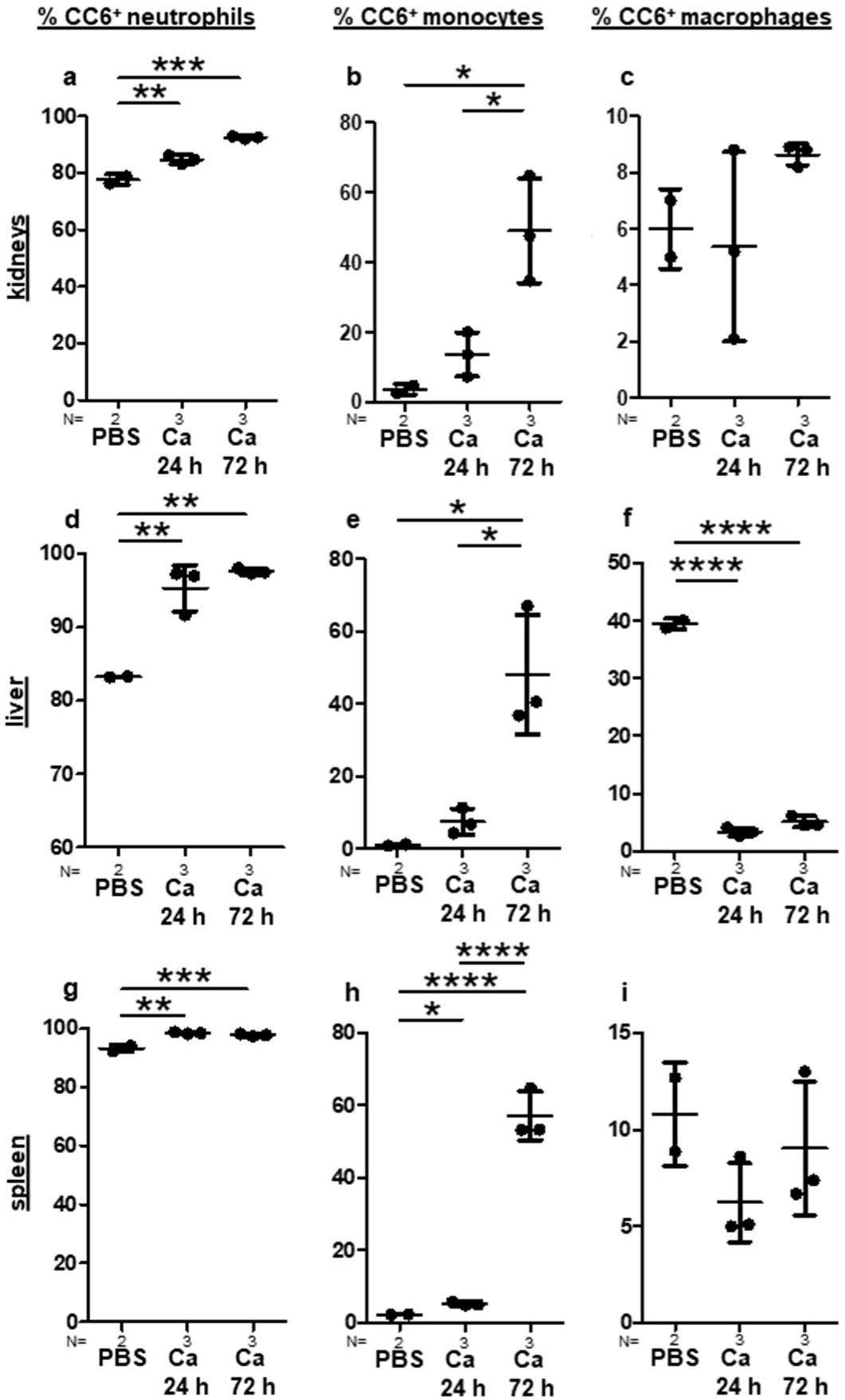
Increased numbers of CEACAM6^+^ myeloid cells in organs of CEABAC10 mice during systemic *C. albicans* infection, and loss of CEACAM6^+^ liver macrophages. CEABAC10 mice were either injected with PBS or infected with 1×10^4^ CFU/g body weight, respectively, and were sacrificed after 24 h or 72 h. Immune cells were isolated from kidneys, liver, and spleen, and stained for viability dye/CD45/CD11b/Ly6G, and viability dye/CD45/F4/80/Ly6C/CD11c, respectively. Neutrophils, monocytes, and macrophages were analyzed for their percentage of human CEACAM6-positive cells (gating: see Figure S15). Graphs give the percentage of CEACAM6-positive cells of the respective cell type with mean and standard deviation. Data are from one experiment. Statistics: One-Way-ANOVA and Bonferronis’ Multiple Comparison Test: *p<0.05 **p<0.01 ***p<0.005 ****p<0.001.

In macrophages, the CEACAM6^+^ subpopulation remained remarkably stable during infection in kidneys and spleen. Approximately 6% of all macrophages in the kidneys (Figure 6c) and around 10% in the spleen (Figure 6i) were CEACAM6-positive cells. Interestingly, a contrasting scenario unfolded in the liver: Here, a substantial proportion of CEACAM6^+^ macrophages (approximately 40%) was observed in PBS-treated control animals (Figure 6f). However, this population dropped below 10% within 24 h p.i. and remained low at 72 h p.i. (Figure 6f). One plausible explanation for the reduced proportion of CEACAM6^+^ macrophages is macrophage cell death, as only viable cells were quantified by flow cytometry, and a substantial number of CEACAM6^+^ dead cells, including macrophages, were visible in immunohistochemistry (Figure 5p).

Since the expression of CEACAM3 and CEACAM6 did not influence BMN interaction with and response to *C. albicans*, we postulated that CEACAM6^+^ transgenic macrophages and monocytes are crucial for the heightened inflammatory response observed in CEABAC10 mice during systemic candidiasis. We therefore studied responses to *C. albicans* infection in transgenic and WT bone marrow-derived macrophages (BMDMs) *in vitro*. The majority of BMDMs stained positive for CEACAM6 (Figure 7a, b). While phagocytosis of *C. albicans* was slightly but significantly higher in CEABAC10 BMDMs (Figure 7c), *C. albicans*-induced macrophage cell death was comparable between both WT and CEABAC10 BMDM (Figure 7d). These findings suggest that macrophage expression of CEACAM6 *per se* is not the primary factor responsible for the substantial number of dead hepatic macrophages observed after infection *in vivo* (Figure 6f). Moreover, the release of the pro-inflammatory cytokines IL-1β, IL-6, and TNFα was not significantly affected by CEACAM6 expression in BMDMs (Figure 7d-f).

**Figure 7:**
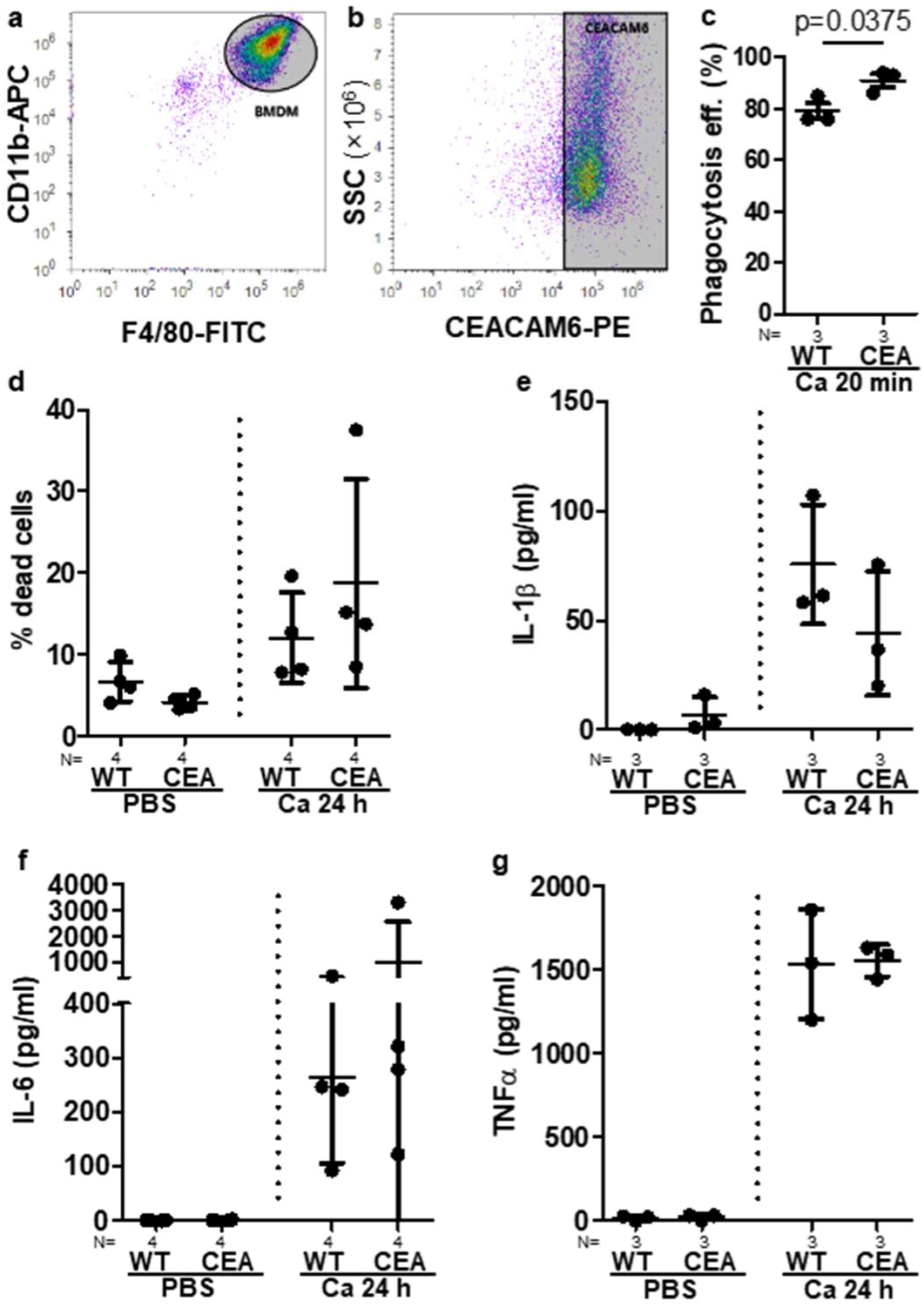
Enhanced phagocytosis but unaltered *C. albicans*-induced cell death of CEACAM6^+^ bone marrow-derived macrophages. (a, b) Bone marrow-derived macrophages (BMDM) from CEABAC10 mice were analyzed via flow cytometry. Single viable cells were gated for F4/80^+^/CD11b^+^ macrophages (a) and analyzed for their CEACAM6 expression (b); for complete gating see Figure S16. Graphs are representative of three independent experiments (mean and standard deviation: 88.9%±5.1%CEACAM6^+^ BMDM). (c) WT and CEABAC10 BMDM were incubated with FITC-labeled *C. albicans* at MOI 5 for 20 min, fixed, and stained for extracellular *C. albicans*. Images were analyzed for *C. albicans* bound externally to BMDM, and phagocytosed/intracellular *C. albicans*. The graph gives the phagocytosis efficiency in %: intracellular *C. albicans* / (externally bound + intracellular *C. albicans*) × 100. N=3, at least 100 cells were analyzed per sample. (d) WT and CEABAC10 BMDM were either left unstimulated or incubated with *C. albicans* at MOI 1 for 24 h and analyzed for the percentage of dead cells by Sytox green staining; at least 100 cells were analyzed per sample. (e-g) WT and CEABAC10 BMDM were either left unstimulated or incubated with *C. albicans* at MOI 1 for 24 h. Concentrations of IL-1β (e), IL-6 (f), and TNFα (g), were determined in cell culture supernatants via ELISA. Graphs give data points with mean and SD. Statistics: (c) unpaired two-sided Student’s T test; (d-g) One-Way-ANOVA and Bonferronis’ Multiple Comparison Test. (c-g) Data points with mean and standard deviation.

In CEABAC10 bone marrow-derived monocytes (BMMs), only the classical subset (CD62L^+^/CD11c^-^), and not the non-classical subset (CD62L^-^/CD11c^+^) expressed CEACAM6 on the surface (Figure 8a-d). Nevertheless, this mixed population of CEABAC10-derived BMMs with >50% CEACAM6^+^ cells demonstrated a significantly increased production of IL-6 and CCL2/MIP-1α after *C. albicans* infection compared to WT BMMs, aligning with the higher cytokine production observed in mouse tissues *in vivo*. In contrast, the amount of secreted TNFα and IL-1β after infection was comparable in both WT-and CEABAC10-derived BMMs irrespective of CEACAM6 expression.

**Figure 8:**
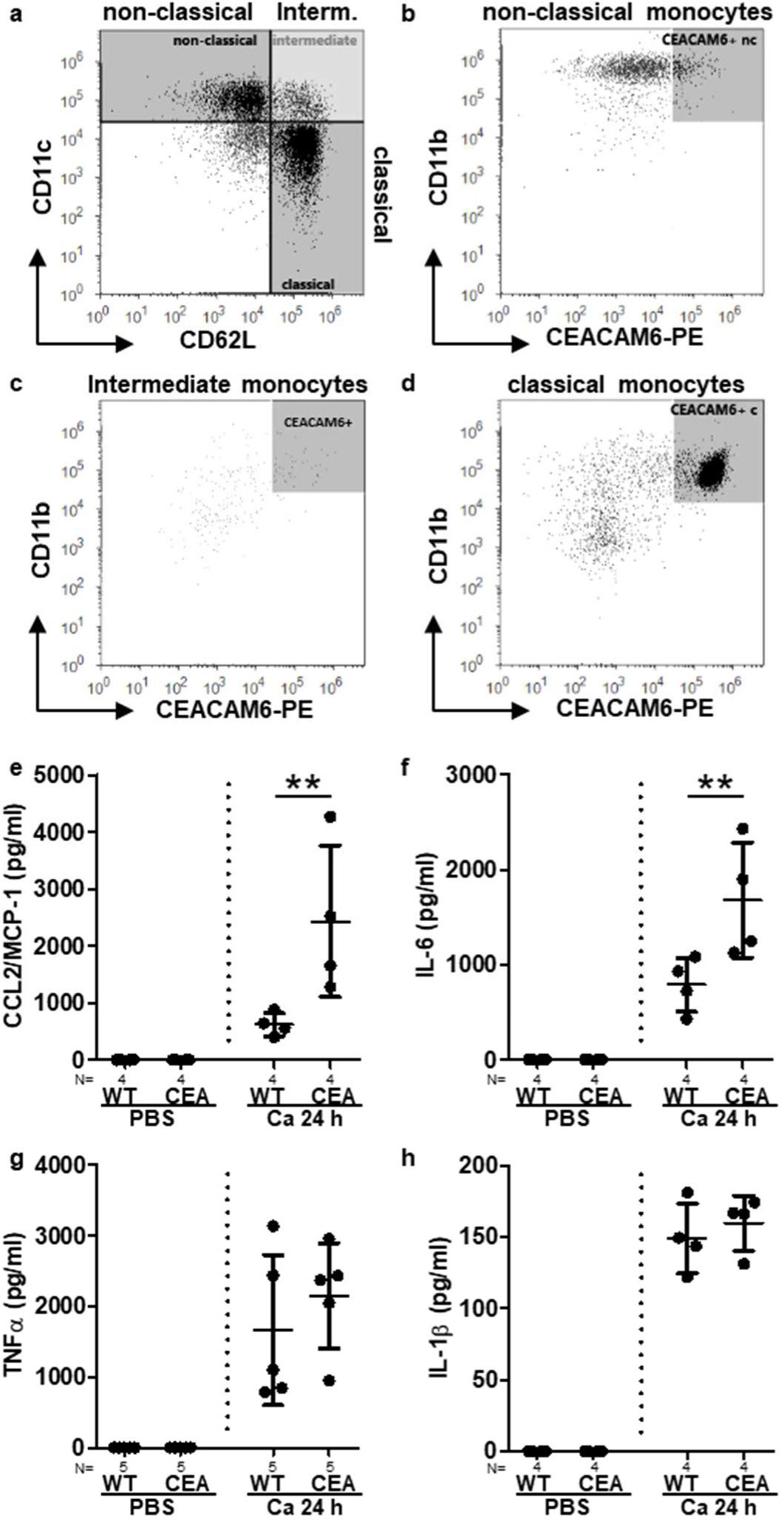
Bone marrow-derived CEACAM6^+^ classical monocytes release enhanced amounts of IL-6 and CCL2 in response to *C. albicans* infection. (a-d) Bone marrow-derived monocytes (BMM) from CEABAC10 mice were analyzed by flow cytometry. Single viable BMM were gated into CD11c^+^/CD62L^-^ non-classical monocytes, CD11c^+^/CD62L^+^ intermediate monocytes, and CD11c^-^/CD62L^+^ classical monocytes (a), and analyzed for their CEACAM6 and CD11b expression (b-d, respectively). For complete gating see Figure S16. Graphs are representative of three independent experiments (mean and standard deviation: CEACAM6^+^ non-classical monocytes = 1,6%±0.4%; CEACAM6^+^ intermediate monocytes = 36.9%±3.8%; CEACAM6^+^ classical monocytes = 83.4%±6.4%). (e-h) WT and CEABAC10 BMM were either left unstimulated or incubated with *C. albicans* at MOI 1 for 24 h. Concentrations of CCL2/MCP-1 (e), IL-6 (f), TNFα (g), and IL-1β (h), were determined in cell culture supernatants. Graphs give data points with mean and standard deviation. Statistics: One-Way-ANOVA and Bonferronis’ Multiple Comparison Test, **p<0.01.

The identification of a specific subset of macrophages expressing CEACAM6 in the livers of transgenic mice aligns with the results from the analysis of human liver sections, where a substantial portion of resident CD68^+^ macrophages^34^ was also positive for CEACAM6 (Figure S17). Additionally, closely mirroring this pattern, about one-third of CD169^+^ alveolar macrophages^35^, as detected via flow cytometry in cell pellets from human bronchoalveolar lavage samples, were CEACAM6-positive (Figure S18). In contrast to CEABAC10 monocytes (Figure 6b, 3, h), peripheral monocytes isolated from fresh blood of healthy human volunteers showed no detectable surface expression of CEACAM6 (Figure S19).

## 3 Discussion

CEACAM family receptors play a crucial role in regulating immune functions during infections and cancer^8, 28, 36^. CEACAM1 is recruited and activated by bacterial and viral pathogens to suppress immune responses^8, 28, 36^. Conversely, CEACAM6 primarily functions as an adhesin for human pathogens like AIEC, *Neisseria spec*., and *Acinetobacter baumannii*, permitting colonization and invasion of cells and tissues without directly affecting host responses^10, 13, 21^. CEABAC10 mice^20^ serve as a valuable model for studying interactions of human pathogens with CEACAM3, CEACAM5, and CEACAM6 both *in vivo* and *in vitro*^13, 21, 22, 23, 37, 38^. In this study, we explore the impact of their expression on the host response to systemic candidiasis.

CEABAC10 mice exhibited a heightened susceptibility and accelerated mortality in response to systemic *C. albicans* infection. This increased vulnerability can be at least partially attributed to an exacerbated systemic immune response, characterized by elevated cytokine levels and an intensified acute phase response, in line with the contribution of progressive sepsis to death of mice systemically infected with *C. albicans*^39^. In addition, increasing fungal burden in the kidneys as the primary target organ^27^ leads to renal failure^39^. Here, we found that CEABAC10 kidneys were severely affected at 72 h p.i. with increased inflammation and fungal burden, indicating that both augmented immunopathology and reduced ability to control fungal growth contributed to the increased susceptibility of transgenic mice.

Neutrophils are crucial for the host defense against systemic candidiasis^3^, and they express functional CEACAM3^23^ and CEACAM6^8, 28^. Thus, functional impairment of transgenic neutrophils would explain the higher fungal burden, and increased recruitment of neutrophils in CEABAC10 mice in response to the high fungal load likely resulted in increased renal injury^40, 41^. However, we found no evidence of reduced anti-fungal activity of CEABAC10 BMNs *in vitro*. Possibly, neutrophil behavior *in vivo* is influenced by infection-specific alterations of the tissue environment that were absent in BMN experiments *in vitro*. Our serum glycoproteome analysis showed markedly dysregulated levels of several matrix proteins and secreted/shed immunoregulatory receptors and enzymes in CEABAC10 serum, including the key enzyme for heparan sulfate chain initiation, EXTL2. In mouse models for vulvovaginal candidiasis, heparan sulfate in the vaginal environment of susceptible mice serves as a competitive ligand for Mac-1 on neutrophils, effectively rendering the neutrophils unable to bind to *Candida* to initiate killing^42^. Pathway analysis of the serum glycoproteomes of WT and CEABAC10 mice 72 h p.i. further identified a dysregulation of the LXR/RXR activation, which regulates neutrophil functions^43^ as well as monocyte/macrophage cytokine release^44, 45^.

Interestingly, CEABAC10 livers also showed clear inflammatory changes and tissue damage. This was unexpected because in WT mice, liver infection is normally controlled by the murine immune system, and tissue architecture is preserved without evident inflammatory changes, except for the transient accumulation of both neutrophils and mononuclear phagocytes^27^. Enhanced numbers of monocytic phagocytes can lead to tissue damage, as shown by mouse liver injury exacerbated by CCL2-mediated monocyte/macrophage recruitment^46^. In fact, our data suggested that CEABAC10 monocytes and macrophages were the driving force behind the observed systemic inflammation and organ damage, acutely affecting the liver. Recently, an increasing number of publications have focused on the role of monocytic phagocytes in systemic candidiasis^3, 47^. Monocytes closely control the early host reaction to *C. albicans* infection, primarily by secretion of various cytokines and chemokines, including IL6 and CCL2/MIP-1α^48, 49, 50^. *Candida*-stimulated CEABAC10 bone marrow-derived monocytes released substantially increased amounts of IL-6 and CCL2/MIP-1α, consistent with their marked increase *in vivo* in infected CEABAC10 mice.

As the primary immune cells in naive tissues, tissue-resident macrophages are essential for innate host defense against *C. albicans* infection^47^. CCL2 further enhances monocyte recruitment to inflammatory sites, where they differentiate into macrophages^47, 51^, as we found in CEABAC10 livers 72 h p.i. CEABAC10 liver macrophages showed a contrasting behavior with the loss of all CEACAM6^+^ macrophages within 24 h p.i. and a strongly increased total number of CEACAM6^-^ macrophages. Regrettably, bone marrow-derived CEABAC10 macrophages were not a useful tool to study the role of macrophages during *Candida* infection, probably due to the marked heterogeneity of macrophage populations as a function of their microenvironment, as reviewed in detail previously^3, 47, 52^. However, in a CX3CR1 knock-out mouse model, the dysfunction of tissue macrophages was associated with a higher fungal load in kidneys, higher neutrophil counts in kidneys, renal failure, and shortened survival^53^. As in most murine studies of systemic *C. albicans* infection, the liver was not further analyzed in this study^53^ as it is usually not severely affected^27^.

In CEABAC10 mice, the liver as key integrator of microbial responses^54^ was likely central to the exacerbation of sepsis progression via enhanced activation of the acute phase response elicited by the increased levels of monocyte-derived IL-6. During microbial infection, IL-6 induces the liver to switch from tolerogenic towards immunogenic responses and initiate the production of acute-phase proteins^54, 55^. In mice as well as in human patients, serum levels of IL-6 are closely related to the severity and outcome of sepsis^56, 57^. In fact, IL-6 is the major factor inducing a systemic inflammatory response syndrome in mouse models of sepsis^58^, furthered by inflammation and coagulation that promote organ dysfunction^26^.

The rapid septic progression in CEABAC10 mice driven by the liver may have obscured further roles of CEACAM6 receptors on monocytes and tissue macrophages in other organs^59^. For example, CEABAC10 brains showed increased fungal burden and hemorrhage. *C. albicans* enters the brain by damaging the blood-brain-barrier^60^, and microglia promote the recruitment of neutrophils to the brain, which mediate fungal clearance^60, 61^. Further studies will be needed to address the possible role of CEACAM6 expression on microglia and other immune cells that may further impair blood vessel permeability.

This study confirms the results of our previous research on human neutrophils^14^, which showed that CEACAM6 expression plays a role in enhancing the responses triggered by other receptors upon *C. albicans* infection rather than directly initiating a response. While CEACAM1 shows very similar behavior to CEACAM6 in response to *C. albicans* infection on human neutrophils^14^, mice transgenic for human CEACAM1 show no difference in their response to systemic candidiasis compared to their WT littermates^15^. Our data show that in human neutrophils, CEACAM6 can influence different pattern recognition receptor pathways, including Toll-like receptor (TLR), NOD-like receptor, and c-type lectin receptor signaling.

However, results from TLR2- and TLR-4-deficient mice infected with *C. albicans* remain controversial, and in the consequences of TLR knock out for survival and fungal burden depend on the genetic background, fungal strain, and infection dose^62^. Therefore, it remains unclear if TLR interference by CEACAM6 might contribute to the increased susceptibility of CEABAC10 mice. Dectin-1 deficiency renders mice susceptible to *C. albicans* infection due to impaired neutrophil, monocyte, and macrophage recruitment and responses, including impaired cytokine release, after infection even in the presence of opsonins^63^. This is in stark contrast to increased recruitment of leucocytes and immunopathology observed with CEABAC10 mice, making it unlikely that interference of CEACAM6 with Dectin-1 signaling contributed to the observed phenotype.

Because of the intravenous route of infection used in this study, we believe that CEACAM6 expression on myeloid cells, particularly monocytes and monocyte-derived cells, rather than mucosal CEACAM expression, lead to the exacerbated systemic inflammatory response syndrome found in CEABAC10 mice. While there are some early publications indicating the expression of CEACAM6 (also named CD66c, NCA-50/90, non-specific cross-reacting antigen) on human monocytic cells and tissue macrophages^64, 65, 66^, we were only able to verify the expression of CEACAM6 in human tissue macrophages. The role of CEACAM6^+^ mononuclear phagocytes remains to be further elucidated in primary human cells. Also, the possibility of CEACAM6^+^ monocyte subpopulations in systemically infected or septic patients should be further explored.

## 4 Materials and Methods

### 4.1 *Candida albicans* strain and culture

*Candida albicans* Berkhout strain SC5314 was used for all experiments and was grown as described^15^. In some cases, germ tubes were induced as described^15^.

For FITC labelling, 10 × 10^7^ yeast cells were suspended in 10 ml carbonate buffer (pH 9.5; 70% of sodium bicarbonate, 30% of sodium carbonate) and 100 µl of a FITC stock solution (10 mg/ml in PBS) and incubated for one hour at 23°C, 150 rpm in the dark. Labeled yeast cells were washed twice with PBS.

### 4.2 Mouse strains

FVB mice transgenic for the human CEACAM3, 5, 6, and 7 gene, CEABAC10^20^, were crossed into the C57BL/6NRj background using Speed Congenics. The status of the background was supervised by GVG Genetic Monitoring (Leipzig, Germany). Mice were bred heterozygous and a minimum of 12 backcrosses were performed prior to the first experiment. The genotypes (CEABAC10+/- or wild type) were determined by PCR analysis of tail biopsies using the following primer pairs for the transgenic human CEACAMs (5’–3’): CEACAM3 (AACCCCAGGACAGCAGCTTC and GAGAGGCCTTTGTCCTGACC), CEACAM5 (CATTTGCAACAGCTACAGTC and AGTGCAGTGGTATCAGAAAC) CEACAM6 (TACTCAGCGTCAAAAGGAAC and AGAGACGTGGATCATCATCGTGA), CEACAM7 (TGATCCTCCTGATTGTCACA CTACTGGGCAATACAACAGT). Mouse interferon beta primers ATAAGCAGCTCCAGCTCCAA and GCAACCACCACTCATTCTGA were used as positive control. Wild type littermates were co-housed and used as controls. Mice were maintained under specific pathogen-free conditions at the animal facility *Forschungszentrum Beutenberg, Zentrale Experimentelle Tierhaltung*, University Hospital Jena, Germany, according to European and German animal welfare regulations.

### 4.3 Ethics statement

Animal studies were performed in strict accordance with European (The Council of Europe’s European Convention, March 18, 1986, with the revised Annex A 2010/63/EU; The European Parliament and Council Directive 2010/63/EU d 22.09.2010) and German animal welfare regulations and the recommendations of the Society for Laboratory Animal Science (GV-Solas). All experiments were approved by the ethics committee “Beratende Kommission nach §15 Abs. 1 Tierschutzgesetz” and the responsible Federal State authority *Thüringer Landesamt für Verbraucherschutz*, Bad Langensalza, Germany (Permit No. 02-019/14).

Studies with human materials were conducted according to the principles expressed in the Declaration of Helsinki. Written informed consent to participate in this study or to allow the use of residual material for research purposes was obtained. Protocols and use of cells for this study were reviewed and approved by the institutional ethics committee of the Ethics Committee of the Friedrich Schiller University Jena, Medical Faculty (permission numbers 5070-02/17 and 2020–1773).

### 4.4 Systemic *C. albicans* infection model

Systemic infection was performed as described previously^15^. Briefly, co-housed young adult CEABAC10-transgenic mice (CEA) and their wild type littermates (WT) were injected with 2.5 × 10^4^ CFU *C. albicans* /g body weight or 1 × 10^4^ CFU *C. albicans*/g body weight via the lateral tail vein. After the infection, mice were scored at least twice a day and sacrificed when reaching a humane endpoint as described^15^. Log Rank comparison of the survival curves was performed with the GraphPad PRISM 5 software.

For analyses at pre-defined time points after infection, mice were infected as described above with 1 × 10^4^ CFU *C. albicans* /g body weight (“Ca”). Control mice received endotoxin-free buffered saline solution (“PBS”, InVivoPure pH 6.5 Dilution Buffer, Hoelzel Diagnostika GmbH, Germany; vehicle control). After infection, the health status of the mice was examined as described^15^, and mice were sacrificed after 24 h (PBS and Ca) or 72 h (Ca). Since the vehicle control, PBS, does not result in any immune reaction^15^, and any reaction due to the stress of the application would be only short-lived, no additional PBS group was analyzed at 72 h. Note that one PBS-injected CEABAC10 mouse was removed from all analyses after histological analysis identified a high-grade hydronephrosis with associated, presumably ascending pyelonephritis that was not associated with the experiment.

### 4.5 Post mortem analyses

When mice reached deep anesthesia, blood was taken retro-orbital and analyzed in an automated hemocytometer (Mindray 5300Vet Hematology Analyzer). Peripheral blood for the determination of cytokine levels and glycoproteome analysis (see below) was obtained via cardiac puncture, kept at room temperature for 1 h and centrifuged at 1500 × g for 15 min without break in order to obtain serum. Aliquots were kept at -80°C until further analysis.

During necropsy, kidneys, spleen, liver, and brain were removed, weighed and either fixed immediately in Roti Histofix 4% (Carl Roth GmbH, Germany) and transferred into ethanol the next day, or kept on ice in 1-3 ml PBS until homogenization. For pathohistological analysis, a representative section from all organs was stained with hematoxylin and eosin and evaluated histopathological. For visualization of fungal structures, a PAS reaction and Grocott methenamine silver staining were performed. Histological preparations were digitalized with an Aperio CS2 (Whole Slide Scanner, Leica), and size, area, and numbers of inflammatory foci of the kidneys were quantified using the Image-Scope software (Leica). For development, validation, evaluation, and statistical analysis, all embedded in the visual programming language JIPipe^67^ (https://jipipe.hki-jena.de/), where the automated image processing of *Candida albicans* in mouse kidneys was realized using deep learning, please refer to Supplemental Methods. DAB (3, 3’-diaminobenzidine) immunohistochemistry on paraffin sections was performed after dewaxing (3 × xylol, 100% ethanol, 96% ethanol, 70% ethanol, distilled water, 10 min, respectively), followed by citrate buffer, pH 6.0, at 96°C for 30 min. After cooling, sections were washed with PBS and blocked with 1% BSA/PBS. Consecutive sections were and incubated with 10 µg/ml 1H7-4B, anti-NE (neutrophil elastase) antibody (Abcam, ab68672), and IgG control antibody (BioGenex, Rabbit NEGATIVE CONTROL) respectively, followed by goat-anti-rabbit-Conjugate (Vector, VEC-BA-1000) and developed using DAB (Carl Roth GmbH, Germany).

CFUs were determined by plating dilutions of organ homogenates on yeast extract-peptone-dextrose (YPD) agar plates with 80 µg/ml chloramphenicol. The detection limits were as follows: 50 CFU/g kidney and liver, respectively; 100 CFU/g spleen; 85 CFU/g brain. For Multiplex analysis (Cytokine Mouse Magnetic 20-Plex Panel for the Luminex platform, Thermo Fischer GmbH, Germany) or ELISA (mouse IL-6, IL-1β and TNF-Α Ready Set Go, both eBioscience, Germany; mouse IFNγ, BD Biosciences, Germany), homogenates were centrifuged immediately at 3000 × g, 4°C for 10 min and aliquots of supernatants were kept at -80°C until analysis.

For analysis of bone marrow cells, femurs were removed and kept in PBS on ice until cell isolation procedures. The expression of human CEACAMs in bone marrow-derived leukocytes fixed in 4% PFA/PBS was determined by flow cytometry. Cells were blocked in 500 µl of 10% BSA/PBS overnight at 4°C, treated with 1:30 mouse Fc block (eBioscience) for 30 min at 4°C, and stained and analyzed as described in the respective (supplemental) figure legends with the following antibodies and the corresponding isotype controls, respectively: Ly6C PE-Cy 7, Ly6G-APC, huCD66acde-PE (all REA, Miltenyi Biotec), B220 PerCP-Cy 5.5, CD3-APC (both eBioscience). Samples were analyzed on an Attune Acoustic Focusing Cytometer (Life Technologies, Thermo Fisher Scientific) using the FlowJo software version 10.0.6.

The expression of human CEACAMs in viable immune cell populations isolated from kidney, liver, and spleen was determined by flow cytometry. Cells were stained and analyzed on an Attune Acoustic Focusing Cytometer (Life Technologies, Thermo Fisher Scientific) using the Attune software v2.1 as described in the respective (supplemental) figure legends with the following antibodies or the corresponding isotype controls: CD45-PE, Ly-6G-PerCP-Vio700, CD3ε-PE-Vio770, CD19-APC, F4/80-FITC, CD45-PE, CD11c-PE-Vio770, CD335-APC (all REA, Miltenyi Biotec), and viability dye eFluor780 (eBioscience).

### 4.6 Glycoproteome analysis

Glycoprotein enrichment from serum: After thawing, 20 µl serum were added to 20 µl 4% SDS in PBS and heated for 5 min at 95°C. After cooling to room temperature, 60 µl of PBS were added and the mixture was centrifuged at 20.000 x g for 10 min at RT. The supernatant was used to enrich for glycoproteins as described^68^. Trypsin-released peptides and PNGase F-released N-glycopeptides were collected and dried in a SpeedVac (ThermoScientific). Shotgun MS-proteomics: Samples were reconstituted in 0.3% formic acid and peptide concentrations were measured using a NanoDrop spectrometer. 2.5 µg of tryptic and of deglycosylated peptides (former N-glycopeptides) were analyzed in each LC-MS/MS run in duplicates on an Orbitrap Fusion (Thermo Scientific) coupled to a Dionex Ultimate 3000 (Thermo Scientific) by a nanoelectrospray ion source. Samples were loaded on a 2 cm C18 trap column (Acclaim PepMap100, Thermo Scientific) and separated using a 2.5 h non-linear gradient (2-80% acetonitrile/0.1% formic acid, flow rate 300 nl/min) on a 50 cm C18 analytical column (75 µm i.d., PepMap RSLC, Thermo Scientific). Full MS scans were acquired with resolution 120.000 at m/z 400 in the Orbitrap analyzer (m/z range 370-1570, quadrupole isolation, isolation window 1.6). MS1 parent ions were fragmented by higher energy collisional dissociation (HCD, 30% collision energy) and 20 fragment ion spectra were acquired in the ion trap in rapid mode. The following conditions were used: spray voltage of 2.0 kV, heated capillary temperature of 275°C, S-lens RF level of 60%, maximum ion accumulation times of 50 ms (AGC 1 x 106) for full scans and 35 ms (AGC 1 x 104) for HCD. Protein Identification and Quantification: All RAW files were searched against the human UniProt database (Version 05.2016, reviewed sequences) with MaxQuant version 1.5.5.1 (Max Planck Institute of Biochemistry^69^. The parameters were set as follows: main search peptide tolerance: 4.5ppm; enzyme: trypsin, max. 2 missed cleavages; static modification: cysteine carbamidomethylation; variable modification in the tryptic peptide fraction: methionine oxidation; variable modification in PNGase F fractions: methionine oxidation and asparagine deamidation. PSM (peptide specific matches) and protein FDR was set to 0.01. For advanced identification the Second Peptide Search in MS2 spectra and the Match Between Runs feature were enabled. Label-free quantification of proteins with normalization was done in MaxQuant^69^. LFQ min. ratio count was set to one. Peptides from both fractions were integrated in the LFQ protein intensity calculations. Only unique and razor peptides, unmodified or modified, were used for quantification. LFQ protein intensities (see Supplementary Table S7) were then loaded into the Perseus framework (Max Planck Institute of Biochemistry)^70^. Known contaminants and reverse identified peptides/ proteins were discarded. Intensities were log(2) transformed and missing values were imputed from the normal distribution of the data set (width: 0.3, downshift 1.8). Two-sample T-test was used to calculate statistical differences of protein abundances in the compared groups. P-values were adjusted according to Benjamini and Hochberg^71^, and proteins demonstrating at least a two-fold expression difference and an adjusted p-value < 0.05 were considered to be significantly changed in abundance. The mass spectrometry proteomics data have been deposited to the ProteomeXchange Consortium via the PRIDE partner repository^72^ with the dataset identifier PXD061893 (reviewer account: Username, Password: ueEk3ftjm3Eq).

### 4.7 Isolation and analysis of bone marrow-derived neutrophils (BMNs)

Young adult mice were sacrificed by CO_2_ inhalation. BMN were isolated and assessed for purity and viability as described^15^. All assays were performed with freshly prepared BMN in Eppendorf tubes blocked with 10% BSA/PBS, for at least 1 h at 37°C.

Concentration of released MPO was determined in cell culture supernatants from BMN that were either left untreated or stimulated with live *C. albicans* yeast cells (MOI 10) for 60 min using the mouse MPO ELISA kit (Hycultec GmbH).

*C. albicans* killing assay was performed using the Colorimetric Cell Viability Kits III (XTT) from Promokine, Germany. 2 × 10^5^ neutrophils in 100 µl RPMI/10% FBS were left unstimulated or were stimulated with 2 × 10^5^ *C. albicans* cells per well (MOI 1) in a 96 well plate, for 30 min at 37°C, 5% CO_2_. *C. albicans* solution for standards (input) was kept on ice for the incubation times. Just before the following procedures, *C. albicans* standards (sensitivity: 6,500 CFU) were transferred to the 96 well plate. Triton X-100 was added to all wells to reach a final concentration of 0,3% and incubated for 10 min at 37°C, 5% CO_2_, in order inactivate neutrophils. Viable *C. albicans* cells were quantified by the addition of 50 µl XTT reaction mixture per well and incubation for 3-4 h at 37°C, 5% CO_2_. Absorbance was measured using a TECAN M200 at 450 nm and 630 nm (background). For calculations of *C. albicans* CFUs, background and blanks were subtracted.

Phagocytosis assays were performed as described^15^. Briefly, BMN were stimulated with FITC-labeled *C. albicans* yeast cells (MOI 20) for 20 min, fixed and counterstained for Hoechst33342 (Thermo Scientific), mouse anti-mouse CEACAM1 expressed on WT and CEABAC10 BMN (MSCC1, monoclonal mouse IgG1, Bernhard B. Singer, Essen / highly cross-adsorbed goat anti-mouse-Alexa546, Invitrogen) and extracellular *Candida* cells (rabbit-anti Candida, BP1006, Acris antibodies / highly cross-adsorbed goat anti-rabbit-Alexa633, Invitrogen). Micrographs were analyzed for phagocytosis by counting BMNs without contact to *Candida* cells, BMN with attached, extracellular *Candida* cells (FITC staining and anti-*Candida* antibody staining) and BMN with intracellular (phagocytosed) *Candida* cells (FITC staining only) with a confocal laser scanning microscope (Zeiss LSM 710) using the ZEN 2010 software (both Carl Zeiss Microscopy GmbH).

All flow cytometric analyses of BMN were performed on an Attune Acoustic Focusing Cytometer (Life Technologies, Thermo Fisher Scientific) using the Attune software v2.1.

The expression of CD11b, and human CEACAM3 and CEACAM6 and their relative fluorescence intensities on BMN was determined in cells either left untreated or stimulated with UV-killed *C. albicans* germ tubes (MOI 10) for 60 min. Cells were stained using viability dye eFluor780 (eBiosciences), anti-CD11b-APC (clone REA592, Miltenyi Biotec), and either 308/3-3 (anti-human CEACAM3/5, LeukoCom, Essen, Germany) / highly cross-adsorbed goat anti-mouse IgG1-PE (eBiosciences), or 1H7-4B (anti-human CEACAM6, LeukoCom, Essen, Germany) / highly cross-adsorbed goat anti-mouse IgG1-PE (eBiosciences), respectively. Note that 308/3-3 cross-reacts to CEACAM5^14^ that is not expressed on neutrophils (therefore we refer to 308/3-3 as “CEACAM3-specific” in the context of this publication). For the analysis of flow cytometry data, dead cells were excluded due to possible false positive staining.

For analysis of apoptosis, BMN were either left untreated for 2 h, or were treated for 2 h with UV-killed *C. albicans* germ tubes (MOI 10) and stained using the Annexin V Detection Kit APC (eBiosciences). Cells negative for Annexin V and propidium iodide were considered viable.

Analysis of intracellular reactive oxygen species was performed as described^15^. Briefly, BMN were pre-incubated for 15 min with 1.5 μg/ml dihydro-rhodamine 123 (DHR, Biomol GmbH) in calcium- and magnesium-free PBS/2.5% BSA and either left untreated or stimulated with UV-killed *C. albicans* germ tubes (MOI 10) for another 15 min. Cells were washed in calcium- and magnesium-free PBS, fixed in 1% PFA/PBS for 10 min, blocked with PBS/50% heat-inactivated fetal bovine serum, and washed with PBS/2% heat-inactivated fetal bovine serum.

### 4.8 Differentiation and analysis of bone marrow-derived monocytes (BMM) and macrophages (BMDM)

Young adult mice were sacrificed by CO_2_ inhalation. Isolated bone marrow cells were differentiated into BMM (non-adherent cells, 5-7 days) and BMDM (adherent cells, 7 days) in RPMI/25 ng/ml recombinant mouse M-CSF (ImmunoTools GmbH, Friesoythe, Germany).

The medium was exchanged every other day (non-adherent cells were collected by centrifugation). CEACAM6 expression was determined by flow cytometry using eFluor 780 (eBioscience, USA), and the following antibodies (all Miltenyi Biotec): CD66c-PE / REA414 (CEACAM6), F4/80-FITC / REA126, CD11b-APC / REA892, CD11c-PE-Vio770/REA754, CD62L-APC / REA828 as described in the respective (supplemental) figure legends. All flow cytometric analyses of BMN and BMDM, respectively, were performed on an Attune Acoustic Focusing Cytometer (Life Technologies, Thermo Fisher Scientific) using the Attune software v2.1.

For stimulation, *C. albicans* was opsonized (2 ×10^8^ / 1 ml of serum, incubation on ice for at least one hour, removal of serum by RPMI wash). In all stimulations longer than 6 h, treatment of the cells with Amphotericin B was included to prevent excessive growth of hyphae (1 µg/ml, added after 1 h of stimulation). For the phagocytosis assay, *C. albicans* cells were labelled with fluorescein isothiocyanate (FITC) prior to the experiment (see above).

For ELISAs, 5 ×10^5^ BMN and BMDM, respectively, in 500 µl medium were seeded in 24-well-plates, and were either left untreated, or were stimulated for 24 h at MOI 1. Concentrations of mouse cytokines IL-1β (eBioscience), IL-6 (Thermo Scientific), CCL2 (Becton Dickinson), and TNFα (Thermo Scientific) were determined according to manufacturer’s instructions.

To quantify the overall *C. albicans*-induced cell death, BMDM were seeded in 48-well-plates at a density of 2.5 × 10^5^ cells in 250 µl of medium. Following 24 h of *C. albicans* stimulation at MOI 1, SYTOX^TM^ Green (Invitrogen, USA) was added to the cells to a final concentration of 200 nM and incubated for 10 min in the dark at 37 °C 5 % CO_2_. Two images per well were taken at 20x magnification using the Vert.A1 + AxioCam MRc (Carl Zeiss, Germany) and exported as .tiff files from ZEN software (Carl Zeiss, Germany). Cells were counted using the multi-point tool in Image J (open source). Phase-contrast images were used to count all cells and dead cells were counted in the SYTOX^TM^ Green channel.

For analysis of phagocytosis, 2.5 ×10^5^ BMDM/well were seeded in collagen-coated 8-well-chambers (Permanox, Nunc Lab-TEK, Thermo Scientific) in 500 µl of medium and stimulated at an MOI 5 with FITC-labeled *C. albicans* for 20 min. Cells were washed twice with pre-warmed PBS, fixed with 250 µl 4% paraformaldehyde in PBS for 20 min at RT, and blocked with 200 µl 10 % BSA/PBS at 4°C overnight. Mouse Fc-block (eBioscience) was added for 30 min, and rabbit-anti-*Candida* antibody (BP1006, Acris antibodies) was added at 5 µg/ml for 1 h. Cells were washed three times with PBS, and goat anti-rabbit-Alexa Fluor 633 antibody (1:200, Invitrogen) and Hoechst33342 (1:1000, Thermo Scientific) in 5% BSA/PBS were added. Cells were washed three times with PBS, chambers were removed, and the slides were dipped consecutively in PBS, water and 70 % ethanol. After drying, slides were mounted with Vectashield mounting medium and allowed to harden in the dark overnight at RT and sealed using nail polish. Samples were analyzed with a Cell Observer Z1 microscope (Carl Zeiss, Germany) using the ZEN2010 software. Intracellular Ca and BMDM-associated extracellular Ca were counted, and the number of extracellular Ca was divided by the number of intracellular plus associated Ca, giving the phagocytosis index.

### 4.9 Human liver section analysis

Human PFA-fixed, paraffin-embedded liver sections were incubated at room temperature consecutively with xylol (2x 15 min), 100% ethanol, 96% ethanol, 70% ethanol, and with distilled water (each for 10 min), followed by Tris-EDTA buffer, pH 9.0, at boiling temperature for 15 min. After cooling, sections were washed with PBS and blocked with 1% BSA/PBS, incubated with monoclonal rabbit-anti-CD68 antibody (EPR20545, Abcam) and mouse anti-CEACAM6 1H7-4B, (LeukoCom, Essen, Germany) at 10 µg/ml in 0.5% BSA over night at 4°C in a humid chamber, washed and incubated with “Goat anti-Rabbit IgG (H+L) Cross-Adsorbed Secondary Antibody -Alexa Fluor 660” (Thermo Scientific), “Alexa Flour 546 Goat anti Mouse IgG (H+L)” (Invitrogen GmbH), each 1:200 in 0.5% BSA, and 1:5000 Hoechst 33342” (Thermo Scientific) for 1h at RT. Slides were washed twice with PBS. After drying, slides were mounted with Vectashield mounting medium (Biozol), allowed to harden in the dark overnight at RT, and sealed using nail polish. Samples were analyzed with an Axio Observer.Z1/7 microscope using the ZEN 2010 software (both Carl Zeiss Microscopy GmbH).

### 4.10 Human alveolar macrophage analysis

Cells from human bronchoalveolar lavage were stained for CD169-PE (marker for alveolar macrophages), CD66b-FITC (marker for human neutrophils), and CEACAM6 (CD66c-PE-Vio770) (all REA, Miltenyi Biotec) and analyzed on an Attune Acoustic Focusing Cytometer (Life Technologies, Thermo Fisher Scientific) using the Attune software v2.1.

### 4.11 Human peripheral monocyte analysis

Human PBMCs were isolated and stained directly using anti-CD14-FITC (REA, Miltenyi Biotec), anti-CD66c.PE-Vio770 (CEACAM6, REA, Miltenyi Biotec, and eFluor780. Cells were analyzed on an on an Attune Acoustic Focusing Cytometer (Life Technologies, Thermo Fisher Scientific) using the Attune software v2.1.

### 4.12 Statistical analysis

Except for proteomics data, statistical analysis was performed using GraphPad Prism 5.04 Software. For parametric data with 2 groups, unpaired, two-tailed Students t-test was performed; for matched pairs, a paired two-tailed Students t-test was performed. For non-matched parametric data with more than 2 groups, one-way ANOVA with Bonferroni post-tests was performed. In case of exponential data (CFUs, relative fluorescence intensity) log(10) transformed data were used for statistical analysis. In case of samples with no detectable CFU counts, statistical analysis was performed twice, inserting either 0.1 or the respective detection limit; the outcome was “not significant” in both cases. In the present manuscript, p values for the former analysis are given.

## 5 Data Availability

The datasets generated during and/or analyzed during the current study are available in the ProteomeXchange Consortium via the PRIDE partner repository72 with the dataset identifier PXD061893 (reviewer account: Username, Password: ueEk3ftjm3Eq). Code used for image analysis (see supplemental Material / supplemental method) is available at: https://asbdata.hki-jena.de/Klaile_PraetoriusEtAl2023_NatCommun

## 6 Acknowledgment

We thank Simone Tänzer, Moira Stark, Birgit Weber and Katja Schubert for their excellent technical assistance. We sincerely thank Bernhard B. Singer (†2024), a close colleague, friend, and leading expert in the field. His constant support, expertise, and groundbreaking design of CEACAM-specific antibodies were instrumental in raising the quality of CEACAM research to a level that would not have been possible without him. His visionary contributions significantly advanced CEACAM research worldwide.

## 7 Author Contributions

EK and HS conceived the study. EK, MMM, JS, AKB, SB, JE, TEK, SK and IDJ performed experiments. EK, MMM, JS, AKB, SB, JE, SK, AG, KD, OK, JPP, and MTF analyzed data. EK drafted the manuscript. EK, MMM, JS, AKB, SB, JE, TEK, SK, KD, OK, JPP, MTF, TB, AG, GM, IDJ and HS revised and approved the manuscript.

## 10 Contribution to the Field Statement

Systemic *C. albicans* infections result in an extremely high morbidity and mortality and still pose a major challenge regarding early diagnosis and treatment. Three myeloid cell types are first responding to systemic microbial infections: neutrophils, monocytes, and macrophages. The CEABAC10 transgenic mice express *Candida* receptors of the human CEACAM receptor family on their innate immune cells and on their mucosa. In the present study we show for the first time that the expression of human CEACAM6 on monocytes and macrophages can be essential to the regulation of the host response to pathogens. So far, only the expression of different CEACAM receptors on neutrophils and in mucosal epithelial cells was described to regulate the innate immune response to microbial infection. This work therefore points toward a completely new direction regarding the CEACAM-dependent regulation of immune responses.

## 12 Supplementary Material

Supplementary Figures and Method (PDF file):

Figure S1: CEABAC10 mice are more susceptible to systemic *C. albicans* infection. Survival analysis with an infection dose of 2.5×10^4^ CFU/g body weight. Related to Figure 1.

Figure S2: CEABAC10 mice display enhanced cytokine levels in peripheral blood during systemic *C. albicans* infection. Multiplex/ELISA. Related to Figure 1.

Figure S3: No differences in bone marrow compositions between WT and CEABAC10 mice during systemic *C. albicans* infection. Flow cytometry.

Figure S4: CEABAC10 mice have reduced white blood cell and platelets in counts in their peripheral blood during systemic *C. albicans* infection. Automated cytometric analysis of peripheral blood. Related to Figure 1.

Figure S5: CEABAC10 mice have enhanced cytokine concentrations in their kidneys during systemic *C. albicans* infection. Multiplex/ELISA. Related to Figure 3.

Figure S6: Immune cell populations in kidneys of WT and CEABAC10 mice during systemic *C. albicans* infection and automated image analysis of *Candida* cells kidney sections. Related to Figure 3.

Figure S7: Gating of immune cell populations in kidneys. Related to Figure 3 and Figure S6.

Figure S8: BMN gating. Related to Figure 4.

Figure S9: CEABAC10 mice have enhanced CCL3 and CXCL10 concentrations in their livers during systemic *C. albicans* infection. Multiplex/ELISA. Related to Figure 5.

Figure S10: Immune cell populations in livers of WT and CEABAC10 mice during systemic *C. albicans* infection. Flow cytometry and IHC. Related to Figure 5.

Figure S11: Gating of immune cell populations in livers. Related to Figure 5 and Figure S10.

Figure S12: Immune cell populations in spleen during systemic *C. albicans* infection. Flow cytometry and IHC. Related to Figure 5.

Figure S13: Gating of immune cell populations in spleens. Related to Figure S12.

Figure S14: CEABAC10 mice display brain hemorrhage but no increase in encephalitis 72 h post-*C. albicans* infection. Fungal load, histopathologic analysis, IHC.

Figure S15: Gating of CEACAM6^+^ myeloid cells in organs of CEABAC10 mice during systemic *C. albicans* infection. Related to Figure 6

Figure S16: Gating of bone marrow-derived macrophages and monocytes. Related to Figures 7 and 8.

Figure S17: CEACAM6 expression on human liver macrophages. Figure S18: CEACAM6 expression on human alveolar macrophages.

Figure S19: No CEACAM6 expression on human peripheral monocytes.

Supplementaal Method: Automated image analysis of *C. albicans* cells in kidney sections. Development, evaluation and detailed results.

Supplementary Tables (XLSX file):

Supplementary Tables S1, S2, S3, S4, S5, and S6: Serum glycoproteome analyses. Comparisons of glycoproteomes and pathway analysis of these comparisons, Tables S1, S2, S3, S4, S5 and S6 relate to Figure 2b, 2c, 2g, 2h, 2f, and 2i, respectively.

Supplementary Table S7: Protein groups (results of glycoproteome analysis).

## 13 References

1. Eggimann P, Pittet D. Candida colonization index and subsequent infection in critically ill surgical patients: 20 years later. Intensive Care Med 40, 1429–1448 (2014).

2. Lopes JP, Lionakis MS. Pathogenesis and virulence of Candida albicans. Virulence 13, 89–121 (2022).

3. Lionakis MS. New insights into innate immune control of systemic candidiasis. Med Mycol 52, 555–564 (2014).

4. Goyal S, Castrillon-Betancur JC, Klaile E, Slevogt H. The Interaction of Human Pathogenic Fungi With C-Type Lectin Receptors. Front Immunol 9, 1261 (2018).

5. Bourgeois C, Kuchler K. Fungal pathogens-a sweet and sour treat for toll-like receptors. Front Cell Infect Microbiol 2, 142 (2012).

6. Klaile E, et al. Binding of Candida albicans to Human CEACAM1 and CEACAM6 Modulates the Inflammatory Response of Intestinal Epithelial Cells. mBio 8, (2017).

7. Bonsignore P, Kuiper JWP, Adrian J, Goob G, Hauck CR. CEACAM3-A Prim(at)e Invention for Opsonin-Independent Phagocytosis of Bacteria. Front Immunol 10, 3160 (2019).

8. Tchoupa AK, Schuhmacher T, Hauck CR. Signaling by epithelial members of the CEACAM family - mucosal docking sites for pathogenic bacteria. Cell Commun Signal 12, 27 (2014).

9. Heinrich A, et al. Moraxella catarrhalis induces CEACAM3-Syk-CARD9-dependent activation of human granulocytes. Cell Microbiol 18, 1570–1582 (2016).

10. Ambrosi C, Scribano D, Sarshar M, Zagaglia C, Singer BB, Palamara AT. Acinetobacter baumannii Targets Human Carcinoembryonic Antigen-Related Cell Adhesion Molecules (CEACAMs) for Invasion of Pneumocytes. mSystems 5, (2020).

11. Javaheri A, et al. Helicobacter pylori adhesin HopQ engages in a virulence-enhancing interaction with human CEACAMs. Nat Microbiol 2, 16189 (2016).

12. Tegtmeyer N, et al. Type IV secretion of Helicobacter pylori CagA into oral epithelial cells is prevented by the absence of CEACAM receptor expression. Gut Pathog 12, 25 (2020).

13. Sarantis H, Gray-Owen SD. Defining the roles of human carcinoembryonic antigen-related cellular adhesion molecules during neutrophil responses to Neisseria gonorrhoeae. Infect Immun 80, 345–358 (2012).

14. Klaile E, et al. Antibody ligation of CEACAM1, CEACAM3, and CEACAM6, differentially enhance the cytokine release of human neutrophils in responses to Candida albicans. *Cell Immunol* **371**, 104459 (2022).

15. Klaile E, et al. Unaltered Fungal Burden and Lethality in Human CEACAM1-Transgenic Mice During Candida albicans Dissemination and Systemic Infection. Front Microbiol 10, 2703 (2019).

16. Hirai A, et al. Role of mouse hepatitis virus (MHV) receptor murine CEACAM1 in the resistance of mice to MHV infection: studies of mice with chimeric mCEACAM1a and mCEACAM1b. J Virol 84, 6654–6666 (2010).

17. Zhang Z, et al. CEACAM1 regulates the IL-6 mediated fever response to LPS through the RP105 receptor in murine monocytes. BMC Immunol 20, 7 (2019).

18. Zoller J, et al. CEACAM1 regulates CD8(+) T cell immunity and protects from severe pathology during Citrobacter rodentium induced colitis. Gut Microbes 11, 1790–1805 (2020).

19. Zebhauser R, Kammerer R, Eisenried A, McLellan A, Moore T, Zimmermann W. Identification of a novel group of evolutionarily conserved members within the rapidly diverging murine Cea family. Genomics 86, 566–580 (2005).

20. Chan CH, Stanners CP. Novel mouse model for carcinoembryonic antigen-based therapy. Mol Ther 9, 775–785 (2004).

21. Carvalho FA, et al. Crohn’s disease adherent-invasive Escherichia coli colonize and induce strong gut inflammation in transgenic mice expressing human CEACAM. J Exp Med 206, 2179–2189 (2009).

22. Denizot J, et al. Adherent-invasive Escherichia coli induce claudin-2 expression and barrier defect in CEABAC10 mice and Crohn’s disease patients. Inflamm Bowel Dis 18, 294–304 (2012).

23. Sintsova A, Guo CX, Sarantis H, Mak TW, Glogauer M, Gray-Owen SD. Bcl10 synergistically links CEACAM3 and TLR-dependent inflammatory signalling. Cell Microbiol 20, (2018).

24. Hebecker B, et al. Dual-species transcriptional profiling during systemic candidiasis reveals organ-specific host-pathogen interactions. Sci Rep 6, 36055 (2016).

25. Jacobsen ID, Luttich A, Kurzai O, Hube B, Brock M. In vivo imaging of disseminated murine Candida albicans infection reveals unexpected host sites of fungal persistence during antifungal therapy. J Antimicrob Chemother 69, 2785–2796 (2014).

26. Iba T, Umemura Y, Wada H, Levy JH. Roles of Coagulation Abnormalities and Microthrombosis in Sepsis: Pathophysiology, Diagnosis, and Treatment. Arch Med Res 52, 788–797 (2021).

27. Lionakis MS, Lim JK, Lee CC, Murphy PM. Organ-specific innate immune responses in a mouse model of invasive candidiasis. J Innate Immun 3, 180–199 (2011).

28. Thomas J, et al. CEACAMS 1, 5, and 6 in disease and cancer: interactions with pathogens. Genes Cancer 14, 12-29 (2023).

29. Sonego F, et al. Paradoxical Roles of the Neutrophil in Sepsis: Protective and Deleterious. Front Immunol 7, 155 (2016).

30. Metzemaekers M, Gouwy M, Proost P. Neutrophil chemoattractant receptors in health and disease: double-edged swords. Cell Mol Immunol 17, 433–450 (2020).

31. Kuroki M, Yamanaka T, Matsuo Y, Oikawa S, Nakazato H, Matsuoka Y. Immunochemical analysis of carcinoembryonic antigen (CEA)-related antigens differentially localized in intracellular granules of human neutrophils. Immunol Invest 24, 829–843 (1995).

32. Skubitz KM, Skubitz AP. Two new synthetic peptides from the N-domain of CEACAM1 (CD66a) stimulate neutrophil adhesion to endothelial cells. Biopolymers 96, 25–31 (2011).

33. Skubitz KM, Skubitz AP. Interdependency of CEACAM-1, -3, -6, and -8 induced human neutrophil adhesion to endothelial cells. J Transl Med 6, 78 (2008).

34. Wu X, et al. Human Liver Macrophage Subsets Defined by CD32. Front Immunol 11, 2108 (2020).

35. Baharom F, Rankin G, Blomberg A, Smed-Sorensen A. Human Lung Mononuclear Phagocytes in Health and Disease. Front Immunol 8, 499 (2017).

36. Gray-Owen SD, Blumberg RS. CEACAM1: contact-dependent control of immunity. Nat Rev Immunol 6, 433–446 (2006).

37. Behrens IK, et al. The HopQ-CEACAM Interaction Controls CagA Translocation, Phosphorylation, and Phagocytosis of Helicobacter pylori in Neutrophils. mBio 11, (2020).

38. Martinez-Medina M, et al. Western diet induces dysbiosis with increased E coli in CEABAC10 mice, alters host barrier function favouring AIEC colonisation. Gut 63, 116–124 (2014).

39. Spellberg B, Ibrahim AS, Edwards JE, Jr., Filler SG. Mice with disseminated candidiasis die of progressive sepsis. J Infect Dis 192, 336–343 (2005).

40. Lionakis MS, et al. Chemokine receptor Ccr1 drives neutrophil-mediated kidney immunopathology and mortality in invasive candidiasis. PLoS Pathog 8, e1002865 (2012).

41. Lionakis MS, Albert ND, Swamydas M, Lee CR, Loetscher P, Kontoyiannis DP. Pharmacological Blockade of the Chemokine Receptor CCR1 Protects Mice from Systemic Candidiasis of Hematogenous Origin. Antimicrob Agents Chemother 61, (2017).

42. Yano J, Peters BM, Noverr MC, Fidel PL, Jr. Novel Mechanism behind the Immunopathogenesis of Vulvovaginal Candidiasis: “Neutrophil Anergy”. Infect Immun 86, (2018).

43. Souto FO, et al. Liver X Receptor Activation Impairs Neutrophil Functions and Aggravates Sepsis. J Infect Dis 221, 1542–1553 (2020).

44. Wang YY, et al. Liver X receptor agonist GW3965 dose-dependently regulates lps-mediated liver injury and modulates posttranscriptional TNF-alpha production and p38 mitogen-activated protein kinase activation in liver macrophages. Shock 32, 548–553 (2009).

45. Myhre AE, et al. Liver X receptor is a key regulator of cytokine release in human monocytes. Shock 29, 468–474 (2008).

46. Ruiqi W, et al. Monocyte-derived macrophages contribute to the deterioration of immunological liver injury in mice. Int Immunopharmacol 124, 111036 (2023).

47. Austermeier S, Kasper L, Westman J, Gresnigt MS. I want to break free - macrophage strategies to recognize and kill Candida albicans, and fungal counter-strategies to escape. Curr Opin Microbiol 58, 15–23 (2020).

48. Smeekens SP, et al. The classical CD14(+)(+) CD16(-) monocytes, but not the patrolling CD14(+) CD16(+) monocytes, promote Th17 responses to Candida albicans. Eur J Immunol 41, 2915–2924 (2011).

49. Ngo LY, Kasahara S, Kumasaka DK, Knoblaugh SE, Jhingran A, Hohl TM. Inflammatory monocytes mediate early and organ-specific innate defense during systemic candidiasis. J Infect Dis 209, 109–119 (2014).

50. Langenhorst D, et al. Soluble Enolase 1 of Candida albicans and Aspergillus fumigatus Stimulates Human and Mouse B Cells and Monocytes. J Immunol 211, 804–815 (2023).

51. Zhang X, Liu H, Hashimoto K, Yuan S, Zhang J. The gut-liver axis in sepsis: interaction mechanisms and therapeutic potential. Crit Care 26, 213 (2022).

52. Xu S, Shinohara ML. Tissue-Resident Macrophages in Fungal Infections. Front Immunol 8, 1798 (2017).

53. Lionakis MS, et al. CX3CR1-dependent renal macrophage survival promotes Candida control and host survival. J Clin Invest 123, 5035–5051 (2013).

54. Strnad P, Tacke F, Koch A, Trautwein C. Liver - guardian, modifier and target of sepsis. Nat Rev Gastroenterol Hepatol 14, 55–66 (2017).

55. Bauer M, Press AT, Trauner M. The liver in sepsis: patterns of response and injury. Curr Opin Crit Care 19, 123-127 (2013).

56. MacCallum DM, Castillo L, Brown AJ, Gow NA, Odds FC. Early-expressed chemokines predict kidney immunopathology in experimental disseminated Candida albicans infections. PLoS One 4, e6420 (2009).

57. Bloos F, Reinhart K. Rapid diagnosis of sepsis. Virulence 5, 154–160 (2014).

58. Honda S, et al. Marginal zone B cells exacerbate endotoxic shock via interleukin-6 secretion induced by Fcalpha/muR-coupled TLR4 signalling. Nat Commun 7, 11498 (2016).

59. Drummond RA, Lionakis MS. Organ-specific mechanisms linking innate and adaptive antifungal immunity. Semin Cell Dev Biol 89, 78–90 (2019).

60. Drummond RA. What fungal CNS infections can teach us about neuroimmunology and CNS-specific immunity. Semin Immunol 67, 101751 (2023).

61. Drummond RA, et al. CARD9(+) microglia promote antifungal immunity via IL-1beta- and CXCL1-mediated neutrophil recruitment. Nat Immunol 20, 559–570 (2019).

62. Gil ML, Gozalbo D. Role of Toll-like receptors in systemic Candida albicans infections. Front Biosci (Landmark Ed) 14, 570–582 (2009).

63. Taylor PR, et al. Dectin-1 is required for beta-glucan recognition and control of fungal infection. Nat Immunol 8, 31–38 (2007).

64. Burtin P, Quan PC, Sabine MC. Nonspecific cross reacting antigen as a marker for human polymorphs, macrophages and monocytes. Nature 255, 714–716 (1975).

65. Bordes M, Knobel S, Martin F. Carcinoembryonic antigen (CEA) and related antigens in blood cells and hematopoietic tissues. Eur J Cancer (1965) 11, 783–786 (1975).

66. Nap M, et al. Specificity and affinity of monoclonal antibodies against carcinoembryonic antigen. Cancer Res 52, 2329–2339 (1992).

67. Gerst R, Cseresnyes Z, Figge MT. JIPipe: visual batch processing for ImageJ. Nat Methods 20, 168–169 (2023).

68. Muller MM, et al. Global analysis of glycoproteins identifies markers of endotoxin tolerant monocytes and GPR84 as a modulator of TNFalpha expression. Sci Rep 7, 838 (2017).

69. Cox J, Mann M. MaxQuant enables high peptide identification rates, individualized p.p.b.-range mass accuracies and proteome-wide protein quantification. Nat Biotechnol 26, 1367–1372 (2008).

70. Tyanova S, et al. The Perseus computational platform for comprehensive analysis of (prote)omics data. Nat Methods 13, 731–740 (2016).

71. Benjamini Y, Drai D, Elmer G, Kafkafi N, Golani I. Controlling the false discovery rate in behavior genetics research. Behav Brain Res 125, 279–284 (2001).

72. Perez-Riverol Y, et al. The PRIDE database at 20 years: 2025 update. Nucleic Acids Res 53, D543–d553 (2025).

