## Supplementary material for "Expression of human CEACAMs promotes inflammation and organ damage during systemic *Candida albicans* infection in mice": Supplemetnal Figures and Supplemental Method

### Supplemental Material

#### 1. Supplemental Figures

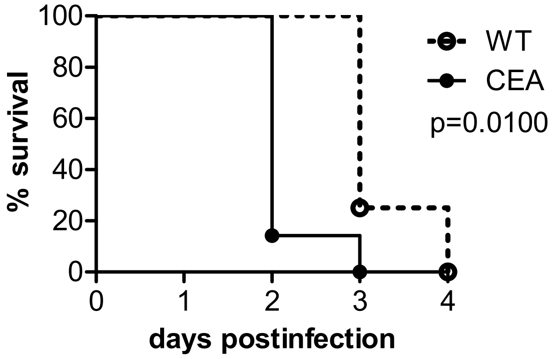


**Figure S1: CEABAC10 mice are more susceptible to systemic *C. albicans* infection.** CEABAC10 mice (CEA, N=7) and wild type littermates (WT, N=4) were injected with $2.5 \times 10^4$ CFU/g body weight *C. albicans* yeast cells into the lateral tail vein and sacrificed when they reached a humane endpoint. Log rank test (Mantel Cox) was used for statistical analysis of survival. Related to Figure 1a.

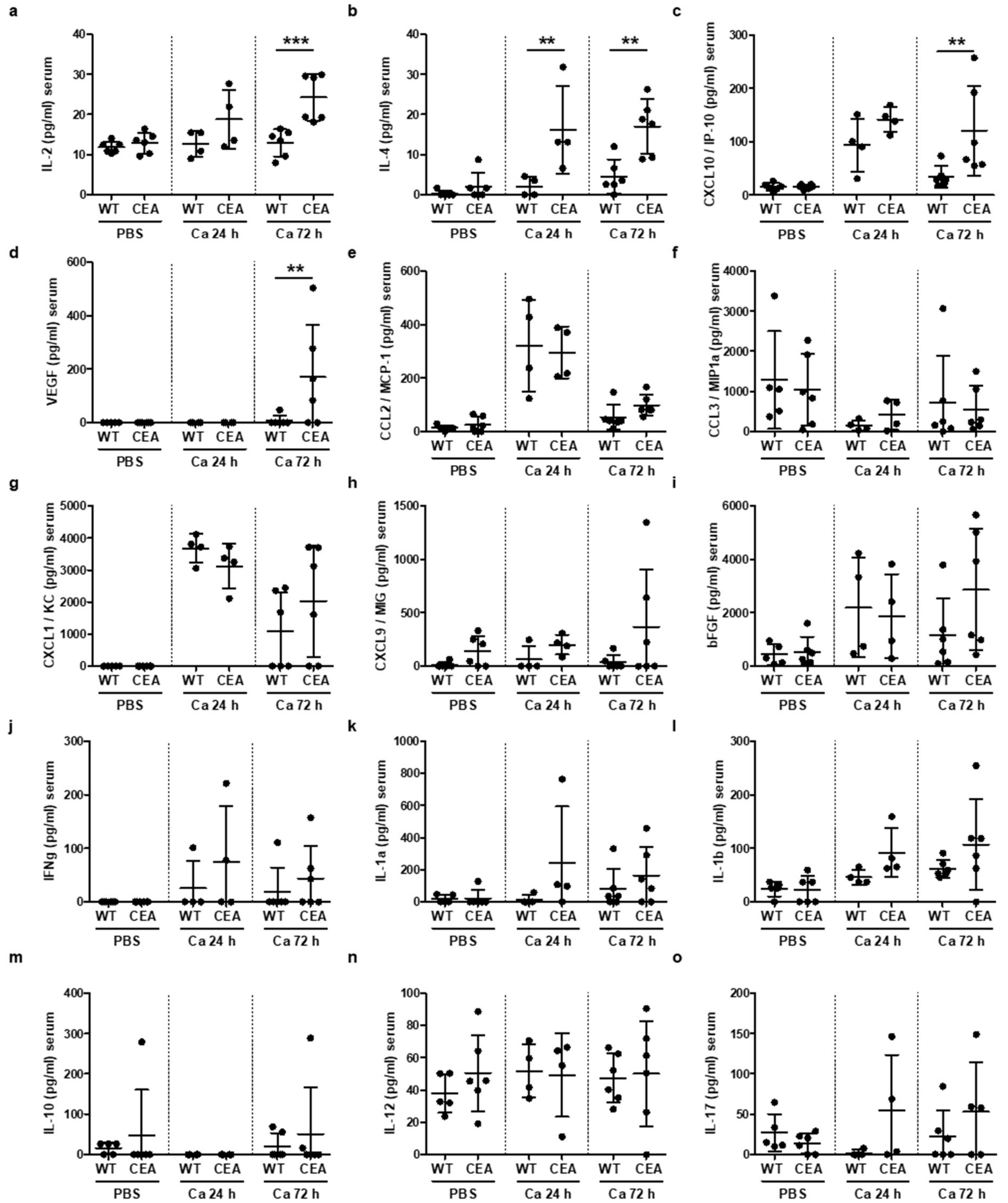


**Figure S2: CEABAC10 mice display enhanced cytokine levels in peripheral blood during systemic *C. albicans* infection.** CEABAC10 mice (CEA) and their wild type littermates (WT) were either injected with PBS or infected with $1\times10^4$ CFU/g body weight, respectively, and were sacrificed after 24 h or 72 h (6 WT and 6 CEABAC10 for PBS and Ca 72 h, respectively, and 4 WT and 4 CEABAC10 for 24 h Ca). Cytokine levels in peripheral blood were determined by multiplex assay (Luminex) or ELISA. Note that IL-5 and IL-13 were not detected in any blood sample. Statistics: One-Way-ANOVA and Bonferroni Multiple Comparison Test: $^{**}p<0.01$ $^{***}p<0.005$. Data are combined from two independent experiments. Graphs give data points with mean and standard deviation. Related to Figure 1b-d.

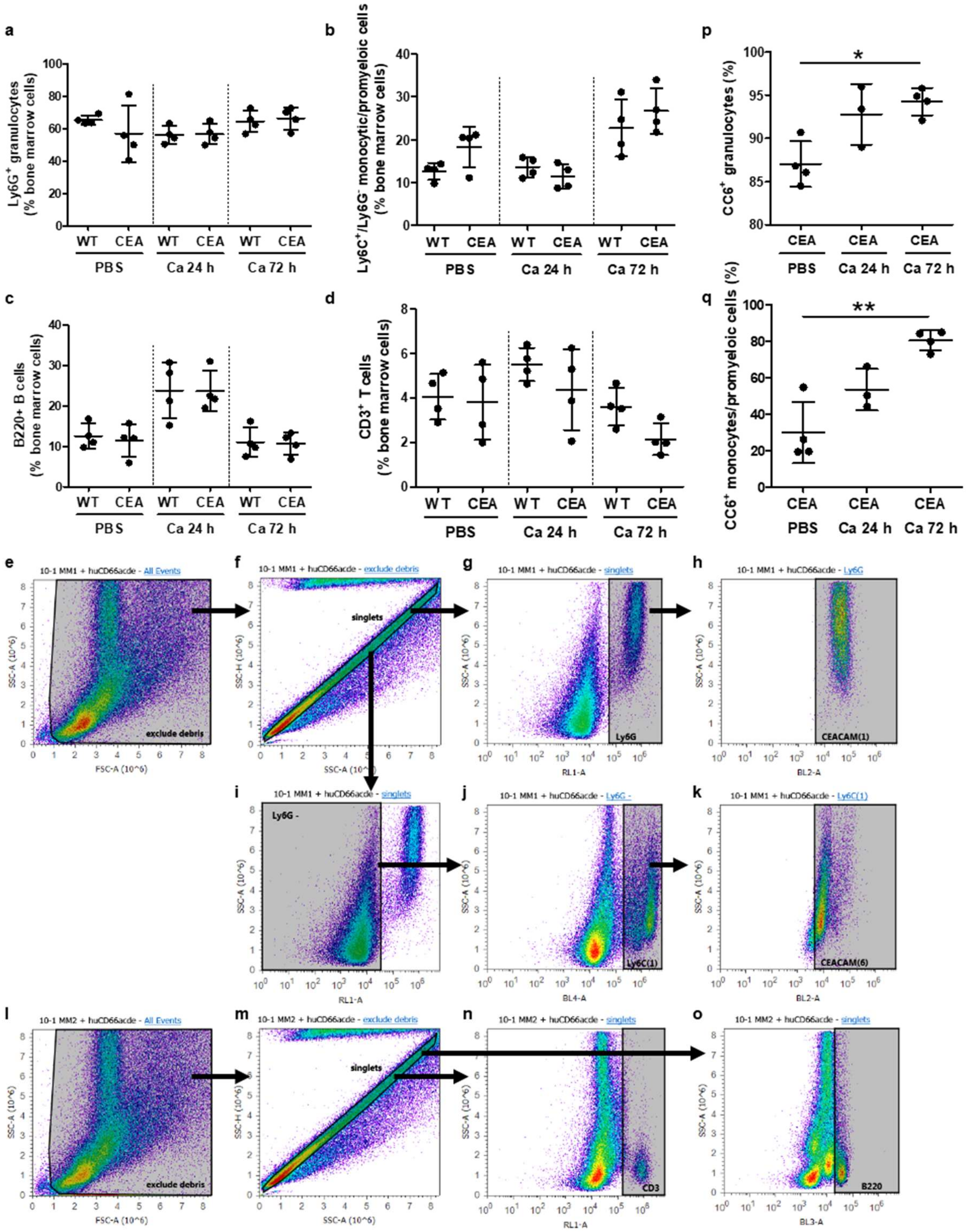


**Figure S3: No differences in bone marrow compositions between WT and CEABAC10 mice during systemic *C. albicans* infection.** CEABAC10 mice (CEA) and their wild type littermates (WT) were either injected with PBS or infected with $1\times10^4$ CFU/g body weight, respectively, and were sacrificed after 24 h or 72 h (4 per group). Bone marrow cells were isolated and stained for either Ly6G/Ly6C/CD66acde (a, b, e-k) or B220/CD3 (c, d, l-o). Cells were analyzed on an on an Attune Acoustic Focusing Cytometer (Life Technologies, Thermo Fisher Scientific) using the Attune software v2.1. (a, b) Single bone marrow cells were gated for $Ly6G^+$ neutrophilic granulocytes (a), or $Ly6G^-/Ly6C^+$/ monocytic/promyeloid cells (b) as shown in (e-g) and (e, f, j), respectively. (c, d) Single bone marrow cells were gated for B220$^{+}$ B lymphocytes (c) or CD3$^{+}$ T lymphocytes (d) as shown in (l-n) and (l, m, o), respectively. (h, k) Ly6G$^{+}$ neutrophilic granulocytes (g) and Ly6G$^{-}$/Ly6C$^{+}$/ monocytic/promyeloid cells (j) were analyzed for their human CEACAM3/6 expression (p, q, respectively; related to Figure 6). Graphs give the percentage of the respective cell type of total bone marrow cells (a-d) or of the parental population (p, q). Corresponding isotype controls were used to determine positive/negative subpopulations, respectively. Statistics: One-Way-ANOVA and Bonferroni Multiple Comparison Test: *$p<0.05$ **$p<0.01$. Data are from one experiment with 4 mice per group. Graphs give data points with mean and standard deviation.

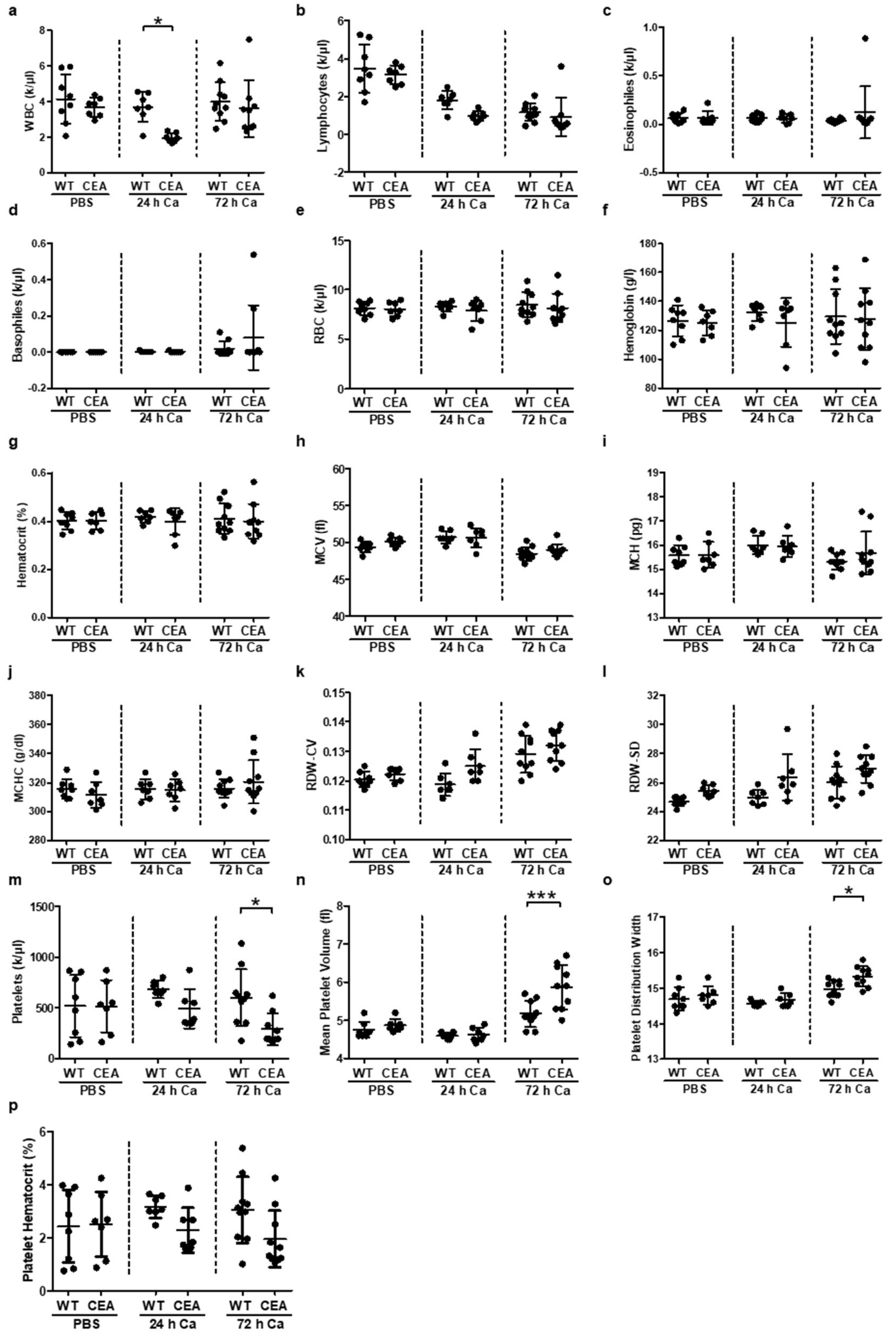


**Figure S4: CEABAC10 mice have reduced white blood cell and platelets in counts in their peripheral blood during systemic *C. albicans* infection.** CEABAC10 mice (CEA) and their wild type littermates (WT) were either injected with PBS or infected with $1\times10^4$ CFU/g body weight, respectively, and were sacrificed after 24 h (7 WT and 7 CEABAC10 for PBS and Ca, respectively) or 72 h (9 WT and 11 CEABAC10). Blood parameters were determined in an automated hemocytometer (a-p). Statistics: One-Way-ANOVA and Bonferroni Multiple Comparison Test: *p<0.05 ***p<0.005. Data are combined from two independent experiments. Graphs give data points with mean and standard deviation. Related to Figure 1e, f.

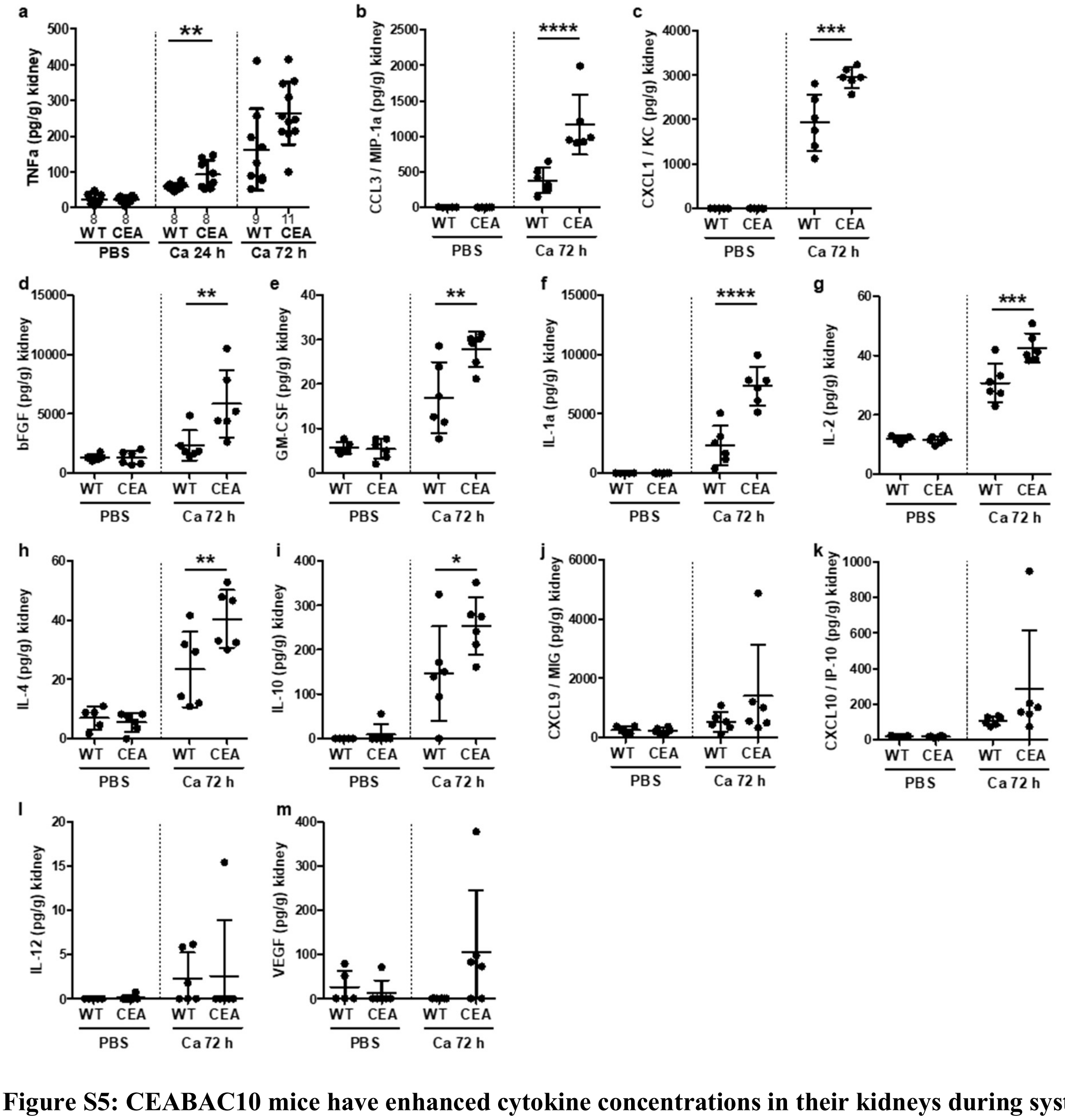


**Figure S5: CEABAC10 mice have enhanced cytokine concentrations in their kidneys during systemic *C. albicans* infection.** CEABAC10 mice (CEA) and their wild type littermates (WT) were either injected with PBS or infected with $1\times10^4$ CFU/g body weight, respectively, and were sacrificed after 24 h or 72 h. (a-m) Concentrations of cytokines were determined in kidney homogenates by multiplex assay (Luminex, 6 per group). For a, up to 5 additional samples per group were analyzed via ELISA. Data are combined from two independent experiments. Statistics: One-Way-ANOVA and Bonferroni Multiple Comparison Test: *p<0.05 **p<0.01 ***p<0.005 ****p<0.001. Note that IL-5, IL-13, IL-15 and IFNγ were not detected in any kidney sample. Graphs give data points with mean and standard deviation. Related to Figure 3i-k.

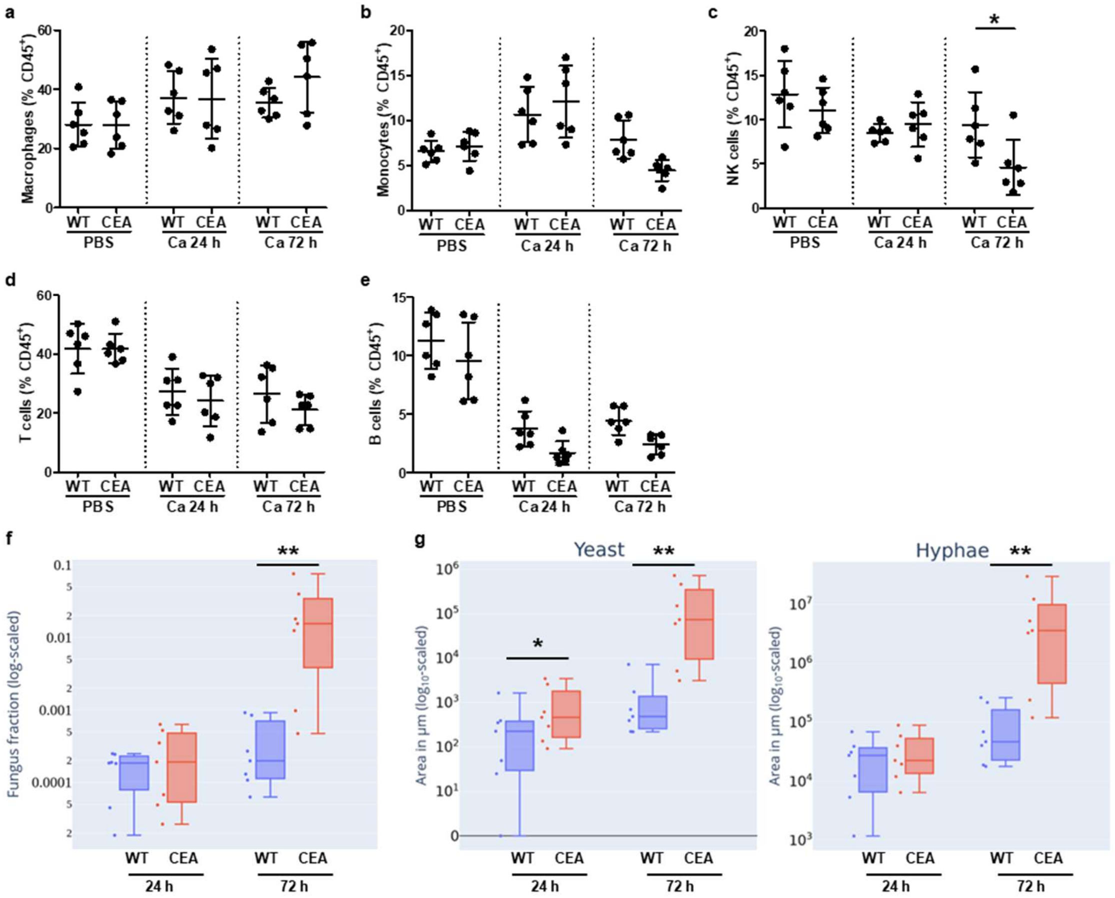


**Figure S6: Immune cell populations in kidneys of WT and CEABAC10 mice during systemic *C. albicans* infection and automated quantification of *Candida* morphotypes by image analysis in kidney sections.** CEABAC10 mice (CEA) and their wild type littermates (WT) were either injected with PBS or infected with $1\times10^4$ CFU/g body weight, respectively, and were sacrificed after 24 h or 72 h. (a-e) Immune cells were isolated from kidneys and stained for viability dye/CD45/Ly6G/CD3/CD19 and viability dye/CD45/F4/80/CD11c/CD335, respectively, and analyzed by flow cytometry (N=6 per group and staining, see Figure S7 for gating). Macrophages (a), monocytes (b), NK cells (c), T lymphocytes (d), and $CD19^+$ B lymphocytes (e) are given as the percentage of $CD45^+$ cells with mean and standard deviation. Note that data displayed in Figure S6d, e are from the same data sets as Figure 3p. Statistics: (a-e) One-Way-ANOVA and Bonferroni Multiple Comparison Test, *p<0.05. (f, g) Grocott-silver-stained Kidney sections were automated and objectively quantified using deep learning-based image analysis. (f) Computationally annotated total fungal fraction per kidney. The fungal fraction is defined by the area covered by fungal cells divided by kidney area. **p-value < 0.002331, effect size = 1.22 (related to Figure 3l). (g) Areas covered by yeast and hyphae morphotypes per mouse kidney section. While serial plating resulted in no significant differences 24 h p.i., the automated segmentation revealed an increase in yeast cells in CEABAC10 kidneys already at this time point (*p-value < 0.01, effect size = 0.69). Consistent with both, the serial Plating (Figure 3l) and the hyphae score determined by pathologists (Figure 3o), both, yeast cells (**p-value < 0.002, effect size = 1.07) and hyphae (**p-value < 0.004, effect size = 1.03) are significantly increased in CEABAC10 kidneys72 h p.i. Data are combined from two independent experiments. (f, g) Statistical tests for the fungal fraction were performed using the unpaired Wilcoxon rank sum test and using the R package "effectsize" to calculate the Hedge ‘g effect size[73]. The ranges of Hedges ‘g effect size magnitudes are referred to as being negligible for | Statistical Methods for Meta-Analysis g| < 0.2, small for |g| < 0.5, medium for |g| < 0.8 and large for |g| ≥ 0.8.

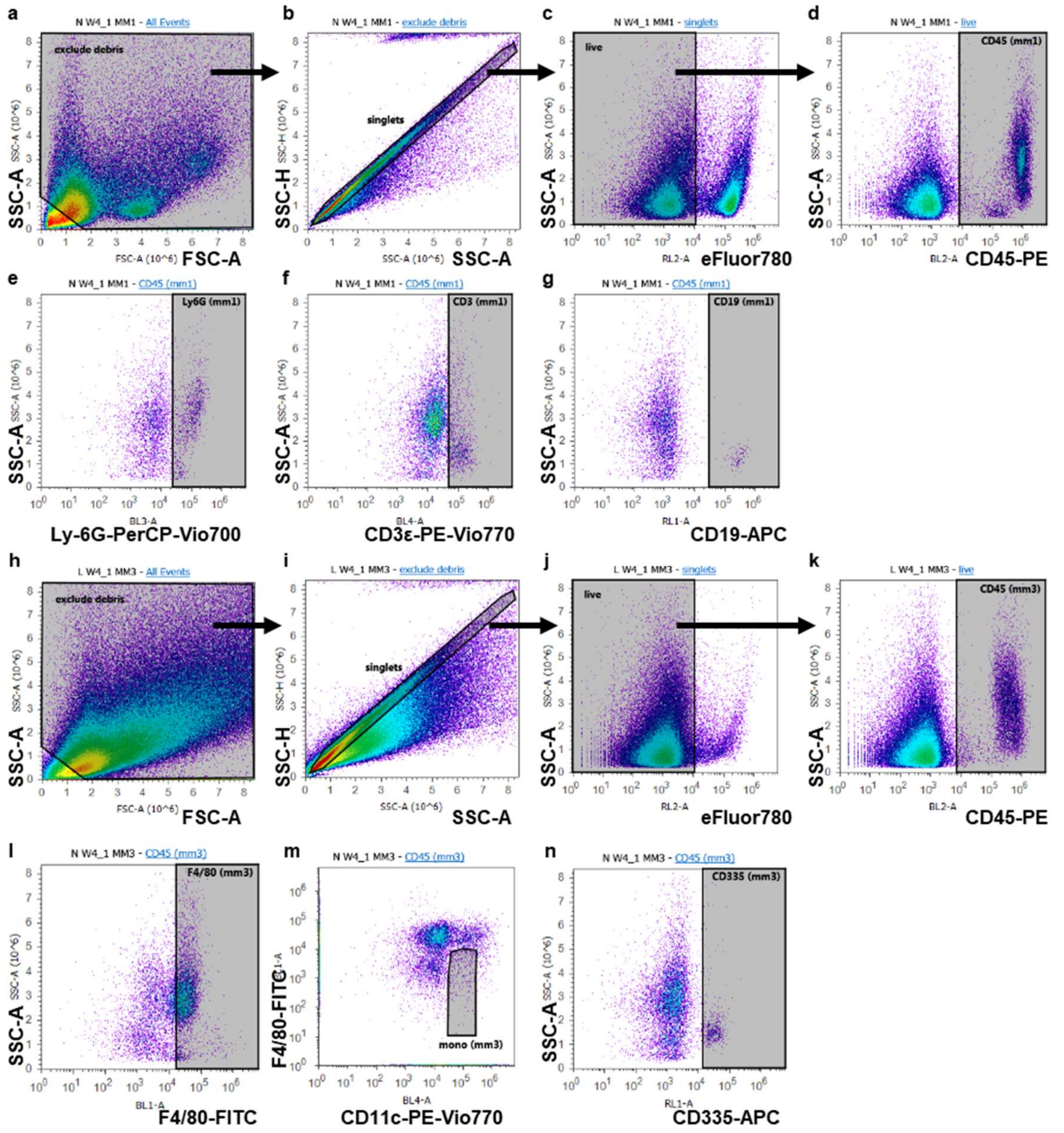


**Figure S7: Gating of immune cell populations in kidneys.** Representative panels are from a CEABAC10 mouse 72 h p.i.: (infected with $1\times10^4$ CFU/g body weight). Immune cells were isolated from kidneys and stained for CD45-PE, Ly-6G-PerCP-Vio700, CD3ε-PE-Vio770, CD19-APC, and viability dye eFluor780 (a-g), and F4/80-FITC, CD45-PE, CD11c-PE-Vio770, CD335-APC, viability dye eFluor780 (h-n), respectively, and analyzed by flow cytometry. (a-g) Single (b) / viable (c) / $CD45^+$ cells (d) were gated for $Ly6G^+$ granulocytes (e), $CD3^+$ T lymphocytes (f), or $CD19^+$ B lymphocytes (g). (h-m) Single (i) / viable (j) / $CD45^+$ cells (k) were gated for $F4/80^{hi}$ macrophages (l), $CD11c^{int}/F4/80^{lo/neg}$ monocytes (m), and $CD335^+$ NK cells (n). Corresponding isotype controls were used to determine positive/negative subpopulations, respectively. Cells were analyzed on an on an Attune Acoustic Focusing Cytometer (Life Technologies, Thermo Fisher Scientific) using the Attune software v2.1. Related to Figures 3 and S6.

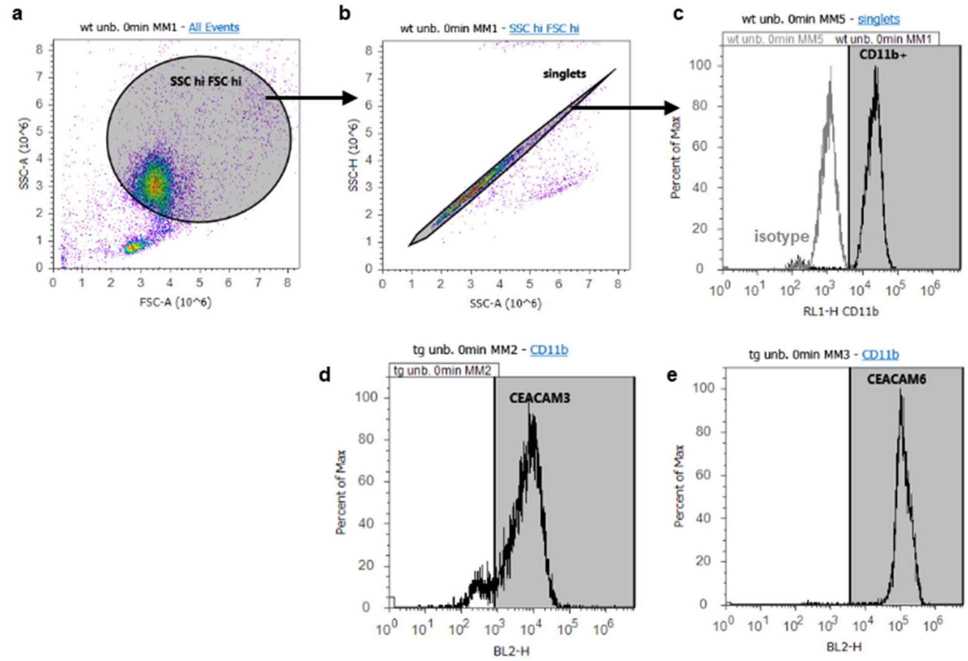


**Figure S8: BMN gating.** CEABAC10 BMN were stained for CD11b and either CEACAM3 (c) or CEACAM6 (d), respectively, and analyzed by flow cytometry. Single cells were gated for CD11b$^+$ BMN, and CD11b$^+$ CEABAC10 BMN were subsequently analyzed for their CEACAM expression (d, e). Corresponding isotype controls were used to determine positive/negative subpopulations, respectively. Panels show representative results N=4). Note that WT BMN were only stained and analyzed for CD11b. Cells were analyzed on an on an Attune Acoustic Focusing Cytometer (Life Technologies, Thermo Fisher Scientific) using the Attune software v2.1. Related to Figure 4.

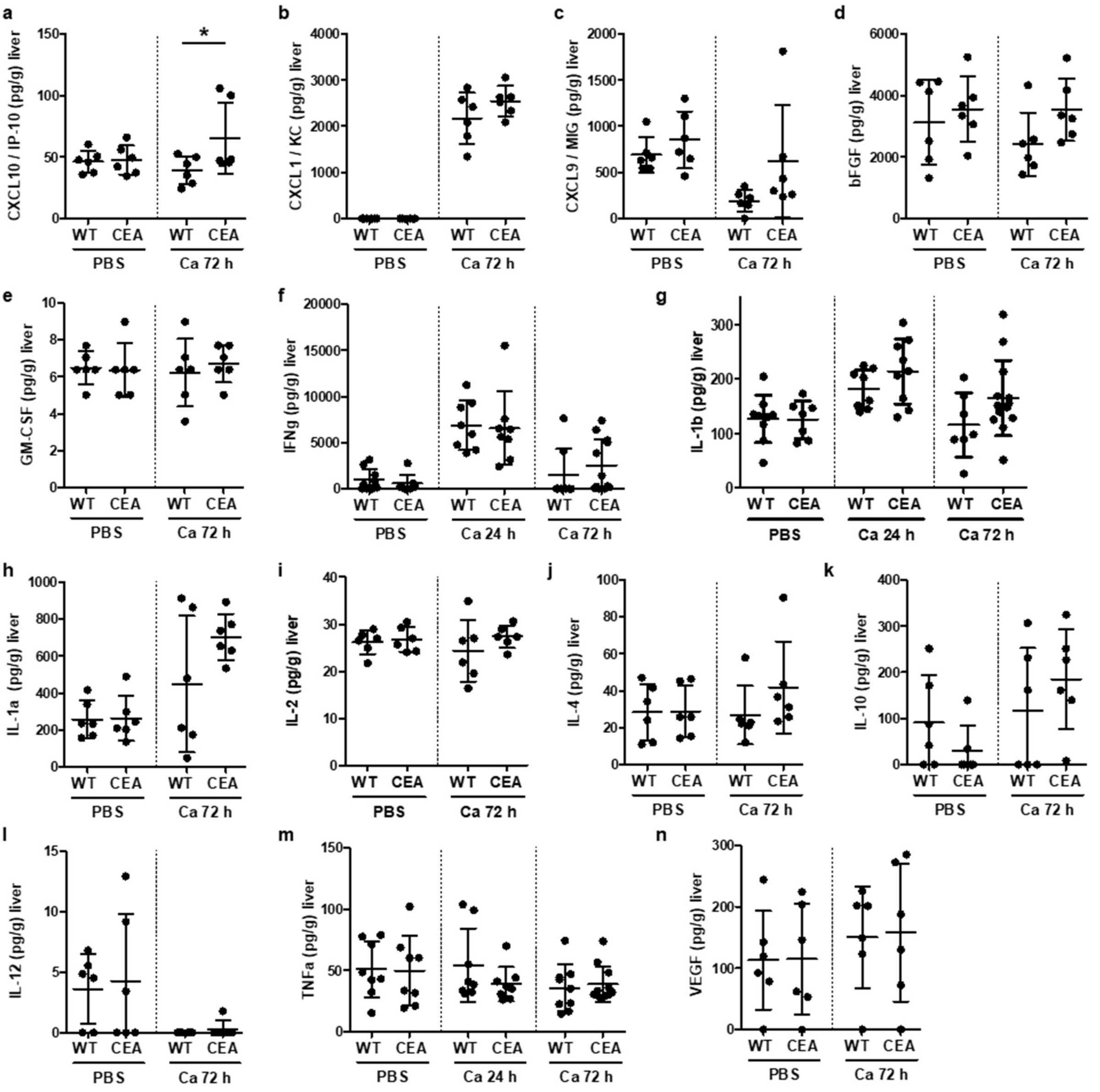


**Figure S9: CEABAC10 mice have enhanced CCL3 and CXCL10 concentrations in their livers during systemic *C. albicans* infection.** CEABAC10 mice (CEA) and their wild type littermates (WT) were either injected with PBS or infected with $1\times10^4$ CFU/g body weight, respectively, and were sacrificed after 24 h or 72 h. Concentrations of cytokines were determined in liver homogenates by multiplex assay (Luminex, 6 per group). For f, g, and m up to 5 additional samples per group were analyzed via ELISA). Data are combined from two independent experiments. Statistics: One-Way-ANOVA and Bonferroni Multiple Comparison Test: *p<0.05 **p<0.01 ***p<0.005, ****p<0.001. Note that IL-5, IL-13, IL-15 and IL-17 were not detected in any liver sample. Graphs give data points with mean and standard deviation. Related to Figure 5.

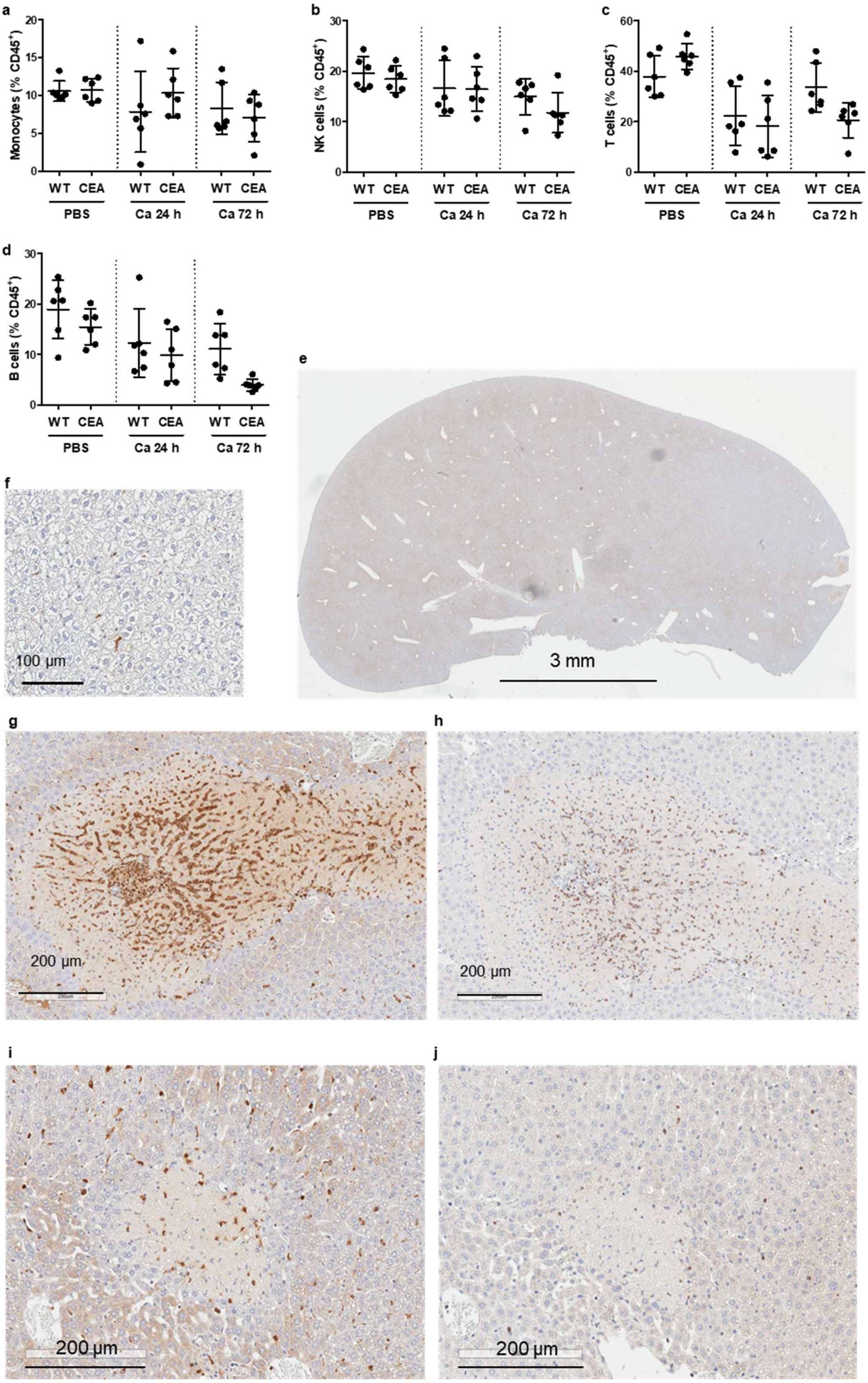
a
Monocytes (% CD45+)
b
NK cells (% CD45+)
c
T cells (% CD45+)
d
B cells (% CD45+)
WT CEA WT CEA WT CEA
PBS
Ca 24 h
Ca 72 h
e
3 mm
f
100 µm
g
200 µm
h
200 µm
i
200 µm
j
200 µm

**Figure S10: Immune cell populations in livers of WT and CEABAC10 mice during systemic *C. albicans* infection.** CEABAC10 mice (CEA) and their wild type littermates (WT) were either injected with PBS or infected with $1\times10^4$CFU/g body weight, respectively, and were sacrificed after 24 h or 72 h. Immune cells were isolated from livers and analyzed via flow cytometry (N=6 per group and staining, combined from two experiments; gating is shown in Figure S11). Monocytes (a), NK cells (b), T lymphocytes (c), and B lymphocytes (d) respectively are given as the percentage of $CD45^+$ cells. Note that data displayed in Figure S10a, b are from the same data sets as Figure 5n, and data in Figure S10c, d are from the same data sets as Figure 5m. Statistics: One-Way-ANOVA and Bonferroni Multiple Comparison Test. Graphs give data points with mean and standard deviation. (e, f) Representative immunohistochemical staining for CEACAM6 of formalin-fixed, paraffin embedded livers of PBS-injected CEA mice. Note that the image in (f) only displays CEACAM6 positive macrophages, but that flow cytometry of immune cells isolated from livers revealed that the majority of the few neutrophils present in a healthy liver were also CEACAM6 positive (Fig. 5). (g-j) Consecutive sections of formalin-fixed, paraffin embedded CEA livers 72 h p.i were used for immunohistochemical staining for CEACAM6 (g, i) or neutrophil elastase (NE, h, j), respectively. Panels display representative images of a necrotic area (N=3, related to Figure 5o-q). Note that only live neutrophils are NE+, but that CEACAM6+ cells include live and dead neutrophils and macrophages/monocytes, and that CEACAM6+ cells (f) outnumber live neutrophils (g) by one or two orders of magnitude. Related to Figure 5.

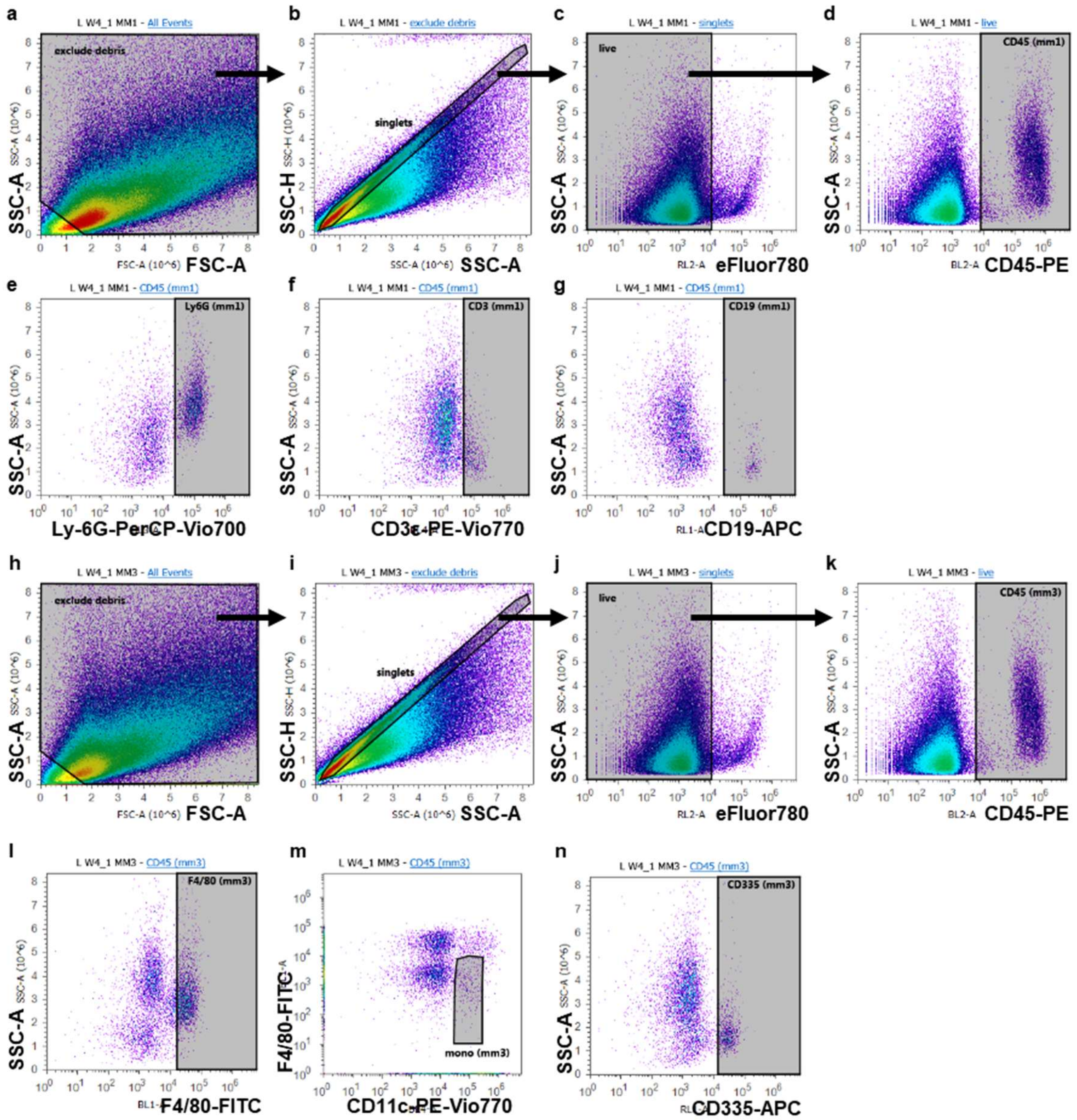


**Figure S11: Gating of immune cell populations in livers.** Representative panels are from a CEABAC10 mouse 72 h p.i., infected with $1\times10^{4}$ CFU/g body weight. Immune cells were isolated from livers and stained for CD45-PE, Ly-6G-PerCP-Vio700, CD3ε-PE-Vio770, CD19-APC, and viability dye eFluor780 (a-g), and F4/80-FITC, CD45-PE, CD11c-PE-Vio770, CD335-APC, viability dye eFluor780 (h-n), respectively, and analyzed by flow cytometry. (a-g) Single (b) / viable (c) / CD45$^{+}$ cells (d) were gated for Ly6G$^{+}$ neutrophils (e), CD3$^{+}$ T lymphocytes (f), and CD19$^{+}$ B lymphocytes (g), respectively. (h-m) Single (i) / viable (j) / CD45$^{+}$ cells (k) were gated for F4/80$^{hi}$ macrophages (l), CD11c$^{int}$/F4/80$^{lo/neg}$ monocytes (m), and CD335$^{+}$ NK cells (n), respectively. Corresponding isotype controls were used to determine positive/negative populations. Cells were analyzed on an on an Attune Acoustic Focusing Cytometer (Life Technologies, Thermo Fisher Scientific) using the Attune software v2.1. Related to Figures 5 and S10.

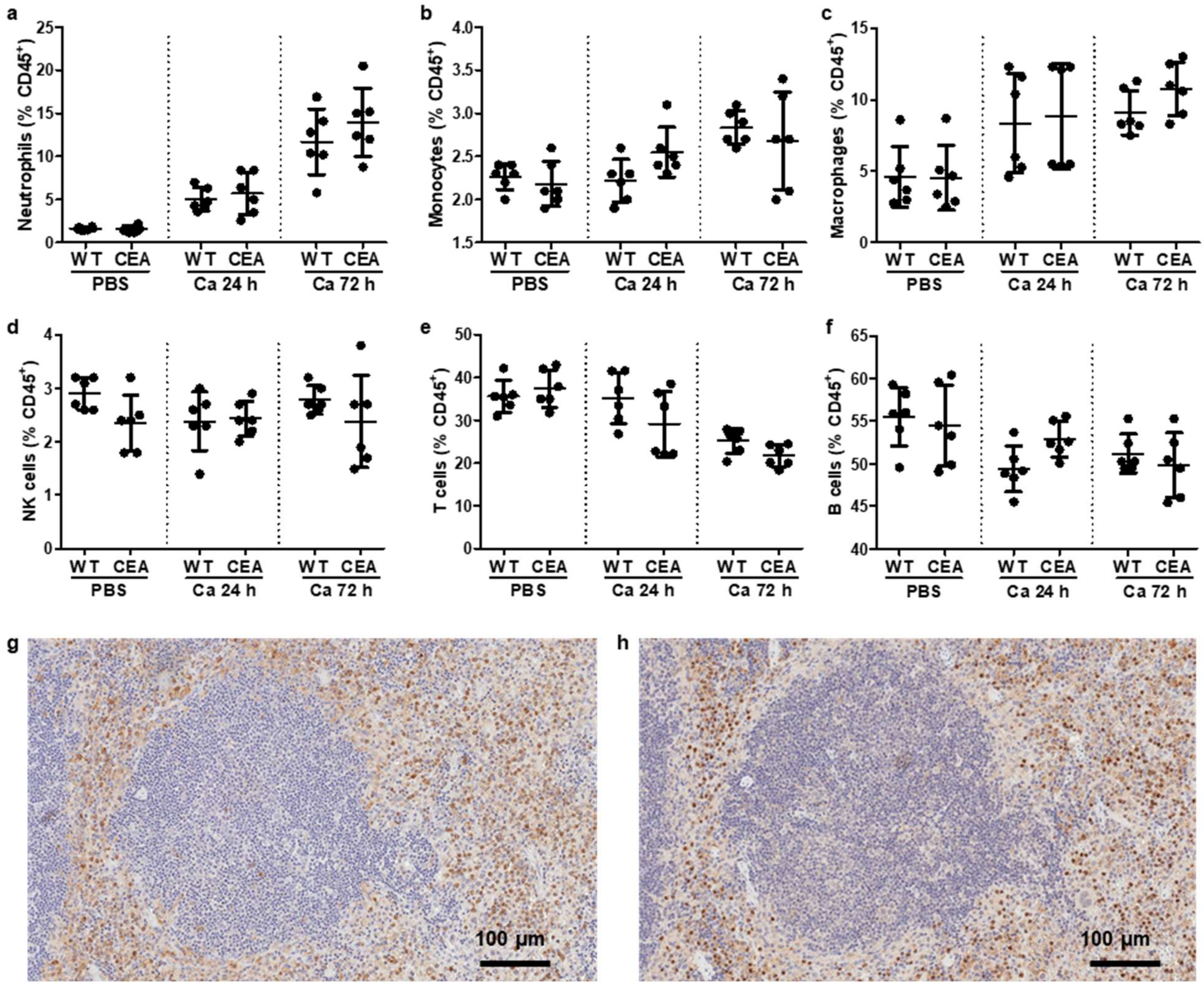


**Figure S12: Immune cell populations in spleen during systemic *C. albicans* infection.** CEABAC10 mice (CEA) and their wild type littermates (WT) were either injected with PBS or infected with 1×10$^4$ CFU/g body weight, respectively, and were sacrificed after 24 h or 72 h. (c-i) Immune cells were isolated from kidneys (N=6 per group and staining), and analyzed by flow cytometry (gating: see Figure S11). Neutrophils (a), monocytes (b), macrophages (c), NK cells (d), T lymphocytes (e), and B lymphocytes (f) are given as the percentage of CD45$^+$ cells with mean and standard deviation. (g, h) Consecutive sections of CEABAC10 spleens 72 h p.i were used for immunohistochemical staining for CEACAM6 (g) or neutrophil elastase (h), respectively. Panels display representative images (N=3). Note that only live neutrophils are NE$^+$, but that CEACAM6$^+$ cells include live and dead neutrophils. In contrast to kidneys (Fig.3q, r), livers (Fig. 5p, q) and brain (Fig. S10), similar numbers of cells are positive for CEACAM6 and neutrophil elastase. Data are combined from two independent experiments. Statistics: (a-f) One-Way-ANOVA and Bonferroni Multiple Comparison Test. Related to Figure 5.

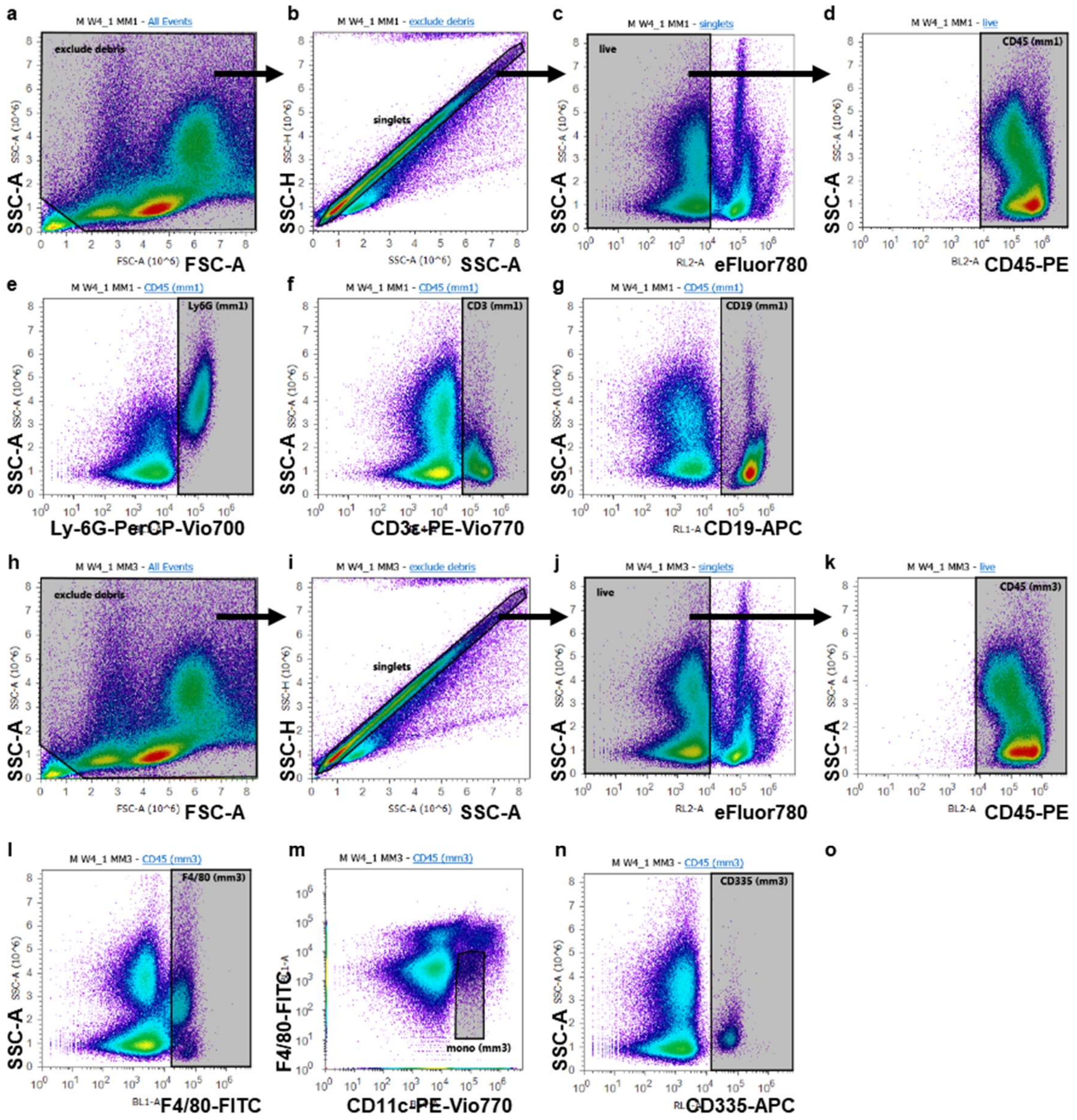


**Figure S13: Gating of immune cell populations in spleens.** Representative panels are from a CEABAC10 mouse 72 h p.i., infected with $1\times10^4$ CFU/g body weight. Immune cells were isolated from spleens and stained for CD45-PE, Ly-6G-PerCP-Vio700, CD3ε-PE-Vio770, CD19-APC, and viability dye eFluor780 (a-g), and F4/80-FITC, CD45-PE, CD11c-PE-Vio770, CD335-APC, viability dye eFluor780 (h-n), respectively, and analyzed by flow cytometry. (a-g) Single (b) / viable (c) / $CD45^+$ cells (d) were gated for $Ly6G^+$ granulocytes (e), $CD3^+$ T lymphocytes (f), or $CD19^+$ B lymphocytes (g), respectively. (h-m) Single (i) / viable (j) / $CD45^+$ cells (k) were gated for $F4/80^{hi}$ macrophages (l), $CD11c^{int}/F4/80^{lo/neg}$ monocytes (m), and $CD335^+$ NK cells (n), respectively. Corresponding isotype controls were used to determine positive/negative populations. Cells were analyzed on an on an Attune Acoustic Focusing Cytometer (Life Technologies, Thermo Fisher Scientific) using the Attune software v2.1. Related to Figure S12.

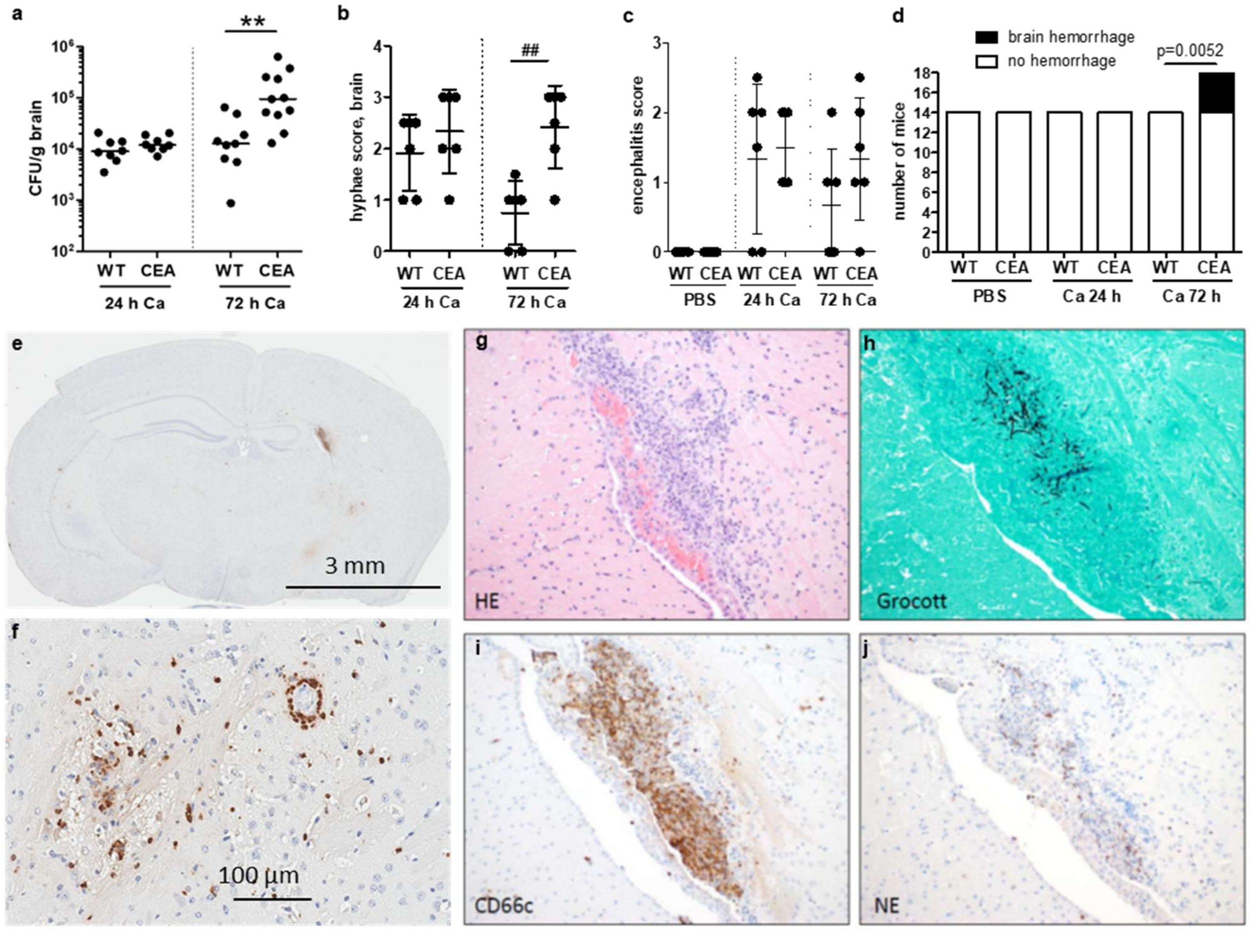


**Figure S14: CEABAC10 mice display brain hemorrhage but no increase in encephalitis 72 h post-*C. albicans* infection.** CEABAC10 mice (CEA) and their wild type littermates (WT) were either injected with PBS or infected with $1\times10^4$ CFU/g body weight, respectively, and were sacrificed after 24 h or 72 h. (a) CFUs were determined in brain homogenates (8 WT and 8 CEABAC10 for 24 h Ca, 9 WT and 11 CEABAC10 for 72 h Ca). (b) Grocott-silver-stained brain sections were scored for the occurrence of hyphal growth (6 per group). Note that brains from uninfected animals (PBS) did not display any fungal growth (a: 8 WT and 8 CEA; b: 6 WT and 6 CEA). (c) HE-stained brain sections were scored for the degree of encephalitis (N=6 per group). (d) Occurrence of macroscopic brain hemorrhage observed during necropsy. Data are combined from two (a-c) or four (d) independent experiments. Statistics: (a) One-Way-ANOVA and Bonferroni Multiple Comparison Test: **$p<0.01$; (b, c) Kruskal-Wallis/Dunn's Multiple Comparison Test, $^{\#\#}p<0.01$; (d) Chi square test for trend. (a-c) Graphs give data points with mean and standard deviation. (e, f) Panels display representative CEACAM6 immunohistochemistry staining on sections of three CEABAC10 brain sections 72 h p.i. from one experiment. Note that only neutrophils and macrophages are CEACAM6-positive. (g-i) Representative panels of consecutive sections from a brain of a CEABAC10 mouse infected with $1\text{x}10^5$ CFU/g body weight and sacrificed after 72 h. (g) Hematoxylin-eosin stain (HE). (h) Grocott-silver stain. (i) Immunohistochemistry for CEACAM6 (CD66c). (j) Immunohistochemistry for neutrophil elastase (NE). Note that only live neutrophils are $NE^+$, but that $CEACAM6^+$ cells include live and dead neutrophils and macrophages, and that $CEACAM6^+$ cells (i) outnumber live neutrophils (j) by ca. one or two orders of magnitude.

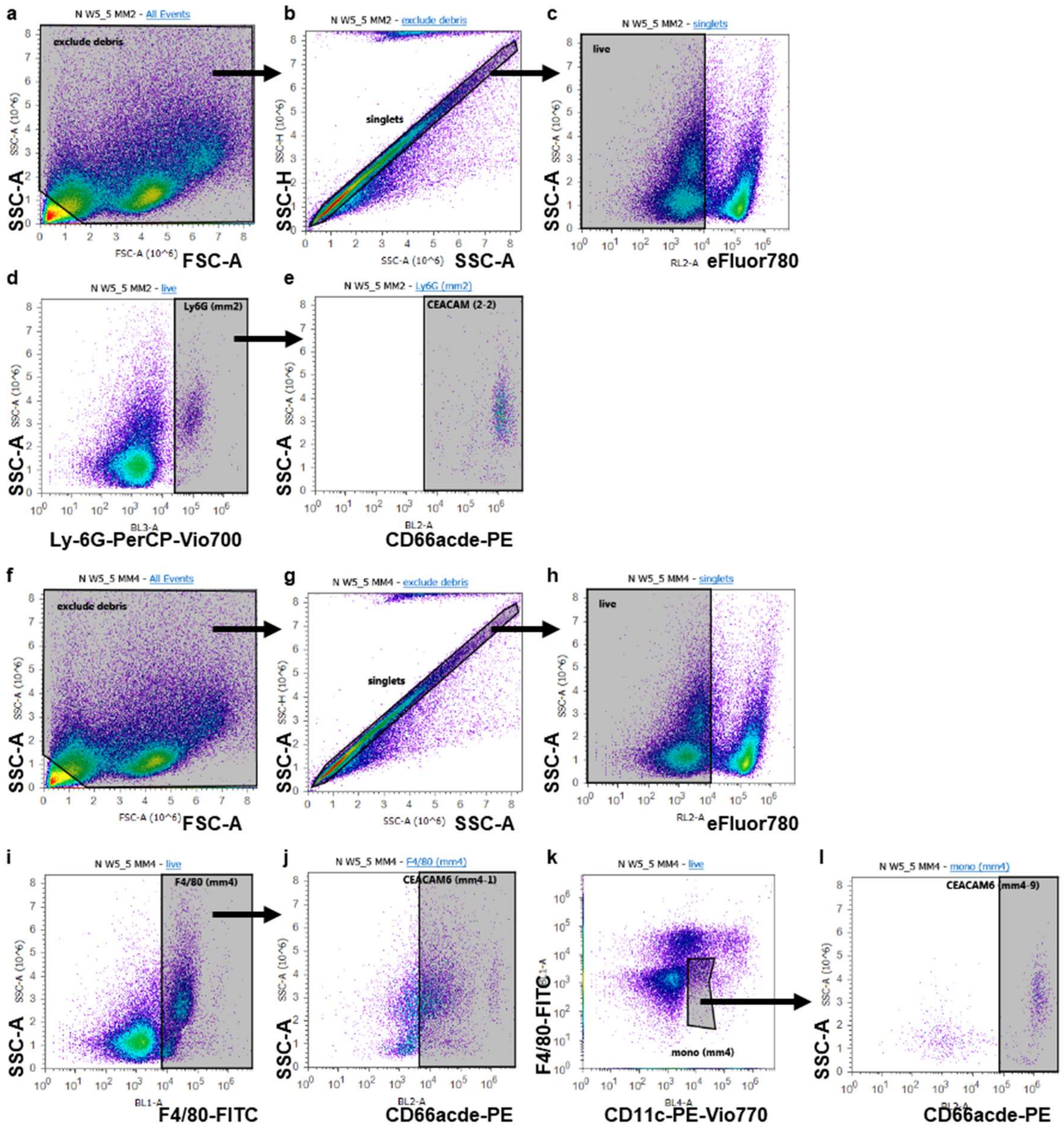


**Figure S15: Gating of CEACAM6$^+$ myeloid cells in organs of CEABAC10 mice during systemic *C. albicans* infection.** Panels display gating of CEACAM6+ immune cells isolated from CEABAC10 kidneys, liver, and spleen (related to Figure 6). Immune cells were isolated from kidneys, liver, and spleen, and stained for viability dye eFluor780/Ly6G-PerCP-Vio700/CEACAM3/6 (CD66acde-PE), and viability dye eFluor780/F4/80-FITC/ CD11c-PE-Vio770/CEACAM3/6 (CD66acde-PE), respectively. Representative panels are from a kidney 72 p.i. Single (b, g), viable (c, h) cells were gated for Ly6G$^+$ neutrophils (d), F4/80$^{hi}$ macrophages (i), and CD11c$^{int}$/F4/80$^{lo/neg}$ monocytes (k), respectively, and then analyzed for their percentage of human CEACAM6-positive cells (e, j, l). Corresponding isotype controls were used to determine positive/negative populations. Cells were analyzed on an on an Attune Acoustic Focusing Cytometer (Life Technologies, Thermo Fisher Scientific) using the Attune software v2.1. Related to Figure 6.

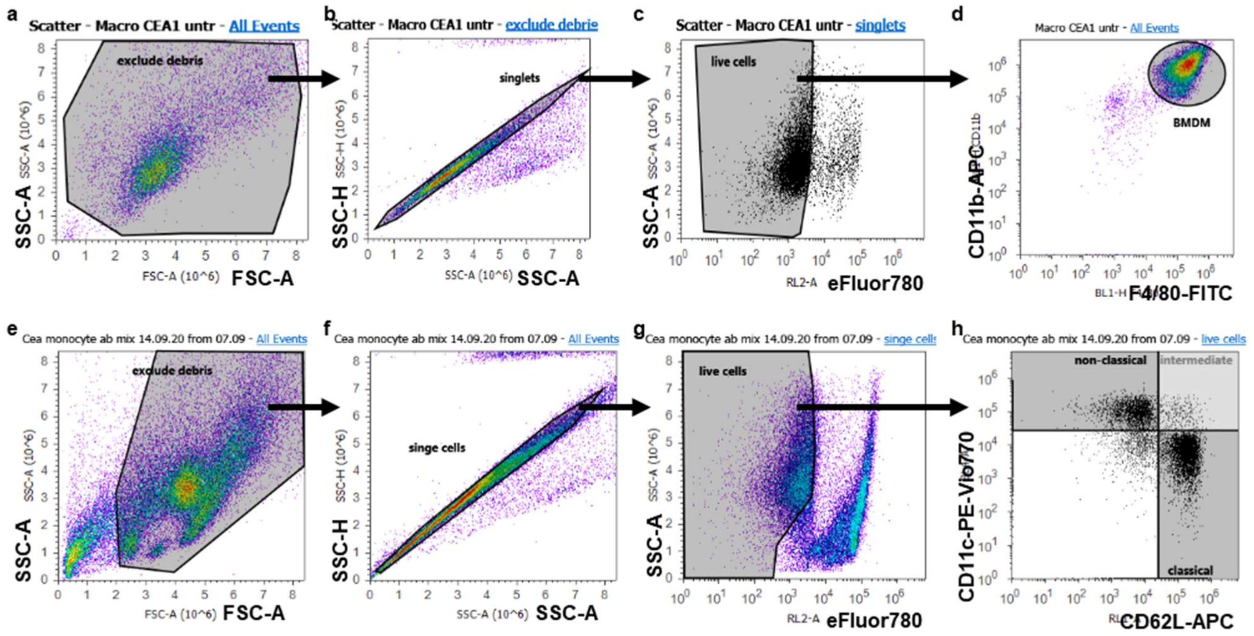


**Figure S16: Gating of bone marrow-derived macrophages and monocytes**. (a-d) Bone marrow-derived macrophages (BMDM) from CEABAC10 mice were obtained after 7 days of differentiation with 25 ng/ml rmM-CSF. Adherent cells were collected and stained for viability dye eFluor780, F4/80-FITC, CD11b-APC, and CEACAM6 (CD66c-PE), and analyzed via flow cytometry. Single (b) viable (c) cells were gated for F4/80$^+$/CD11b$^+$ macrophages (d) and analyzed for their CEACAM6 expression (shown in Figure 7b). (e-h) Bone marrow-derived monocytes (BMM) from CEABAC10 mice were obtained after 5 days of differentiation with 25 ng/ml rmM-CSF. Non-adherent cells were collected and stained for viability dye, CD11c-PE-Vio770, CD62L-APC, and CEACAM6 (CD66c-PE), and analyzed via flow cytometry. Single (f) viable (g) cells were gated for CD11c$^+$/CD62L$^-$ non-classical monocytes, CD11c$^+$/CD62L$^+$ intermediate monocytes, and CD11c$^-$/CD62L$^+$ classical monocytes (h) and analyzed for their CEACAM6 expression, respectively (shown in Figure 8b-d). Graphs are representative of three independent experiments. Corresponding isotype controls were used to determine positive/negative populations. Cells were analyzed on an on an Attune Acoustic Focusing Cytometer (Life Technologies, Thermo Fisher Scientific) using the Attune software v2.1. Related to Figures 7 and 8.

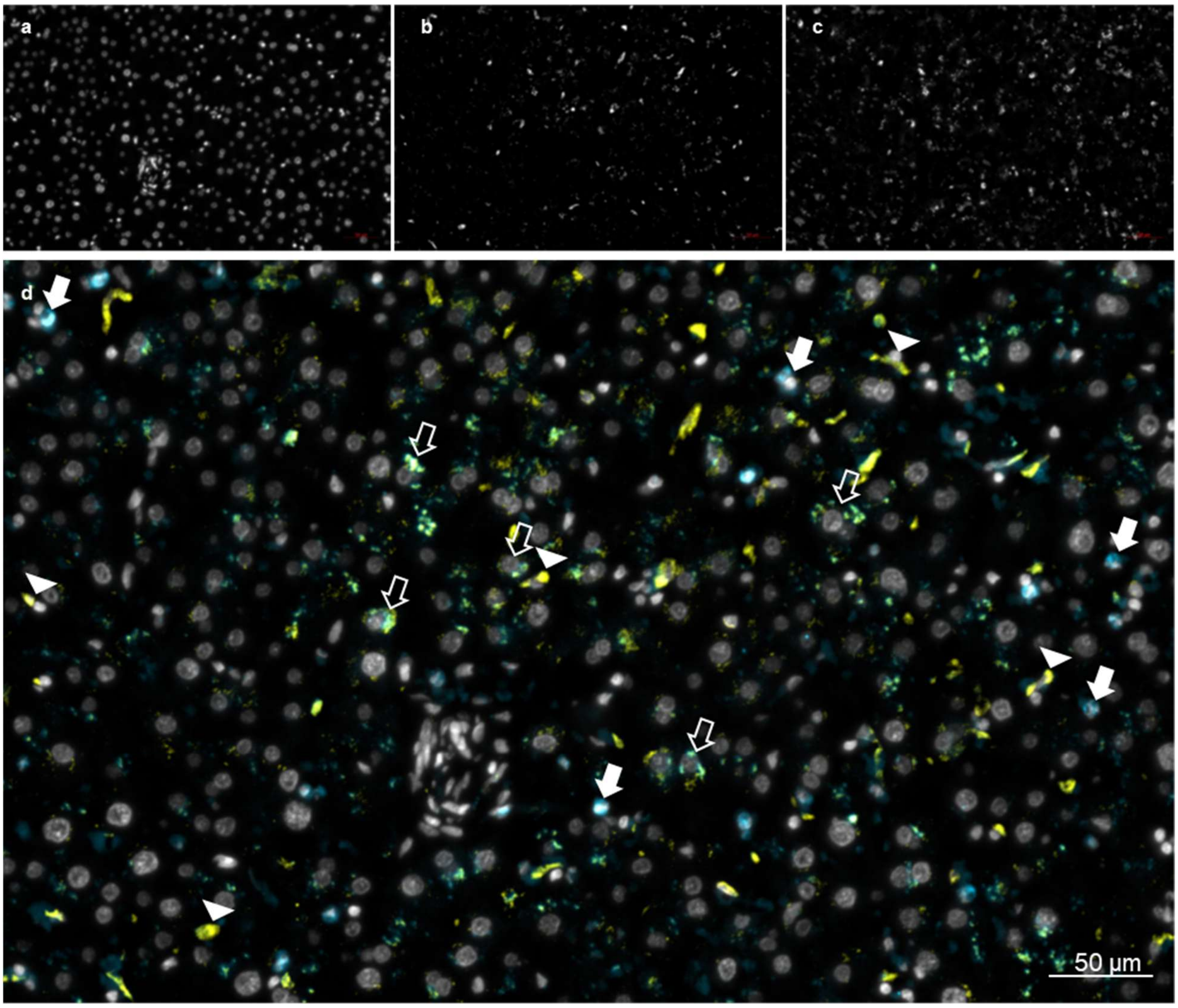


**Figure S17: CEACAM6 expression on human liver macrophages.** Human liver sections were stained for nuclei (Hoechst 33342) (a), the marker of resident liver macrophages CD68/GaR-Alexa660 (b), and CEACAM6 (1H7-4B)/GaM-Alexa 546 (c). Stained sections were analyzed by confocal laser scanning microscopy. (d) displays an overlay of (a-c) with yellow coloring for CD68, blue coloring for CEACAM6, and no coloring (white) for nuclei. Five examples of resident macrophages expressing both, CD68 and CEACAM6 (open arrows), of resident macrophages expressing CD68 but no CEACAM6 (filled arrowheads), and of polymorphonuclear neutrophils expressing CEACAM6 but no CD68 (filled arrows), are indicated, respectively. Panels are representative of sections from two livers, though only one liver (shown above) displayed neutrophil infiltrations.

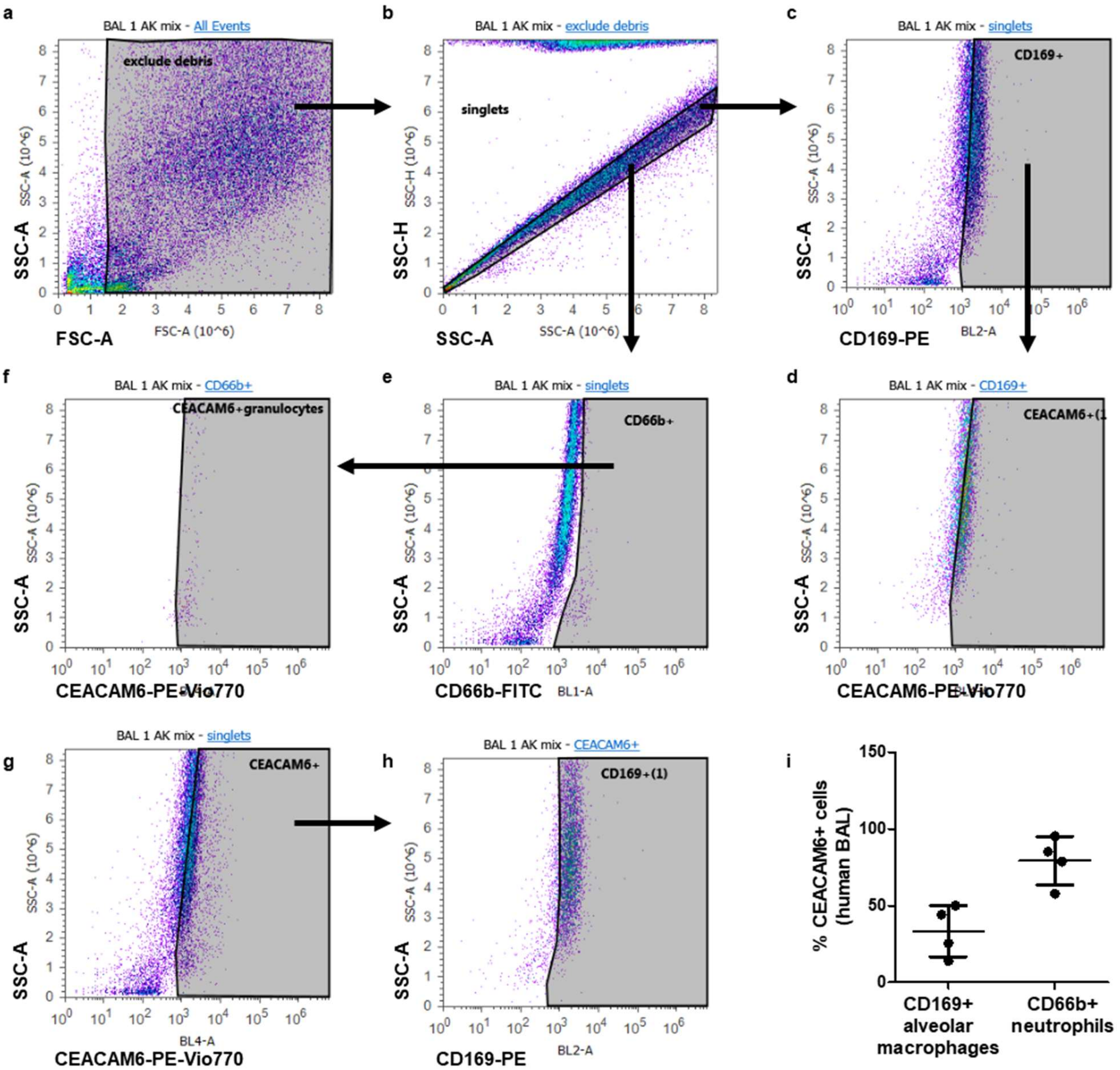


**Figure S18: CEACAM6 expression on human alveolar macrophages**. (a) Cells from human bronchoalveolar lavage were stained for CD169-PE (marker for alveolar macrophages), CD66b-FITC (marker for human neutrophils), and CEACAM6 (CD66c-PE-Vio770). Cells were analyzed on an Attune Acoustic Focusing Cytometer (Life Technologies, Thermo Fisher Scientific) using the Attune software v2.1. Single cells (b) were gated for $CD169^+$ alveolar macrophages (c) or $CD66b^+$ neutrophils (e) and analyzed for their CEACAM6 expression (d, f, respectively). Single cells were also gated for $CEACAM6^+$ cells (g) and analyzed for their CD169 expression (h). Note that the majority of $CEACAM6^+$ cells are also $CD169^+$ alveolar macrophages. Panels are representative of four different bronchoalveolar lavages analyzed in two independent experiments. The graph displays the percentage of $CEACAM6^+$ alveolar macrophages (33.5% ±16.6%) and of $CEACAM6^+$ neutrophils (79.3% ±15.8%) with mean and standard deviation.

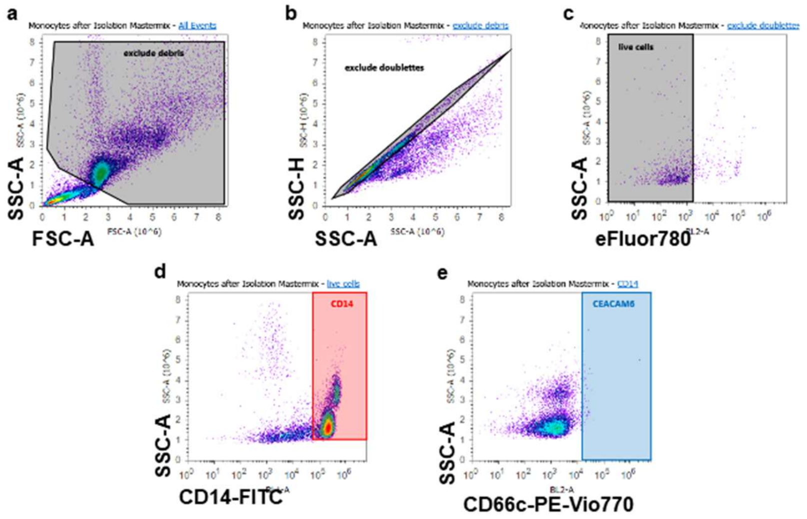


**Figure S19: No CEACAM6 expression on human peripheral monocytes**. Human PBMCs were isolated and stained for CD14-FITC, CD66c-PE-Vio770 (CEACAM6), and eFluor780. Cells were analyzed on an on an Attune Acoustic Focusing Cytometer (Life Technologies, Thermo Fisher Scientific) using the Attune software v2.1. Single cells (b) were gated for live (c), $CD14^{+}$ cells (d), and analyzed for their CEACAM6 expression (e). Panels are representative of three independent experiments.

### 2. Supplemental Method

#### Automated image processing of *Candida albicans* in mouse kidneys

In order to ensure a precise and objective quantification of *Candida albicans* in histopathological kidney sections of mice compared to a subjective evaluation by a pathologist, an automated and objective image analysis is required, which is validated to achieve consistently accurate results under various demanding image modalities. To address this, we have developed a computer-aided pipeline to (i) segment fungi in histopathological stained mouse tissue sections and (ii) distinguish the detected fungus according to its morphotype as a yeast cell or in its filamentous form, which we refer here as hyphae.

**Segmentation of kidney tissue.** To determine the size of the histologically stained tissue sections, the mouse kidneys had to be segmented as a whole. This was realized using the visual programming language JIPipe[67] which is based on the image processing tool ImageJ[74]. JIPipe takes image analysis to a new, user-friendly level by functioning as a visual programming language that allows users to create flowcharts instead of programming concatenated image analysis operations. At the same time, the FAIR[75] principles are ensured by JIPipe. To obtain a binary image that distinguishes between the kidney in the foreground and all the rest as background, Otsu[76] thresholding was applied to the stained images, where the threshold was determined by minimizing the intra-class intensity variance per tissue section. To close perforated cells of the kidney section after binarization, we applied morphological closing with a disk shape and kernel size seven pixel. Besides joining perforated regions of interest (ROI), it was necessary to remove remaining artifacts, for which we used the Remove Outliers 2D function of ImageJ, which was designed for dead pixel correction. This operation works by replacing a pixel with the median of the surrounding pixels if the intensity value deviates from the median by more than a certain threshold. The number of surrounding pixels used for the calculation of the median is determined by the size of the previously chosen kernel size with 26 pixels. On visual inspection, we found that some areas in the tissue of the kidney are very susceptible to errors in the staining process. This involves the appearance of small indentations that resemble cracks, which would noticeably reduce the total area of the underlying tissue per section. To reduce these errors of the segmentation in retrospect, we examined the individual size of the cracks within a histogram and determined a threshold of $1,5 \times 10^5$ pixels (i.e. 37995 $\mu m^2$) to fill these holes and include them in the foreground. For each sample an image was created for visual control in which the outlines of the segmented kidney were drawn in the originally stained image and the final segmentation result was assessed by experimental experts. Subsequently, the total area of each mouse kidney was quantified.

**Data preparation for the quantification of fungal burden.** The quantification of fungal burden within a histopathological tissue section is usually performed by a pathologist through visual inspection resulting in an ordinally scaled score per slice. To perform the analysis as objective as possible, especially to avoid subjective bias, we conducted the image analysis in an automated fashion. Although standardized histopathological silver staining was used, which is particularly suitable for objects with a high sugar content such as fungi, the outcome of the staining process itself is highly variable. Sample handling, e. g. differences in the length of fixation and the time span between fixation and embedding, as well as the timing within the staining process itself can easily lead to a variety of hues and intensity saturation. Consequently, it was not feasible to apply a single model approach to all samples. Despite this, in order to be able to validate the automated image analysis, eleven representative cutouts were selected from a total of eight different samples that visually cover the entire bandwidth of different hues and intensity saturation of fungi in the tissue. The fungus was subsequently manually annotated within the pruned cutout regions by experimental experts, where the manual annotations differentiated between (i) yeast cells and (ii) hyphae. The latter also included overlying fungus, which may also contain unidentifiable yeast cells.

**Proof of concept with deep learning neural networks**. Upon visual inspection of the histologically stained kidney tissue, it was readily apparent that the hue of the fungus-infected tissue resulted in either black or dark red colors after the staining process. Due to this, it was not possible to achieve satisfactory results by color thresholding, making structure and shape-based methods necessary for a segmentation independent of staining. Here we used SegNet[77], a well-established deep neural network capable of performing semantic segmentation, which is based on the encoder-decoder U-Net[78] architecture. This model was applied as part of two leave-one-out cross-validations (LOOCV) [78] per sample, one for each staining setting (5-k-fold and 6-k-fold). For the training, a total of 437 patches were available, each with a size of 512 x 512 pixels, derived from the eleven cutouts. The training for each model was done with a maximum of 300 epochs and a batch size of 32 using standard data augmentation techniques such as rotation, flipping and zooming, as well as elastic transformation[79], which introduces additional shape-variability to the image content. To prevent overfitting, we applied Gaussian Dropout as a regularization method on the intermediate convolutional layers. Since the model output is a probability value per pixel, with the value of one and maximum probability to be identified as fungus or zero with minimum probability to be assigned fungus, we binarized each image using the Isodata[80] algorithm. Filtering out possible artifacts that are smaller than the smallest manually annotated yeast cell, we removed all ROIs smaller than 29 pixels in area, which corresponds to 7.35 µm². Assuming a circular shape of yeast cells, the accepted minimal diameter for yeast cells must be at least 3 µm. A final post-processing was done by filtering stained objects that are at least $10^4$ pixels (2533 µm²) in size and have a donut shape (glycosylated tissue structures), while the outer shell has the typical diameter and color of hyphae, resulting in an increased false positive error and falsely magnifying the fungus fraction. Thus, artefacts derived from the sectioning process and tissue artefacts possessing highly glycosylated surfaces were excluded efficiently. To achieve this, we filtered out all objects larger than $10^4$ pixels with a solidity of less than 0.3, which is defined by the surface area divided by the area of the object's convex hull. The threshold 0.3 was chosen, because the solidity of such artifacts was typically observed to be below 0.2. We then evaluated each test cutout per validation step, with two cutouts per sample present in three k-folds. The final quantitative evaluation is based on the (i) Dice coefficient, which is mathematically identical to the F1 measure and the two scores for (ii) Precision and (iii) Recall, which, respectively, take into account the error of the false positives (FP) as well as the false negatives (FN). All measures utilized here consider (non-)overlap between two sets, where the two sets consist of the manual annotations and the model predictions. The complete overlap between both sets is defined as true positive (TP), while the over-detection of the predictions which are not present in the manual annotations is defined as FP. In the opposite case, all non-detected values of the manual annotations are defined as FN. The Dice coefficient is defined as twice the TP divided by twice the TP plus FP and FN. The Precision metric is defined by TP divided by TP plus FP, while in Recall the FP parameter is replaced by the other type of error (FN). After the proof of concept that SegNet works on the underlying data, we follow best practice using cross-validation evaluation in common machine learning settings, having found the best parameters for retraining the model by taking all training data into account. For this purpose, we trained two models from scratch, one per staining with five cutouts in which the fungus appears black and six cutouts with fungus appearing dark red. These two models were then used to predict whole samples and quantify the fungi within the kidneys.

**Differentiation of yeast and hyphae.** The fungus segmentation was then used to distinguish between yeast cells and hyphae, where the most obvious feature is their difference in size. Considering and analyzing the manual annotations, we classified yeast cells based on size in a range of 74 to 140 pixels (19 - 35 µm²). Any object larger than 140 pixels (35 µm²) were assigned to be hyphae. Correctly identified yeast cells and hyphae were scored by comparing the covered area and the total amount of counted objects, as well as the relative fraction per morphotype within the fungus. This approach was then applied to all samples to finally quantify yeast cells and hyphae.

**Statistical analysis.** Statistical tests for the fungal fraction were performed using the unpaired Wilcoxon rank sum test and using the R package "effectsize" to calculate the Hedge 'g effect size[10]. The ranges of Hedges 'g effect size magnitudes are referred to as being negligible for $|g| < 0.2$, small for $|g| < 0.5$, medium for $|g| < 0.8$ and large for $|g| \geq 0.8$. Furthermore, to facilitate a comparison between the manual annotation and the model prediction in terms of the fraction of fungus, the objects counted, and the area covered by *Candida albicans*, we calculated the Pearson's product-moment correlation coefficient[11]. This test not only provides the correlation coefficient, but also tests the correlation between paired samples and provides a p-value if the correlation is significantly different from zero across all samples, meaning that the two variables are significantly correlated with each other.

**Data and code availability.** All material, consisting of the JIPipe files and Python scripts are available for download at: https://asbdata.hki-jena.de/Klaile_PraetoriusEtAl2023_NatCommun.

**Evaluation of *C. albicans* segmentation in histological tissue sections.** Within the LOOCV, we obtained a median Dice coefficient of 64% with the SegNet model, which acts as a heuristic mean between Precision (median of 88%) and Recall (median of 57%). The model generally misses rather than over-segments the fungus. Considering the retrained model for each of the two staining procedures, we obtained a median of 76% for the Dice coefficient, 83% for Precision, and 79% for Recall.

**Evaluation of the differentiation between the morphotypes.** After evaluating the fraction of fungus per cutout, we determined measures such as the relative amount of counted objects from yeast and hyphae within the fungus for both, the manual annotations, and the predictions within the LOOCV and for the retrained model. The Pearson correlation coefficient within the LOOCV evaluation for yeast and hyphae in relation of fractional counted objects is 0.42, whereas the same correlation coefficient for the absolute number of counted objects of yeast cells is 0.78 and for hyphae is 0.96. For both yeast cells (p-value $< 0.004$) and hyphae (p-value $\ll 10^{-5}$), the Pearson's product-moment correlation coefficient is significantly different from zero, indicating a strong correlation between manual annotation and prediction. In relation to the measured value of the covered area of the yeast cells, there is a correlation coefficient of 0.83 between the manual annotations and the prediction, which is significantly different from zero (p-value $< 0.0016$), and a coefficient of 0.97 for the yeast cells, where there is also a considerably significant difference (p-value $\ll 10^{-6}$). Considering the retrained model, the correlation coefficient of the fractional yeast/hyphae on the whole fungus per cutout is 0.36, which is not significantly different from zero between the manual annotations and the model prediction. In contrast, the measurement of the counted objects for both morphotypes, yeast cells (correlation = 0.85, p-value $< 0.0008$) and hyphae (correlation = 0.96, p-value $\ll 10^{-5}$), correlates significantly. The comparison between the ground truth and the prediction concerning the area covered by yeast cells (correlation = 0.89, p-value $< 0.0002$) and hyphae (correlation = 0.99, p-value $\ll 10^{-10}$) is of similar extent.

**Fungus fraction in mouse kidneys.** After segmenting the kidney sections and the fungus contained within, i.e. the area covered by fungus, was summarized and the relative fraction of fungus per kidney slice was determined. The results of the final quantification and statistical analysis are shown in Fig. S7f, where the fungal fraction per kidney slice is shown grouped across the corresponding genotype (CEABAC10 and wildtype (WT)). While the CEABAC10 mice exhibit significantly higher fungal burden over both time points combined as compared to the WT (p-value $< 0.027$, effect size = 0.75), significantly more fungal burden is present within the CEABAC10 genotype group after 72 hours compared to 24 hours (p-value $< 0.002331$, effect size = 1.22). To the same extent, significantly more fungus is present in the CEABAC10 mouse kidneys than in the wildtype at 72 hours (p-value $< 0.002331$, effect size = 1.22). While the values for the WT mice range from 0.002% to 0.03% at 24 hours (0.007% to 0.1% at 72 hours), the fungal fraction for the CEABAC10 genotype at 24 hours is between 0.003% and 0.07% (0.0005% to 8% after 72 hours).

**Yeast and hyphae fraction within the fungus.** With respect to both measured values, the yeast and the hyphae fraction of the total fungus per slice, as well as their absolute number of objects, we could not observe any significant differences. However, regarding the covered area for both morphotypes in Fig. S7g, it was found that after 72 hours, CEABAC10 mice had significantly higher covered area for yeast cells (p-value < 0.0023, effect size = 1.07) and hyphae (p-value < 0.0041, effect size = 1.03) compared to the wildtype.

**Cross-validation of automated image processing for fungal segmentation.** Although the median of the Dice coefficient appears to be low with a median of 64%, it should be pointed out that segmenting the fungus in both stainings is an extremely challenging task. While we performed the same LOOCV procedure with a color-based thresholding method (median of 55% w.r.t Dice coefficient) and an interactive machine learning method called ilastik[12] (median of 43% w.r.t Dice coefficient). The latter is particularly capable of using structure- and texture-based features for semantic segmentation in biomedical images, however, the Precision was not acceptable due to many false-positives. The Precision corresponds to a median of 39% in the case for the color-based thresholding and 53% with ilastik, while it is 88% with our SegNet approach. For the subsequent quantification of fungus within the corresponding kidney, this would have likely resulted in considerable over-detection of fungus using both the color thresholding method and ilastik. Therefore, we decided to use the deep learning method SegNet, which was able to learn the features data-driven from the relatively few data points for typical sizes of training data of deep learning methods using only 437 patches.

Regarding the Dice coefficient metric, it is worth mentioning that it does not reflect small ROIs, such as yeast cells, with the same attention as larger objects due to the calculation of pixel-wise overlaps. As a consequence, in an image with a few objects in total, small errors in the segmentation can lead to large quantitative errors. Although the Dice coefficient is frequently used in image analysis and is now used by default for segmentation comparisons alongside the Jaccard index, which is mathematically correlated with the Dice coefficient, its weaknesses and shortcomings are presented in a continuously updated article[13].

**Performance evaluation between yeast cells and hyphae differentiation.** The distinction between the two morphotypes required other metrics compared to segmentation, the most obvious ones here being in the relative fraction of both morphotypes, the objects counted, and their covered area. Although there is a slight under-detection in the trend for both yeast cells and hyphae, looking at the latter two measures and their significant correlation coefficients, it can be concluded that both morphotypes are positively correlated with each other to the same extent between the manual annotation and the prediction. This consistency in the range of values between the relative fraction of yeast and hyphae is equally important and is present, as can be seen in Fig. 4, because misidentification of one morphotype will result in a leverage effect and negatively influence the other morphotype to the same extent.

**Final quantification of the fungal fraction and the respective morphotypes.** Using validated computer-aided and modern deep learning-based methods, the fungal fraction could be determined effectively within the mouse kidney sections, resulting in robust data despite the high variance of the images. The most significant difference appeared between the CEABAC10 mice in which after 72 hours noticeably more *Candida albicans* was detected in the kidneys than in the wildtype littermate control group. When yeast cells and their germinated morphotype, which we generally refer to here as hyphae, were distinguished in the next step, it was found that the relative fraction of *Candida albicans* yeast cells and hyphae was present in the same order of magnitude at both examination time points. The investigation of the absolute numbers with respect to the identified morphotypes and their respective covered surface area per mouse kidney, again shows a significantly higher yeast and hyphae burden in the CEABAC10 mice compared to the WT after 72 hours. Relative and absolute measurements taken together thus reveal that the generally higher fungal burden after 72 hours can be explained by the equally increased presence of yeast cells and hyphae.


73. Wasserman S, Hedges, L. V. & Olkin, I. Statistical Methods for Meta-Analysis *J Educ Stat* **13**, 75 (1988).

67. Gerst R, Cseresnyes Z, Figge MT. JIPipe: visual batch processing for ImageJ. *Nat Methods* **20**, 168-169 (2023).

74. Rueden CT*, et al.* ImageJ2: ImageJ for the next generation of scientific image data. *BMC Bioinformatics* **18**, 529 (2017).

75. Wilkinson MD*, et al.* The FAIR Guiding Principles for scientific data management and stewardship. *Sci Data* **3**, 160018 (2016).

76. N O. A Threshold Selection Method from Gray-Level Histograms. *IEEE Transactions on Systems, Man, and Cybernetics* **9**, 62-66 (1979).

77. V. Badrinarayanan AKRC. SegNet: A Deep Convolutional Encoder-Decoder Architecture for Image Segmentation. *IEEE Transactions on Pattern Analysis and Machine Intelligence* **39**, 2481-2495 (2017).

78. Olaf Ronneberger PF, Thomas Brox. U-Net: Convolutional Networks for Biomedical Image Segmentation. *arXiv* (2015).

79. Facon HAL-AaJ. Image segmentation by learning approach. In: *Seventh International Conference on Document Analysis and Recognition*) (2003).

80. Calvard TWRaS. Picture Thresholding Using an Iterative Selection Method,. *IEEE Transactions on Systems, Man, and Cybernetics* **8**, 630-632 (1978).
